# Postembryonic developmental roles of the Arabidopsis *KEULE* gene

**DOI:** 10.1101/2024.03.14.585090

**Authors:** Alejandro Ruiz-Bayón, Carolina Cara-Rodríguez, Raquel Sarmiento-Mañús, Rafael Muñoz-Viana, Francisca M. Lozano, María Rosa Ponce, José Luis Micol

**Author notes:** Corresponding author: J.L. Micol. Present address: Fundación para la Investigación Biomédica del Hospital Universitario Puerta de Hierro, 28222 Majadahonda, Madrid, Spain (Rafael Muñoz-Viana). Present address: Instituto Bernabeu, 03016 Alicante, Spain (Francisca M. Lozano).

## Abstract

Cytokinesis in plant cells begins with the fusion of vesicles that transport cell wall materials at the center of the cell division plane, where the cell plate forms and expands radially until it fuses with the parental cell wall at the preprophase band. Vesicle fusion is facilitated by *trans*-SNARE complexes, with assistance from Sec1/Munc18 (SM) proteins. The SNARE protein KNOLLE and the SM protein KEULE are required for membrane fusion at the cell plate. Due to the crucial function of KEULE, all Arabidopsis (*Arabidopsis thaliana*) *keule* mutants identified to date are seedling lethal. Here, we identified the Arabidopsis *serrata4-1* (*sea4-1*) and *sea4-2* mutants, which carry recessive, hypomorphic alleles of *KEULE*. Homozygous *sea4-1* and *sea4-2* plants are viable and fertile but exhibit smaller rosettes and fewer leaves at bolting than the wild type. Their leaves are serrated, small, and undulated, with a complex venation pattern, develop necrotic patches, and undergo premature senescence. We established a likely relationship between these phenotypes and their defects in cytokinesis through reduced cell wall integrity and increased unfolded protein response. These findings shed light on the roles of KEULE in postembryonic development, particularly in the patterning of rosette leaves and leaf margins.

## 1. INTRODUCTION

During cytokinesis, the cytoplasm of a cell is divided, resulting in the formation of two daughter cells. In plant cells, the plane of cell division aligns with the phragmoplast, a dynamic matrix of antiparallel microtubules that transports vesicles containing cell wall materials from the trans-Golgi network to the center of the cell division plane (Müller and Jürgens, 2016). Vesicles that accumulate in the division plane fuse together, forming the cellular plate, which expands radially toward the plasma membrane of the parental cell. Fusion between the cellular plate and plasma membrane occurs at the cortical division site where the preprophase band—a densely packed matrix of actin filaments and antiparallel microtubules—is formed. Finally, callose and cellulose synthases modify the composition of the newly formed cell wall, increasing its rigidity and physically delimiting the daughter cells (Müller and Jürgens, 2016).

Proteins involved in cytokinesis in Arabidopsis (*Arabidopsis thaliana*) are classified into three categories: proteins responsible for the proper orientation of the division plane, proteins directly involved in executing cytokinesis, and proteins involved in cell wall biosynthesis (Thiele et al., 2009). The first category includes proteins such as TONNEAU1 (TON1) and TON2 (Struk and Dhonukshe, 2014), TON1 RECRUITING MOTIF1 (TRM1; Drevensek et al., 2012), and TANGLED1 (TAN1; Smith et al., 1996). The second category comprises proteins that organize antiparallel microtubules, such as the kinesin RADIALLY SWOLLEN7 (RSW7; Wiedemeier et al., 2002) and the tubulin folding cofactors PILZ (Steinborn et al., 2002); proteins involved in phragmoplast reorganization, such as the microtubule-associated kinase RUNKEL (RUK; Krupnova et al., 2009), the kinesin HINKEL (HIK; Strompen et al., 2002), and the microtubule-associated protein PLEIADE (PLE; Müller et al., 2002); and proteins that participate in vesicle fusion at the cellular plate, such as the Transport Protein Particle II complex (TRAPPII), the tethering factor CLUB (Söllner et al., 2002), the syntaxin KNOLLE (KN; Lukowitz et al., 1996), and the SM protein KEULE (KEU; Assaad et al., 2001). The third category includes callose synthases such as MASSUE (MAS; Thiele et al., 2009) and CALLOSE SYNTHASE1 (CALS1; Hong et al., 2001), cellulose synthases such as PROSCUTE1 (PRC1; Fagard et al., 2000) and RADIALLY SWOLLEN1 (RSW1; Williamson et al., 2001), the GDP-mannose pyrophosphorylase CYTOKINESIS DEFECTIVE1 (CYT1; Nickle and Meinke, 1998), and the endo-1,4-beta-D-glucanase KORRIGAN (KOR; Zuo et al., 2000).

The formation of the cellular plate during cytokinesis occurs through the fusion of vesicles from the trans-Golgi network. Trafficking of these vesicles begins during anaphase, when the phragmoplast transports the vesicles to the cell division plane along with other membrane and cell wall components. In Arabidopsis, the Soluble N-ethylmaleimide-Sensitive Factor (NSF) Attachment Protein Receptor complexes (SNARE) play a central role in vesicle fusion (Jürgens, 2005). These complexes, along with other components of the vesicle fusion machinery, are highly conserved in eukaryotic cells and have been studied in many species (Blatt et al., 1999).

Vesicle fusion requires proper tethering and docking (Lupashin and Sztul, 2005). During tethering, a vesicle is captured to keep it close to the cellular membrane with which it will merge (Thellmann et al., 2010). This process is performed by specific proteins with supercoiled helical structures or protein complexes such as TRAPP or the exocyst (Lupashin and Sztul, 2005). During docking, SNARE complexes form between a vesicle and a cell membrane (heterotypic fusion) or between biochemically identical membranes (homotypic fusion), which brings the merging membranes closer together (Park et al., 2012). SNARE complexes consist of four SNARE domains, each containing an alpha-helix anchored to a membrane via its C-terminus. SNARE components anchored to a vesicle are referred to as v-SNARE or R-SNARE, as they possess an arginine residue at the center of the SNARE domain. SNARE components anchored to a target membrane are known as t-SNARE or Q-SNARE, as they contain a glutamic acid residue at the same position. The SNARE complex consists of four components: a SNARE domain from an R-SNARE, another from a syntaxin (Qa-SNARE domain), and the remaining two from a Synaptosomal-Associated Protein 25 protein (SNAP25; Qbc-SNARE domain) or from two independent proteins (Qb-SNARE and Qc-SNARE domains) (Fukuda et al., 2000).

Syntaxins can adopt two conformations: closed or open. In the closed conformation, the Habc N-terminal domain, consisting of three alpha-helices, folds over its SNARE domain. In the open conformation, they remain separated (Margittai et al., 2003). The syntaxin KNOLLE plays a central role in cytokinesis. KNOLLE is transported by trans-Golgi vesicles in cis-SNARE complexes with the Qbc-SNARE SOLUBLE N-ETHYLMALEIMIDE-SENSITIVE FACTOR ADAPTOR PROTEIN33 (AtSNAP33), which is the Arabidopsis homolog of human SNAP25 (Heese et al., 2001) and the R-SNARE VESICLE-ASSOCIATED MEMBRANE PROTEIN721 (VAMP721). At the division plane, the ATPase NSF separates the components of the cis-SNARE complexes, and KNOLLE adopts a closed conformation, preventing its interaction with other SNARE components. During cytokinesis, the presence of KEULE stabilizes the open conformation of KNOLLE by interacting with the linker region between its domains (Park et al., 2012). This stabilization promotes the formation of trans-SNARE complexes with the SNARE components of adjacent vesicles (Park et al., 2012). KEULE also interacts with other syntaxins in heterotypic fusions, such as the closed conformation of SYNTAXIN RELATED PROTEIN1 (SYP121), which participates in vesicle trafficking of the cell membrane during drought stress and pathogen defense (Karnik et al., 2013).

All Arabidopsis *keule* (*keu*) mutants studied to date are seedling lethal and have large plurinucleated cells with incomplete cell walls in dividing embryonic cell populations (Assaad et al., 1996). Seedlings with the same *keu* genotype can exhibit various morphologies, ranging from balls of undifferentiated cells to seedlings with well-defined organs, although most are rod-shaped (Assaad et al., 1996). Seedlings with the most severe mutant phenotype exhibit a swollen epidermis, with poorly defined cellular layers containing undifferentiated cells in both the epidermis and inner tissues. This is not observed in seedlings with milder morphologies. The division planes in the *keu* mutants are improperly oriented, and cytokinesis occurs at a slower pace (Assaad et al., 1996). During embryogenesis, the division of the smallest daughter cell of the zygote occurs perpendicular instead of parallel to the apical-basal axis. Additionally, the asynchronous tangential divisions of the zygote often lead to the partial or total loss of the protoderm. However, these embryos usually recover this aspect of the wild-type phenotype during development (Assaad et al., 1996).

Here, we describe two new hypomorphic alleles of the Arabidopsis *KEU* gene, *serrata4-1* (*sea4-1*) and *sea4-2*, which are viable and fertile. The *sea4-1* mutation causes retention of the 9^th^ intron of *KEU*, while the *sea4-2* mutation is predicted to induce an amino acid substitution in the SNARE interaction domain of KEU. Both mutants display reduced rosette size and plant height, abnormal leaf venation patterns, variable cell sizes within the epidermis and palisade mesophyll, and apparently undifferentiated cells throughout these layers, as well as premature leaf senescence. Our findings shed light on some functions of *KEULE* in adult plants, paving the way for future investigations of the postembryonic roles of this essential gene.

## 2. MATERIAL AND METHODS

### 2.1. Plant materials, growth conditions, and crosses

The *Arabidopsis thaliana* (L.) Heynh. wild-type accessions Landsberg *erecta* (L*er*) and Columbia-0 (Col-0), along with the *keu-21* (SALK_101874C; N661055), *keu-22* (SALK_085463; N585463), *keu-23* (SALKseq_089213; N589213), *sec1b-1* (GK-601G09; N457681; previously named *sec1b* in Karnahl et al., 2018), *sec1b-2* (GK-283F10; N427142), *syp21* (SAIL_580_C04; N875090), *syp132^T^* (SAIL_403_B09; N818666), *sec6-2* (SALK_072337C; N660954) and *sec6-3* (SALK_100970; N600970) mutants, were obtained from the Nottingham Arabidopsis Stock Center (NASC; Nottingham, United Kingdom). The *keu^MM125^*and *kn^X37-2^* lines (Karnahl et al., 2018) were kindly provided by Gerd Jürgens. The *sea4-1* and *sea4-2* lines were isolated in the L*er* background after ethyl EMS mutagenesis in our laboratory. Subsequently, they were backcrossed twice to L*er* (Berná et al., 1999). Unless otherwise specified, all the mutants mentioned in this work are homozygous for the indicated mutations. Seed sterilization and sowing, plant culture, crosses, and allelism tests were conducted as previously described (Ponce et al., 1998; Berná et al., 1999; Quesada et al., 2000).

### 2.2. Positional cloning and molecular characterization of the *sea4* mutant alleles

Genomic DNA extraction was carried out as previously described (Ponce et al., 2006). The *sea4-1* and *sea4-2* mutations were initially mapped to a 660-kb candidate interval containing 250 genes using a mapping population of 79 F_2_ plants derived from *sea4-1* × Col-0 and *sea4-2* × Col-0 outcrosses and the primers listed in Table S1, as previously described (Ponce et al., 1999; Ponce et al., 2006). Subsequently, the complete genomes of *sea4-1* and *sea4-2* were sequenced by BGI (Shenzhen, China) using the BGISEQ platform. The raw data were analyzed using Easymap v.2 (Lup et al., 2021; Lup et al., 2022; Lup et al., 2023). Reads were aligned to the L*er* genome (Zapata et al., 2016); the options used were based on the facts that the mutations under study were present in the reference genetic background, the mapping population resulted from a backcross, and the control sample was the parental line of the mutant strain. The reads from *sea4-1* were used as the control sample for *sea4-2*, and vice versa.

For plant genotyping, the wild-type *KEULE* and *sea4* mutant alleles were identified by PCR using the At1g12360-5F/R primers (Table S1). The presence of T-DNA insertions in the insertional lines was confirmed by PCR, and their positions were determined via Sanger sequencing. Gene- and T-DNA-specific primers (Table S1) were employed for these analyses.

### 2.3. Phenotypic analysis and morphometry

Rosettes were photographed using a Nikon SMZ1500 stereomicroscope equipped with a Nikon DXM1200F digital camera. Root length was measured from photographs taken with a Canon PowerShot SX200 IS digital camera using the NIS Elements AR 3.1 image analysis package (Nikon). Shoot length from the soil to the apex of the main shoot was measured *in vivo* using a millimeter ruler. Whole plants were photographed with a Canon PowerShot SX200 IS digital camera.

### 2.4. Differential interference contrast and bright-field microscopy

For bright-field microscopy, all samples were cleared and mounted as previously described (Candela et al., 1999). Micrographs of venation patterns were taken under bright field using a Nikon SMZ1500 stereomicroscope equipped with a Nikon DXM1200F digital camera and NIS Elements AR 3.1 software (Nikon). The venation pattern was drawn using Photoshop CS3 on a Cintiq 18SX Interactive Pen Display screen (Wacom, Kazo, Saitama, Japan) and analyzed using phenoVein software (http://www.plant-image-analysis.org; Bühler et al., 2015). For epidermal and palisade mesophyll cell morphometry, leaves were collected and subjected to the following clearing steps: 15 minutes in 90% acetone, 24 h in 70% ethanol, and 24 h in 16 M chloral hydrate at room temperature. Microscopy of leaf tissues was performed using differential interference contrast optics on a Leica DMRB microscope equipped with a Nikon DXM1200F digital camera. Cell contours were manually outlined using Photoshop CS3 on a Cintiq 18SX Interactive Pen Display screen (Wacom, Kazo, Saitama, Japan). Cell area measurements were performed using the NIS Elements AR 3.1 image analysis package (Nikon). Transverse sections were obtained as described in Serrano-Cartagena et al. (2000). The tissue was embedded in Technovit 7100 resin (Kulzer, Hanau, Germany), and 5-μm sections were cut using a Microm HM350S microtome (Walldorf, Germany).

### 2.5. Confocal microscopy

Confocal laser scanning microscopy images were obtained with a Leica Stellaris 8 STED confocal microscope equipped with HyD X and HyD SMD detectors, HC PL APO CS2 20x/0.75 DRY and HC PL APO CS2 40x/1.10 WATER objectives, and Leica Application Suite X software (LAS X v.4.5.0.25531; Leica). Visualization of fluorescent proteins was performed on leaf primordia and primary roots mounted on glass slides in deionized water. GFP was excited at 489 nm with a white light laser (WLL). The emission was acquired within the range of 494 nm to 583 nm, and TauSeparation was utilized to distinguish the GFP signal (2 ns) from the chlorophyll autofluorescence signal (0.12 ns). The image resolution was set to 1024 × 1024 pixels, with a speed of 600 Hz, a zoom factor of 0.75, and a line accumulation of 6. Ten optical sections, encompassing the adaxial to the abaxial epidermises of the primordia or the entire root thickness, were photographed and overlapped using LAS X software. The configuration of WLL intensity, transmitted light detector gain, and look-up table values for each photograph type are detailed in Table S2.

### 2.6. RNA-seq analysis

Total RNA was extracted from the samples and subjected to massive parallel sequencing as described in Navarro-Quiles et al. (2022), producing paired-end reads of 150 bp (Table S3). Read mapping to the Arabidopsis genome (TAIR10) was performed using HISAT2 v2.0.5 (Kim et al., 2019) with default parameters, and differentially expressed genes between the *sea4* mutants and L*er* were identified by Novogene using the DESeq2 v1.20.0 R package (Love et al., 2014). Genes with a *p*-value < 0.05 adjusted with the Benjamini and Hochberg’s method and a fold change > 1 were considered to be differentially expressed. Gene Ontology (GO; http://www.geneontology.org/) and Kyoto Encyclopedia of Genes and Genomes (KEGG; http://www.genome.jp/kegg/) pathway enrichment analyses of the differentially expressed genes were performed by Novogene using the clusterProfiler R package. Significantly enriched terms were determined with an adjusted *p*-value < 0.05.

### 2.7. Ploidy analysis

Flow-cytometry analysis was performed as previously described (Desvoyes et al., 2006). Briefly, first-node and second-node leaves from four different rosettes were harvested 21 das and chopped with a razor blade in 500 μl of cold nuclear isolation buffer (Galbraith et al., 1991). The cell suspension was filtered through a 30-μm nylon mesh, treated with RNase A (200 μg/ml) for 20 min, and stained with propidium iodide (50 μg/ml) for 40 min. The nuclear DNA content was analyzed using a FACS Canto II flow cytometer (BD Biosciences; https://www.bdbiosciences.com), and the data were processed using the Floreada.io web platform (https://floreada.io). Three biological replicates (15,000 counts each) were analyzed per genotype.

### 2.8. Protein sequence alignment

The protein sequences of Arabidopsis KEU and rat SYNTAXIN-BINDING PROTEIN1 (STXBP1) were obtained from the National Center for Biotechnology Information database (NCBI; http://www.ncbi.nlm.nih.gov/; KEU: NP_563905.1; rat STXBP1: NP_037170.1). Multiple sequence alignment was performed using Clustal Omega (https://www.ebi.ac.uk/Tools/msa/clustalo/; Madeira et al., 2019).

### 2.9. Protein structure visualization and analysis

The 3D structures of the full-length monomers of KEU and rat STXBP1 were obtained from the AlphaFold Protein Structure Database (https://alphafold.ebi.ac.uk/; Tunyasuvunakool et al., 2021; Varadi et al., 2022; KEU: AF-Q9C5X3-F1; rat STXBP1: AF-P61765-F1) and visualized using UCSF ChimeraX 1.2.5 software (https://www.rbvi.ucsf.edu/chimerax/; Goddard et al., 2018; Pettersen et al., 2021). To analyze the impact of the S57L substitution in *sea4-2* on the conformational stability and dynamics of KEU, two web structure-based protein stability predictors were used: DynaMut (https://biosig.lab.uq.edu.au/dynamut/; Rodrigues et al., 2018) and DynaMut2 (https://biosig.lab.uq.edu.au/dynamut2/; Rodrigues et al., 2021). These predictors quantify the difference in unfolding Gibbs free energy (ΔΔG, expressed in kcal·mol^−1^) between wild-type and mutant proteins, classifying mutations as stabilizing when ΔΔG > 0 kcal·mol^−1^ or destabilizing when ΔΔG < 0 kcal·mol^−1^. DynaMut provides ΔΔG results from three additional predictors: SDM (Worth et al., 2011), mCSM (Pires et al., 2014b), and DUET (Pires et al., 2014a). Furthermore, it computes the difference in vibrational entropy energy (ΔΔS_Vib_, expressed in kcal·mol^−1^·K^−1^) between wild-type and mutant proteins using the ENCoM server (Frappier et al., 2015), classifying mutations as rigidifying if ΔΔS_Vib_ < 0 kcal·mol^−1^·K^−1^ or flexibilizing if ΔΔS_Vib_ > 0 kcal·mol^−1^·K^−1^. Finally, Missense3D (http://missense3d.bc.ic.ac.uk/missense3d/; Ittisoponpisan et al., 2019) was used to predict damaging structural effects of the S57L substitution on KEU protein.

### 2.10. Accession numbers

Sequence data from this article can be found at The Arabidopsis Information Resource (https://www.arabidopsis.org/) under the following accession numbers: KEU (At1g12360), KN (At1g08560), SEC1B (At4g12120), SYP21 (At5g16830), SYP132 (At5g08080), and SEC6 (At1g71820).

## 3. RESULTS

### 3.1. The *sea4* mutants carry novel hypomorphic viable alleles of *KEU*

In a large-scale screening for Arabidopsis leaf mutants conducted after ethyl methanesulfonate (EMS) mutagenesis of Landsberg *erecta* (L*er*) seeds (Berná et al., 1999), we isolated two mutants displaying serrated and undulated rosette leaves with necrotic patches. These mutants were found to be allelic and were named *serrata4-1* (*sea4-1*) and *sea4-2*. Initially, the *sea4-1* and *sea4-2* mutations were mapped at low resolution to a 660-kb candidate region of chromosome 1 containing approximately 250 genes (Figure 1A) using iterative linkage analysis to molecular markers (Table S1; Robles and Micol, 2001). We generated backcross mapping populations by selecting F_2_ plants exhibiting the mutant phenotype and subjected their pooled genomic DNA to next-generation sequencing. We analyzed the resulting paired-end reads using Easymap v.2, which also pointed to the same region of chromosome 1 (Figure 1B). By comparing the lists of candidate mutations identified in *sea4-1* (Table S4) and *sea4-2* (Table S5), we identified At1g12360 as the most likely candidate gene, which encodes the Sec1/Munc18 protein KEULE (KEU), as both mutants carried a mutation in this gene.

**Figure 1.**
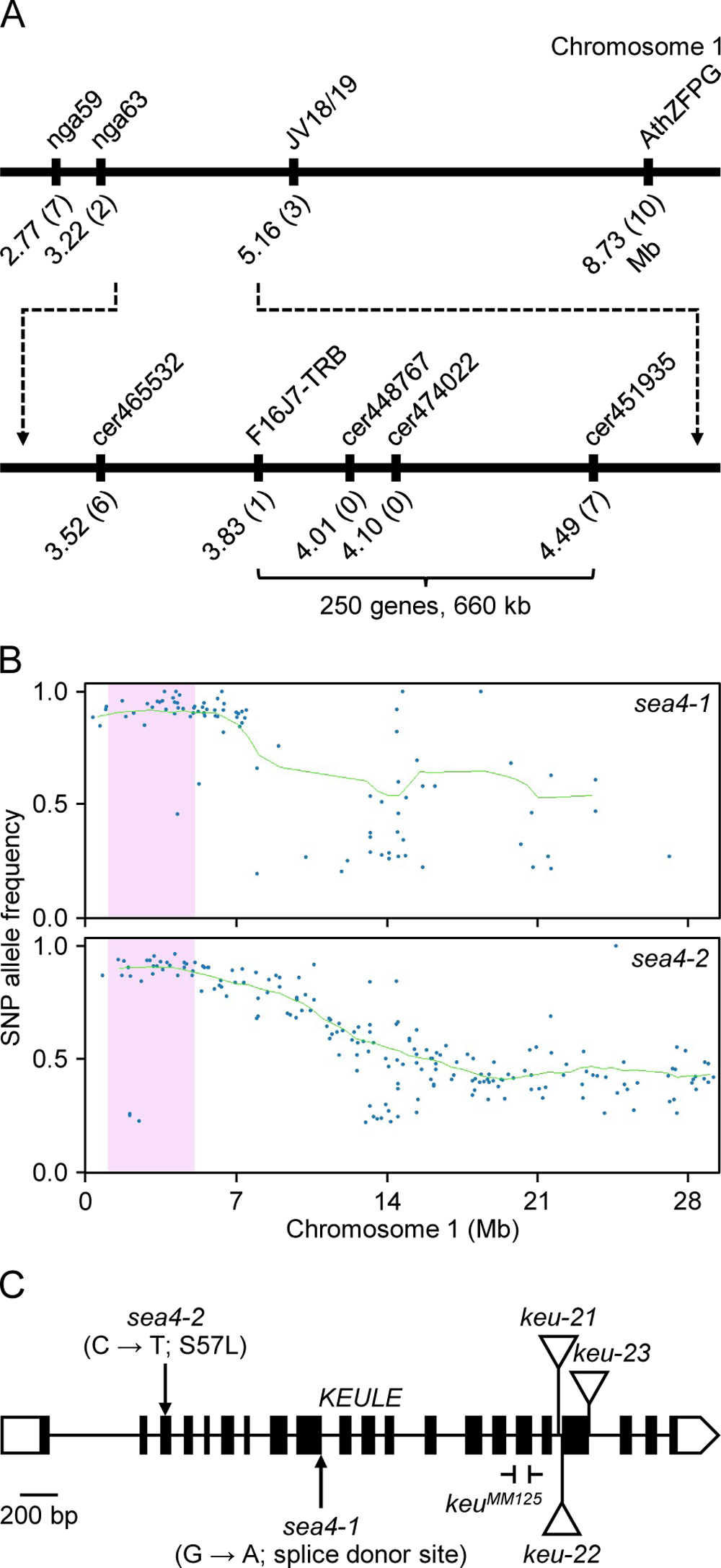
Mapping of the *sea4-1* and *sea4-2* mutations. (A) Analysis of a mapping population of 79 F_2_ plants derived from a *sea4-1* × Col-0 cross revealed a candidate interval of 660 kb on chromosome 1 containing 250 genes. The names and physical map positions of the molecular markers used for linkage analysis are shown. The number of recombinant chromosomes found is indicated in parentheses. (B) Plots showing the allele frequency of single nucleotide polymorphisms (SNPs) versus positions in a mapping-by-sequencing analysis of the *sea4* mutants performed using Easymap v.2. SNPs are represented as blue dots, the candidate region is shaded in pink, and the average allele frequency (AF) in the test sample of SNPs used for mapping is represented by a green line. (C) Structure of the *KEU* gene showing the nature and positions of the *sea4* and *keu* mutations studied in this work. Boxes and lines between boxes indicate exons and introns, respectively. White boxes represent the 5′- and 3′- UTRs. Triangles indicate the T-DNA insertions in *keu-21*, *keu-22,* and *keu-23*, and the Ͱ symbols delimitate the deletion in *keu^MM125^*.

We obtained three lines from the SALK collection that were annotated to carry T-DNA insertions within the At1g12360 transcription unit: SALK_101874C, SALKseq_085463, and SALKseq_089213. We renamed the mutations in these lines *keu-21*, *keu-22*, and *keu-23*, respectively (Figure 1C). Additionally, Gerd Jürgens’ laboratory provided us with the line *keu^MM125^* (Karnahl et al., 2018), which carries a 100-bp deletion in the 16^th^ exon of *KEU* (Figure 1C). As expected, plants homozygous for these putatively null *keu* alleles were embryonic lethal, and the rosettes of the *KEU/keu-21*, *KEU/keu-22*, *KEU/keu-23*, and *KEU/keu^MM125^* heterozygotes were phenotypically wild type. Crosses of *KEU/keu-21*, *KEU/keu-22*, and *KEU/keu-23* heterozygotes to homozygous *sea4* plants produced viable heterozygous *sea4/keu* plants, exhibiting a mutant phenotype more severe than that of *sea4-1/sea4-1* or *sea4-2/sea4-2* homozygotes (Figure 2). Crosses of *KEU/keu^MM125^* heterozygotes to homozygous *sea4* plants produced heterozygous *sea4/keu^MM125^* plants displaying a similar phenotype (Figure 2), but they died at the end of the vegetative phase without undergoing bolting or producing inflorescences. These results confirm the notion that *sea4-1* and *sea4-2* are hypomorphic alleles of *KEU*.

**Figure 2.**
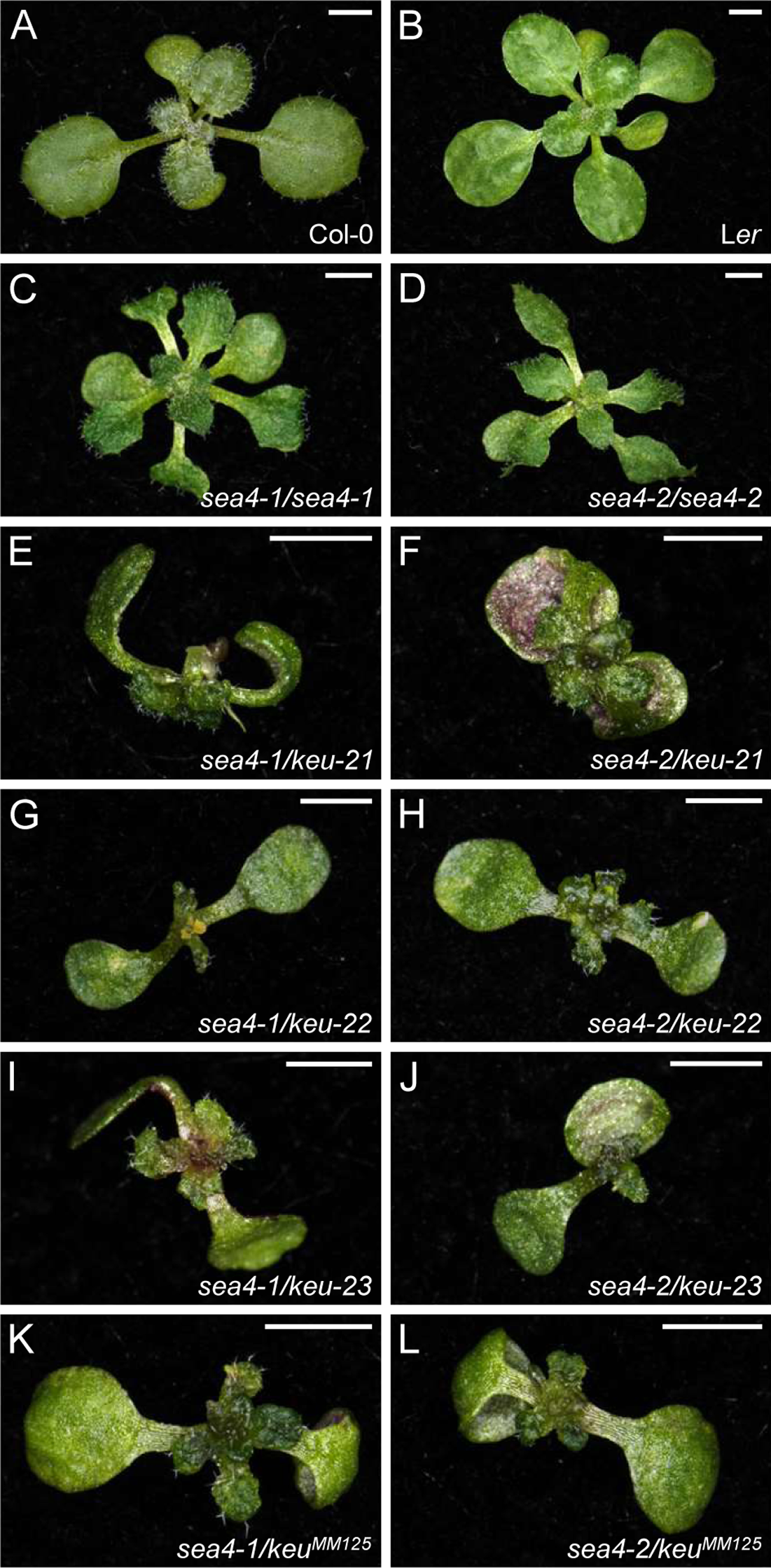
Leaf phenotypes and complementation analysis of the *sea4* and *keu* mutants studied in this work. Rosettes of wild-type (A) Col-0 and (B) L*er*, the (C) *sea4-1/sea4-1* and (D) *sea4-2/sea4-2* homozygous mutants, and the (E) *sea4-1/keu-21*, (F) *sea4-2/keu-21*, (G) *sea4-1/keu-22*, (H) *sea4-2/keu-22*, (I) *sea4-1/keu-23*, (J) *sea4-2/keu-23*, (K) *sea4-1/keu^MM125^*, and (L) *sea4-2/keu^MM125^* heterozygotes. Photographs were taken 14 das. Scale bars: 2 mm.

### 3.2. The *sea4-1* mutation causes mis-splicing of *KEU*, and *sea4-2* appears to perturb the secondary structure of KEU

The Arabidopsis *KEU* gene is expressed throughout the plant, especially in tissues undergoing division (Assaad et al., 2001). KEU exists in soluble form in the cytoplasm or associates with membranes during cytokinesis (Assaad et al., 2001). KEU is involved in cytokinesis, cellular elongation, and root hair growth (Assaad et al., 2001). KEU shares 28-30% identity with its Sec1 orthologs in mammals, *Caenorhabditis elegans*, and *Drosophila melanogaster* and 61% and 65% identity with its Arabidopsis homologs SEC1A and SEC1B, respectively (Assaad et al., 2001).

In *sea4-1*, a G→A transition at the 3′ end of the ninth exon of *KEU* damages the splice donor site. In *sea4-2*, a C→T transition in the third exon is predicted to result in an S57L substitution (Figure 1C). Since *sea4-1* is likely to undergo mis-splicing and produce abnormal mRNA and protein products, we conducted RNA-seq analysis of this line, finding that 99.17% of its mature mRNAs retained the 9^th^ intron (Table S6), leading to a premature stop codon. The mutant SEA4-1 protein consists of 302 amino acids, 38 of which are absent from wild-type KEU (Figure S1). The expression level of *KEU* was five times higher in *sea4-1* than in L*er*, as indicated by the number of RNA-seq reads aligned to *KEU*, likely to compensate for the near absence of wild-type mRNAs (Table S7).

Alignment of the KEU sequence with its rat STXBP1 ortholog revealed that the *sea4-2* mutation alters one of the amino acids that physically interact with syntaxins (Figure S2; Misura et al., 2000). To assess the impact of this mutation on the structural stability and dynamics of KEU, we used Dynamut and DynaMut2 to predict differences in the unfolding Gibbs free energy (ΔΔG) and vibrational entropy energy (ΔΔS_Vib_) between the wild-type and mutant proteins. We obtained the structure of the full-length KEU protein from the AlphaFold Protein Structure Database. However, these two predictors provided conflicting ΔΔG values, suggesting that the S57L mutation does not affect protein stability (Table S8). ENCoM calculated a decrease in flexibility of KEU due to this mutation based on the ΔΔS_Vib_ value (Table S8 and Figure S3). To predict the effects of the S57L mutation on KEU protein structure, we used Missense 3D. This software did not predict any structural damage to KEU, since both residues were exposed to solvents in a similar manner. However, two hydrogen bonds between S57 and its neighboring residues V53 and K54 were lost (Figure S4), and the cavity volume increased by 24.408 Å^3^.

### 3.3. Morphological and histological characterization of the *sea4* mutants

#### 3.3.1. The *sea4* mutants exhibit a pleiotropic morphological phenotype

Both *sea4* mutants exhibited undulated leaves with serrated margins and senescent patches (Figure 2). Both *sea4-1* and *sea4-2* displayed phenotypic variability, including varying rates of cotyledon and leaf expansion and variations in the extent of senescent patches (Table S9). A similar phenotypic variability was observed in lethal seedlings carrying other *KEU* alleles (Assaad et al., 1996) and reappeared in the progeny of selfed *sea4* plants, regardless of the parental phenotype (Figure S5). Inflorescences of *sea4* plants contained fertile flowers that opened prematurely before complete maturation (Figure S6E and S6G). Inflorescences of *KEU/keu-21*, *KEU/keu-22*, and *KEU/keu-23* plants were indistinguishable from those of Col-0 (Figure S6I, S6K and S6M). However, in inflorescences of heterozygotes of *sea4-1* or *sea4-2* with *keu21*, *keu22* or *keu23*, only a few flowers opened prematurely. These flowers exhibited short sepals that were separated from each other and had protuberances on their margins, but they remained fertile (Figure S6Q, S6S, S6U, S6W, S6Y and S6AA). Inflorescences of *KEU/keu^MM125^* plants exhibited a few fertile flowers that began to open slightly before reaching complete maturation (Figure S6O), while heterozygous *sea4/keu^MM125^* plants did not produce inflorescences. Inflorescences of *sea4-1/sea4-2* plants contained fewer flowers than those of homozygous *sea4* mutants, but they exhibited the same phenotypes (Figure S6AC). The height of *sea4-1*, *sea4-2*, and *KEU/keu^MM125^*plants was reduced compared to L*er*, whereas *KEU/keu-21*, *KEU/keu-22*, and *KEU/keu-23* plants did not exhibit a significant difference from the wild type. *sea4-1/sea4-2* plants displayed an intermediate height between that of *sea4-1* and *sea4-2* homozygous plants. Heterozygous *sea4/keu* plants were even smaller, and individuals carrying a *sea4-1* allele were smaller than those carrying a *sea4-2* allele, which is consistent with the smaller height observed in *sea4-1* vs. *sea4-2* plants (Figure S7). The primary root length was also reduced in the *sea4* mutants compared to the wild type (Figure S8).

#### 3.3.2. The venation pattern of *sea4* leaves is dense and complex

The presence of serrations in *sea4* leaves pointed to the existence of other alterations in the internal structures of these leaves. To observe possible alterations in the venation pattern, we decolorized cotyledons, first-node leaves, and third-node leaves of *sea4-1* (n = 12) and *sea4-2* (n = 13–15) plants and compared their venation patterns to L*er* (n = 12–15) (Figure 3). The *sea4-1* cotyledons showed an increased number of terminal veins per unit area compared to L*er*, whereas *sea4-2* cotyledons were smaller and exhibited increased vein length and bifurcation number per unit area. The first-node leaves of *sea4-1* and *sea4-2* were elongated, smaller, and contained more terminal veins per unit area than L*er*. Additionally, *sea4-1* first-node leaves displayed increased vein length and bifurcations per unit area compared to L*er*. The third-node leaves of *sea4-1* and *sea4-2* were smaller, with an increase in terminal veins per unit area compared to L*er*. In *sea4-2*, the leaves were also elongated and exhibited increased vein length and bifurcations per unit area compared to L*er* (Table S10). Overall, the cotyledons and leaves of the *sea4* mutants were smaller and elongated compared to the wild type. They showed a higher vein density, and their venation patterns were more complex, with a greater number of bifurcations and terminal veins per unit area.

**Figure 3.**
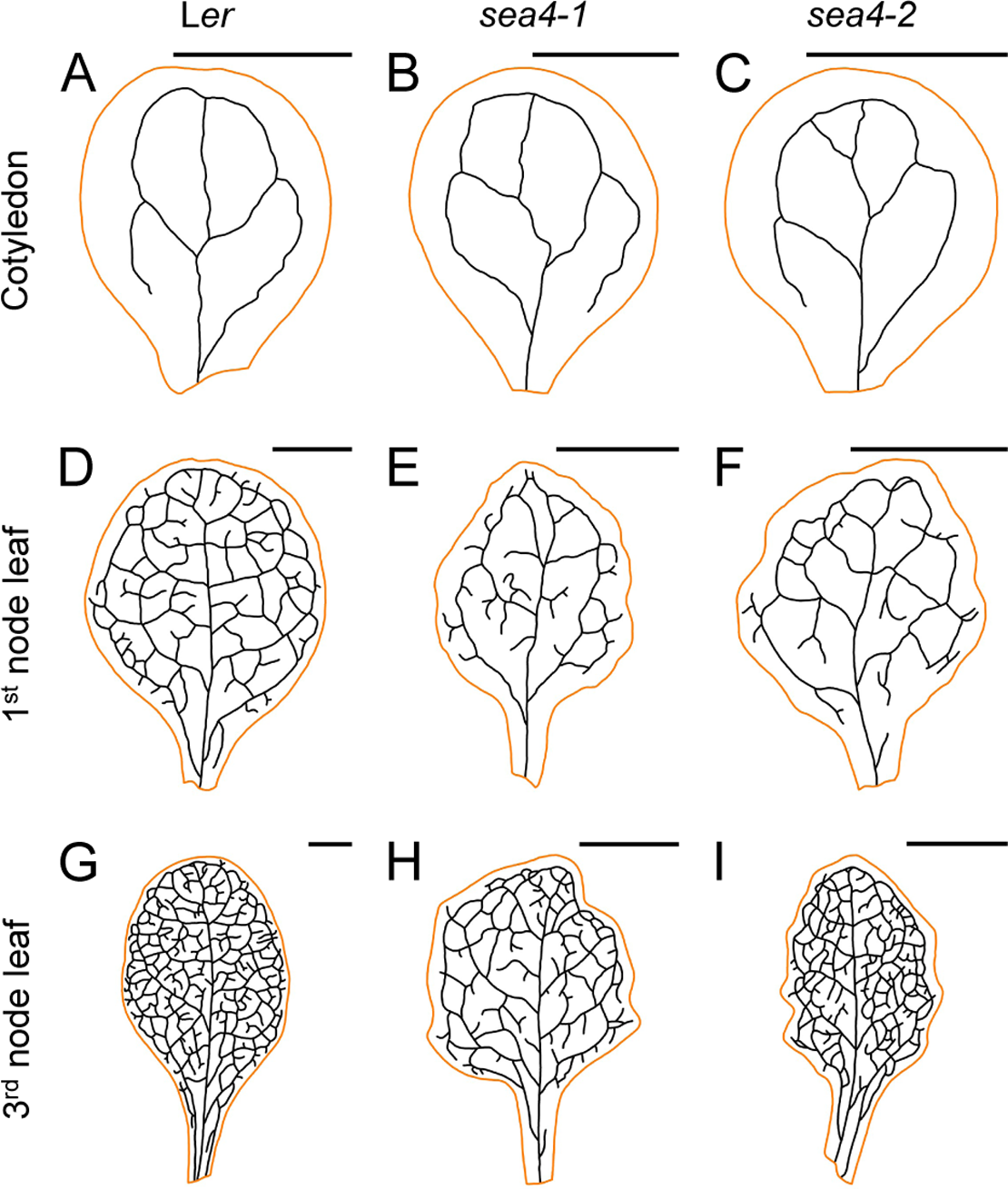
Venation patterns of *sea4-1* and *sea4-2* cotyledons and first- and third-node leaves. Representative diagrams of fully expanded (A, B, C) cotyledons, (D, E, F) first-node leaves, and (G, H, I) third-node leaves from (A, D, G) L*er*, (B, E, H) *sea4-1*, and (C, F, I) *sea4-2* plants. Margins were drawn in orange and veins in black. The line thickness does not represent the actual thickness of the veins. Cotyledons and leaves were collected 21 das. Scale bars: 2 mm.

#### 3.3.3. *sea4* leaves show aberrant epidermal and mesophyll structure

We obtained transverse sections of third-node leaves from *sea4-1*, *sea4-2*, and L*er* plants (Figure 4). In the palisade mesophyll of the *sea4* mutants, the cells were generally well organized but were more variable in size compared to L*er*. Some large cells were interspersed among small cells within the palisade mesophyll. The spongy mesophyll in the *sea4* mutants was disorganized, with very large cells surrounded by small cells arranged in multiple layers. The epidermal cells appeared mostly normal, although protuberances were observed on the surfaces of mutant leaves. These protuberances appeared to result from increased local proliferation or growth of mesophyll cells.

**Figure 4.**
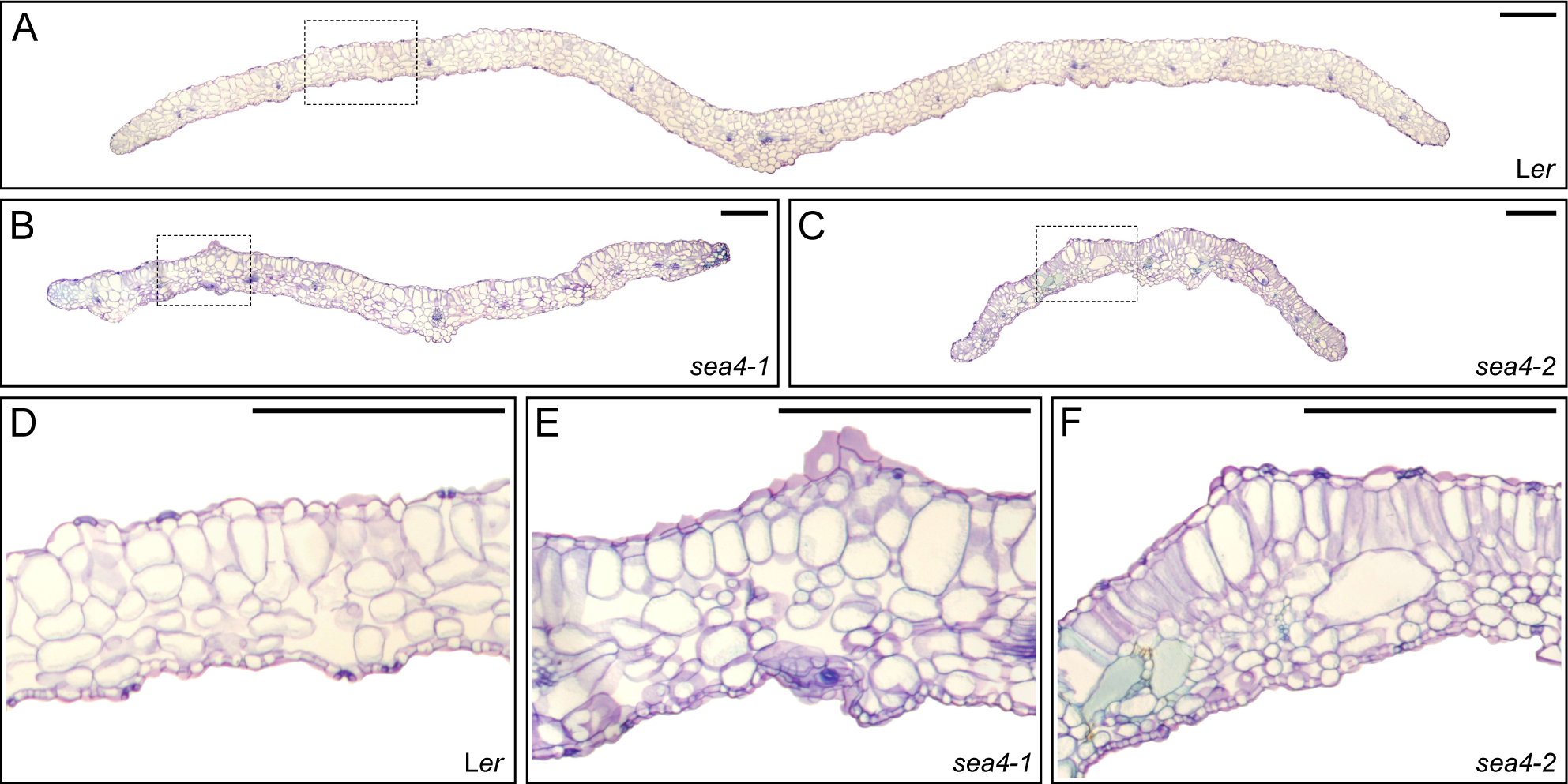
Internal structure of *sea4* third-node leaves. Transverse sections of leaves from (A, D) L*er*, (B, E) *sea4-1*, and (C, F) *sea4-2* plants stained with toluidine blue. Photographs show (A-C) a complete transverse section of the lamina from margin to margin, and (D-F) the central zone of the lamina between the primary vein and margin, which is marked with a rectangle in A, B, and C. Photographs were taken 21 das. Scale bars: 1 mm.

We quantified the variation in palisade mesophyll, adaxial epidermis, and abaxial epidermis cells in transverse sections of first- and third-node leaves (Figure 5 and Figure S9). In the palisade mesophyll of the two leaves analyzed, most *sea4* cells were smaller than L*er* cells, but a small fraction of *sea4* cells were larger. In the epidermis, cells tended to have a similar size, although *sea4* cells had a simpler shape compared to L*er* cells, with fewer protuberances observed in both first- and third-node leaves.

**Figure 5.**
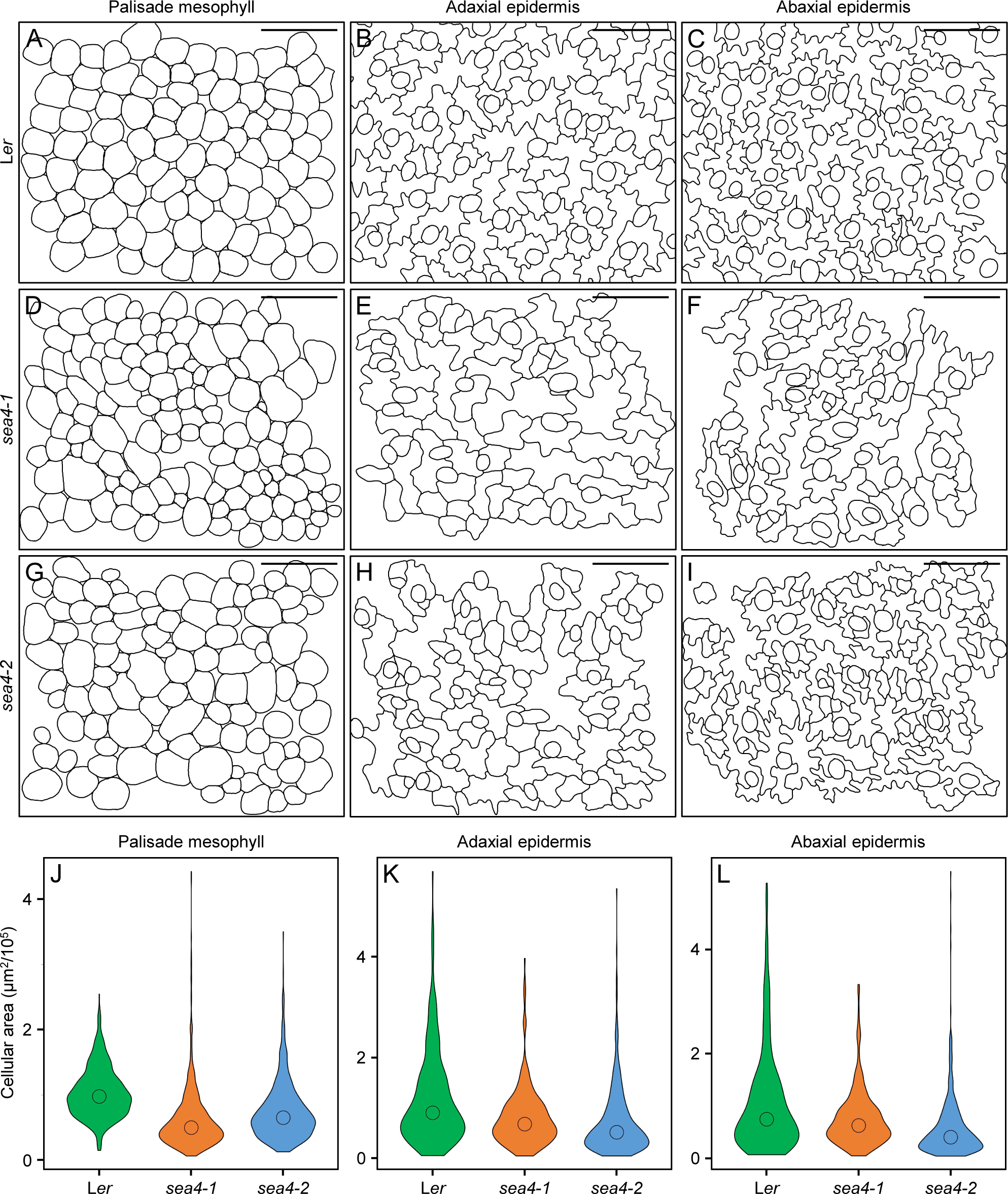
Structure of the cell layers in third-node leaves of the *sea4* mutants. (A-I) Diagrams of cells from the (A, D, G) palisade mesophyll, (B, E, H) adaxial epidermis, and (C, F, I) abaxial epidermis of (A, B, C) L*er*, (D, E, F) *sea4-1*, and (G, H, I) *sea4-2* plants. Leaves were collected 21 das. Scale bars: 20 μm. (J-L) Violin plots representing the distribution of cell sizes in the (J) palisade mesophyll, (K) adaxial epidermis, and (L) abaxial epidermis. The median is represented with a circle.

#### 3.3.4. The *sea4* mutants show early leaf senescence and bolting

A characteristic trait of the *sea4* mutants is the early appearance of senescent patches in their leaves, which gradually expanded until they were completely senescent. To quantify the degree of senescence, we grouped all leaves from *sea4-1* (n = 38), *sea4-2* (n = 45), and L*er* (n = 44) plants into five phenotypic classes based on their degree of senescence at 25 days after stratification (das; Figure S10). L*er* plants displayed some senescence in their first four leaves, particularly in the first and second nodes (Figure S10F). *sea4-1* leaves exhibited senescence from the first to eighth nodes (Figure S10G), with all leaves showing more severe senescence compared to L*er*. Similarly, *sea4-2* plants displayed senescence in their first seven leaves (Figure S10H), with leaves in the first and second nodes exhibiting more severe senescence than L*er*. In conclusion, *sea4* plants demonstrated a more pronounced state of senescence in their initial leaves compared to the wild type and exhibited senescence starting from the fifth-node leaves onwards, which was not observed in L*er*.

Senescence appeared earlier in *sea4* than in the wild type. The first signs of senescence appeared ∼14 das in *sea4-1* (n = 95) and *sea4-2* (n = 96) plants and at ∼21 das in L*er* (n = 79; Figure S11A). The early onset of senescence was consistent across different phenotypic classes in the *sea4* mutants (Figure S11A) and occurred at the same time in both *sea4-1* and *sea4-2* (Figure S11C).

Only *sea4-2* exhibited a significant reduction in bolting time compared to L*er* (Figure S11B). When considering the phenotypic classes of the *sea4* mutants, there was a positive correlation between phenotype severity and an increase in bolting time (Figure S11B).

### 3.4. Double mutant combinations of *sea4* alleles and alleles of genes encoding proteins required for membrane fusion show defects in leaf development

As the *sea4* mutants carry the first known viable alleles of *KEU*, we combined them with alleles of other genes involved in membrane fusion to characterize their genetic interactions in adult plants. *SEC1B* is a *KEU* homolog in Arabidopsis that is expressed at very low levels, but its overexpression rescued the phenotype of *keu* mutants (Karnahl et al., 2018). We obtained two lines, *sec1b-1* and *sec1b-2*, carrying T-DNA insertions interrupting *SEC1B* in the 12^th^ intron and 1^st^ exon, respectively (Figure S12A), and showing a wild-type phenotype in the homozygous state (Figure 6E-F). The *sea4 sec1b-1* double mutants displayed small rosettes with small, highly serrated leaves. The *sea4 sec1b-2* double mutants were very small, contained fully expanded cotyledons, and lacked true leaves, instead showing small protuberances emerging from the shoot apical meristem (Figure 6L-O). *SEC6* encodes a protein that is part of the exocyst complex, which colocalizes with KEU in the cellular plate (Wu et al., 2013). We also obtained the lines *sec6-2* and *sec6-3*, carrying T-DNA insertions in the 23^rd^ and 26^th^ exons of *SEC6*, respectively (Figure S12B), which presented a wild-type phenotype in the homozygous state (Figure 6G-H). The *sea4-1 sec6-2* and *sea4-1 sec6-3* double mutants displayed a reduced size with small, undulated, heavily serrated leaves, and *sea4-2 sec6-2* and *sea4-2 sec6-3* exhibited a small size, with their leaves showing minimal or no development (Figure 6P-S).

**Figure 6.**
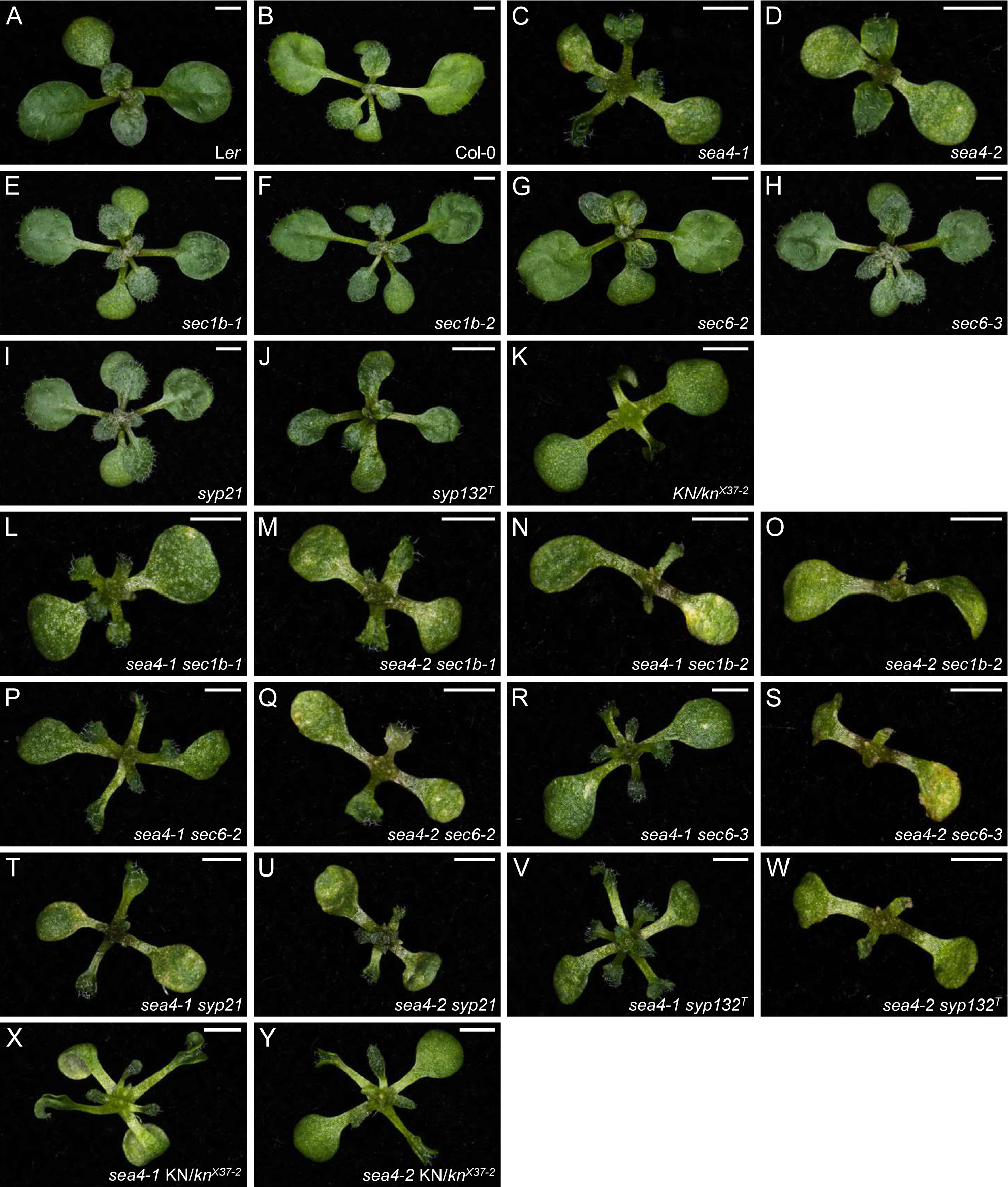
Leaf phenotypes of double mutants between the *sea4* mutants and mutants carrying alleles of genes involved in membrane fusion. Rosettes of wild-type (A) L*er* and (B) Col-0; the (C) *sea4-1*, (D) *sea4-2*, (E) *sec1b-1*, (F) *sec1b-2*, (G) *sec6-2*, (H) *sec6-3*, (I) *syp21*, (J) *syp132^T^*, and (K) *KN/kn^X37-2^* mutants; and (L, M) *sea4 sec1b-1*, (N, O) *sea4 sec1b-2*, (P, Q) *sea4 sec6-2*, (R, S) *sea4 sec6-3*, (T, U) *sea4 syp21*, (V, W) *sea4 syp132^T^*, and (X, Y) *sea4 KN/kn^X37-2^*double mutants. Photographs were taken 14 das. Scale bars: 2 mm.

We also crossed the *sea4* mutants with lines carrying mutations in genes encoding syntaxins: *SYP21*, *SYP132*, and *KNOLLE*. *SYP21* encodes a syntaxin localized to the vacuole and multivesicular bodies that physically interacts with KEU (Park et al., 2012). The *syp21* line carries a T-DNA insertion in the 4^th^ exon of *SYP21* (Figure S12C); homozygous plants were phenotypically wild type (Figure 6I). The *sea4 syp21* double mutants displayed small rosettes with small, highly serrated leaves (Figure 6T-U). *SYP132* encodes a syntaxin that physically interacts with KEU during cytokinesis and with SEC1B in the general secretory pathway (Karnahl et al., 2018). The *syp132^T^*line carries a T-DNA insertion in the 5’ UTR of *SYP132* (Figure S12D); homozygous plants displayed a wild-type phenotype (Figure 6J). The *sea4-1 syp132^T^* double mutants exhibited reduced rosette size, with small, serrated leaves. The *sea4-2 syp132^T^* double mutants showed poorly developed leaves and protuberances emerging from the apical shoot meristem (Figure 6V-W). *KNOLLE* encodes a syntaxin that plays a crucial role in cytokinesis, physically interacting with KEU at the cellular plate (Assaad et al., 2001; Park et al., 2012; Karnahl et al., 2018); this protein accumulated in previously studied *keu* mutants (Waizenegger et al., 2000). The *kn^X37-2^* line has a 1-kb deletion that partially removes the *KN* gene (Figure S12E). Homozygous plants for this deletion were lethal, and heterozygous individuals exhibited small rosettes comprising fully expanded cotyledons and very small leaves that sometimes contained undulations and showed epinasty (Figure 6K). The *sea4/sea4;KN/kn^X37-2^* sesquimutants exhibited a phenotype similar to that of the *KN/kn^X37-2^* heterozygotes, but their leaves were better developed, larger, and displayed undulations and serrations (Figure 6X-Y).

### 3.5. *sea4* leaves show increased endoreduplication

Embryos of other *keu* mutant alleles display multinucleated cells; these nuclei which aggregate in an enlarged nucleus in seedlings (Assaad et al., 1996). To investigate whether adult plant cells exhibit the same phenotype, we conducted flow-cytometry analysis of the first pair of leaves from the *sea4* mutants. As a positive control, we included *den5-1*, which carries a point mutation in *RPL7B* (encoding a ribosomal 60S subunit protein) and shows alterations in ploidy (Horiguchi et al., 2011). Flow cytometry revealed a decrease in populations of cells with lower ploidy levels (2C, 4C and 8C) and an increase in cell populations with higher ploidy levels (16C, 32C, 64C and 128C; Figure S13). These altered ploidy levels were more pronounced in *sea4-1* than in *sea4-2*, which is consistent with the more severe phenotype of *sea4-1* (Table S11). An increase in ploidy levels usually correlates with enlarged cell size (Galbraith et al., 1991), but we did not observed this in the *sea4* mutants. Only a small subset of cells in the mutants were enlarged, and most were smaller than the wild type. Therefore, the increase in ploidy levels in cells of adult leaves is more likely attributed to the aggregation of nuclei in cells unable to complete cytokinesis, as observed in previous studies on lethal seedlings.

### 3.6. Genes related to protein folding, the degradation of misfolded proteins, and plant immunity are upregulated in the *sea4* mutants

To determine whether the *sea4* mutations affect biological processes beyond cytokinesis, we conducted RNA-seq analysis of L*er*, *sea4-1*, and *sea4-2* rosettes collected 14 das. In *sea4-1*, we identified 3812 upregulated and 3824 downregulated genes (Figure S14A and Data Set S1A). In *sea4-2*, we identified 4091 upregulated and 4034 downregulated genes (Figure S14B and Data Set S1B). We classified the differentially expressed genes in *sea4-1* and *sea4-2* by GO (Data Set S2) and KEGG pathway (Data Set S3) analyses separately for upregulated and downregulated genes.

Many upregulated genes in the *sea4* mutants were related to protein folding and the degradation of misfolded proteins in the endoplasmic reticulum (ER). Most genes involved in the sec-dependent protein export pathway, including all those encoding the translocation channel responsible for transporting newly synthesized proteins into the ER (protein export [KEGG:ath03060]), were upregulated in the mutants. Genes related to glycosylation, protein folding, labeling of misfolded proteins, ER-associated degradation, and the unfolded protein response (UPR) were also upregulated in the mutants (protein processing in endoplasmic reticulum [KEGG:ath04141]). Additionally, most genes encoding the 19S regulatory particle and all genes encoding the 20S proteolytic core particle of the proteasome were upregulated in the mutant, as was the gene encoding the PA200 regulatory particle, which stimulates the proteasomal hydrolysis of peptides (proteasome [KEGG:ath03050]). The glycosylation of newly synthesized peptides was also affected in the mutants, as evidenced by the upregulation of genes related to glycan biosynthesis and the oligosaccharyltransferase (OST) complex, which attaches glycan to peptides at the cytoplasmatic face of the ER membrane. Furthermore, some genes involved in modifying glycans within the ER showed increased expression in the mutant (N-glycan biosynthesis [KEGG:ath00510]; various types of N-glycan biosynthesis [KEGG:ath00513]; other types of O-glycan biosynthesis [KEGG:ath00514]).

Other upregulated genes in the *sea4* mutants are related to plant immunity (plant-pathogen interaction [KEGG:ath04626]; MAPK signaling pathway–plant [KEGG:ath04016]; immune response [GO:0006955]; regulation of defense response [GO:0031347]). Among genes involved in pathogen-associated molecular pattern (PAMP)-triggered immunity, most genes of the MAPK signaling pathways triggered by pathogen infection showed increased expression in the mutants (response to bacterium [GO:0009617]; response to fungus [GO:0009620]), leading to the production of ethylene, H_2_O_2_, and cell death (cell death [GO:0008219]). Other genes involved in MAPK signaling pathways triggered by ethylene, reactive oxygen species (ROS), salt, cold, and wounding were also upregulated in the mutants (regulation of response to stress [GO:0080134]; response to wounding [GO:0009611]). Among genes associated with effector-triggered immunity, certain genes associated with the hypersensitive response (HR) to pathogen virulence proteins were upregulated in the mutants, whereas *FDH*, which suppresses HR and defense responses via very-long-chain fatty acids, was downregulated. Genes involved in glucosinolate biosynthesis from methionine and aromatic amino acids, which are defense compounds in plants, were also upregulated in the mutant (glucosinolate biosynthesis [KEGG:ath00966]). Genes in the metabolic pathway of glutathione, which functions as an antioxidant against ROS, were differentially expressed in both *sea4* mutants (glutathione metabolism [KEGG:ath00480]); and genes in the synthesis pathway of alpha-tocopherol and alpha-tocotrienol, which are also antioxidants, were downregulated in the *sea4-1* mutants (ubiquinone and other terpenoid-quinone biosynthesis [KEGG:ath00130]).

Most genes encoding the key enzymes of the tricarboxylic acid cycle were also upregulated in the *sea4* mutants (citrate cycle [KEGG:ath00020]). Additionally, several metabolic pathways responsible for producing important metabolites in plants were upregulated in the mutants. These include the pathway that synthesizes indole-3-acetic acid from tryptophan (tryptophan metabolism [KEGG:ath00380]), the pathway that produces jasmonate from phosphatidylcholine (alpha-linolenic acid metabolism [KEGG:ath00592]), the pathway that releases abscisic acid from abscisic acid glucose ester (carotenoid biosynthesis [KEGG:ath00906]), and the pathway that produces monolignols used in lignin biosynthesis (only in *sea4-2*; phenylpropanoid biosynthesis [KEGG:ath00940]).

### 3.7. Genes related to photosynthesis and the production of various metabolites are downregulated in the *sea4* mutants

Among the downregulated genes in the *sea4* mutants, many are related to the photosynthetic processes (photosynthesis [GO:0015979]; photosynthesis, light reaction [GO:0019684]). Most genes encoding proteins of the F-type ATPase, photosystem I, and photosystem II (photosystem II assembly [GO:0010207]), and all genes encoding proteins of the photosynthetic electron transport chain and the light-harvesting chlorophyll complex (LHC) I and II, were downregulated in the mutants (photosynthesis [KEGG:ath00195]; photosynthesis–antenna proteins [KEGG:ath00196]; photosynthesis, light harvesting [GO:0009765]; photosynthetic electron transport chain [GO:0009767]). Furthermore, pathways responsible for the production of chlorophyll A/B and bacteriochlorophyll A/B from L-glutamate (porphyrin and chlorophyll metabolism [KEGG:ath00860]; chlorophyll biosynthetic process [GO:0015995]; chlorophyll metabolic process [GO:0015994]), as well as alpha/beta-carotene (carotenoid biosynthesis [KEGG:ath00906]), which are essential for photosynthesis, were also downregulated in the mutants. Additionally, genes encoding proteins involved in chlorophyll degradation were upregulated (porphyrin and chlorophyll metabolism [KEGG:ath00860]; porphyrin-containing compound metabolic process [GO:0006778]). In *sea4-1*, most genes of the synthesis pathways of phylloquinone, a photosystem I cofactor, and menaquinone, an electron transport chain component, were downregulated as well (ubiquinone and other terpenoid-quinone biosynthesis [KEGG:ath00130]). Most genes of the Calvin cycle (carbon fixation in photosynthetic organisms [KEGG:ath00710]), as well as the entire pathway that converts guanosine triphosphate (GTP) into flavin mononucleotide (FMN) and flavin adenine dinucleotide (FAD), were also downregulated (riboflavin metabolism [KEGG:ath00740]).

Other downregulated genes in the *sea4* mutants are related to amino acid metabolism. Genes encoding proteins involved in the conversion of glycine to glyoxylate, serine to pyruvate, and aspartate to threonine were downregulated in the mutants, although the pathway responsible for transforming serine into tryptophan was upregulated (glyoxylate and dicarboxylate metabolism [KEGG:ath00630]; glycine, serine and threonine metabolism [KEGG:ath00260]). The pathway involved in converting aspartate to lysine was also downregulated in *sea4-2* (lysine biosynthesis [KEGG:ath00300]). Furthermore, most genes involved in the biosynthesis and degradation of valine, leucine, and isoleucine were downregulated (valine, leucine, and isoleucine biosynthesis [KEGG:ath00290]; valine, leucine, and isoleucine degradation [KEGG:ath00280]). Most genes involved in the biosynthesis of aminoacyl-tRNA, which is used by ribosomes for protein assembly, were also downregulated in the mutants (aminoacyl-tRNA biosynthesis [KEGG:ath00970]).

Lastly, genes in the pathways that interconvert alpha-D-glucose-1P, pyruvate, and acetyl-CoA were downregulated in both *sea4* mutants (glycolysis/gluconeogenesis [KEGG:ath00010]; pentose phosphate pathway [KEGG:ath00030]; pyruvate metabolism [KEGG:ath00620]). In *sea4-1*, the pathway responsible for interconverting alpha-D-glucose-1P and starch was also downregulated (starch and sucrose metabolism [KEGG:ath00500]). In addition, fatty acid metabolism was downregulated in the *sea4* mutants (fatty acid metabolism [KEGG:ath01212]). In *sea4-1*, some genes in the pathway involved in the transformation of acetyl-CoA into malonyl-[acp], as well as most genes related to the elongation pathway, were downregulated (fatty acid biosynthesis [KEGG:ath00061]). In *sea4-2*, genes involved in the biosynthesis of long-chain fatty acids, which are essential for cutin and wax biosynthesis, as well as most genes of the pathway that produces biotin from malonyl-[acp], were also downregulated in the mutant (fatty acid elongation [KEGG:ath00062]; biotin metabolism [KEGG:ath00780]).

### 3.8. The *sea4* mutants exhibit altered auxin accumulation but normal PIN1 distribution

In our RNA-seq analysis, both *sea4* mutants exhibited upregulated genes related to the biosynthesis of indole-3-acetic acid. As the production and distribution of this phytohormone contribute to leaf shape in Arabidopsis (Bilsborough et al., 2011), we studied its spatial distribution in the *sea4* mutants. We crossed *sea4-1* and *sea4-2* plants to the *PIN1_pro_:PIN1:GFP* transgenic line (Xu et al., 2006), which serves as a reporter for the PIN-FORMED1 (PIN1) auxin efflux carrier. Additionally, we crossed the *sea4* mutants to the *DR5rev_pro_:GFP* transgenic line (Friml et al., 2003), harboring *GFP* driven by the auxin-responsive synthetic promoter *DR5rev*.

In Col-0 leaf primordia, the *PIN1_pro_:PIN1:GFP* signal was detected in the basal region where protrusions were forming (Figure S15A-D). In the *sea4* mutants, the spatial distribution of the *PIN1_pro_:PIN1:GFP* signal was similar but more confined to the primordial margins. The signal intensity was also weaker, particularly in *sea4-1* (Figure S15E-L). No difference in PIN1 distribution in the cells forming the primordial protrusions was observed between the *sea4* mutants and the wild type (Figure S16). The *DR5rev_pro_:GFP* signal was detected at the auxin maxima that formed in the margin protrusions and at the apex of Col-0 primordia, as well as in developing veins (Figure 7A-D). However, while the *DR5rev_pro_:GFP* signal showed the same spatial distribution in the basal region of *sea4-1* primordia as in Col-0, in the apical region of the primordia, the signal was present throughout the margin (Figure 7E-H). This change was even more pronounced in *sea4-2* primordia, with the *DR5rev_pro_:GFP* signal detected in some areas of the margin in the basal region of the primordia and in all epidermal cells of the apical region (Figure 7I-L), indicating substantial auxin accumulation. The distribution of auxin within the cells also appeared to differ. While in Col-0, the *DR5rev_pro_:GFP* signal was relatively uniform in the cytoplasm (Figure S17A), in the *sea4* mutants, this signal was only detected near the cytoplasmic membrane (Figure S17B-C).

**Figure 7.**
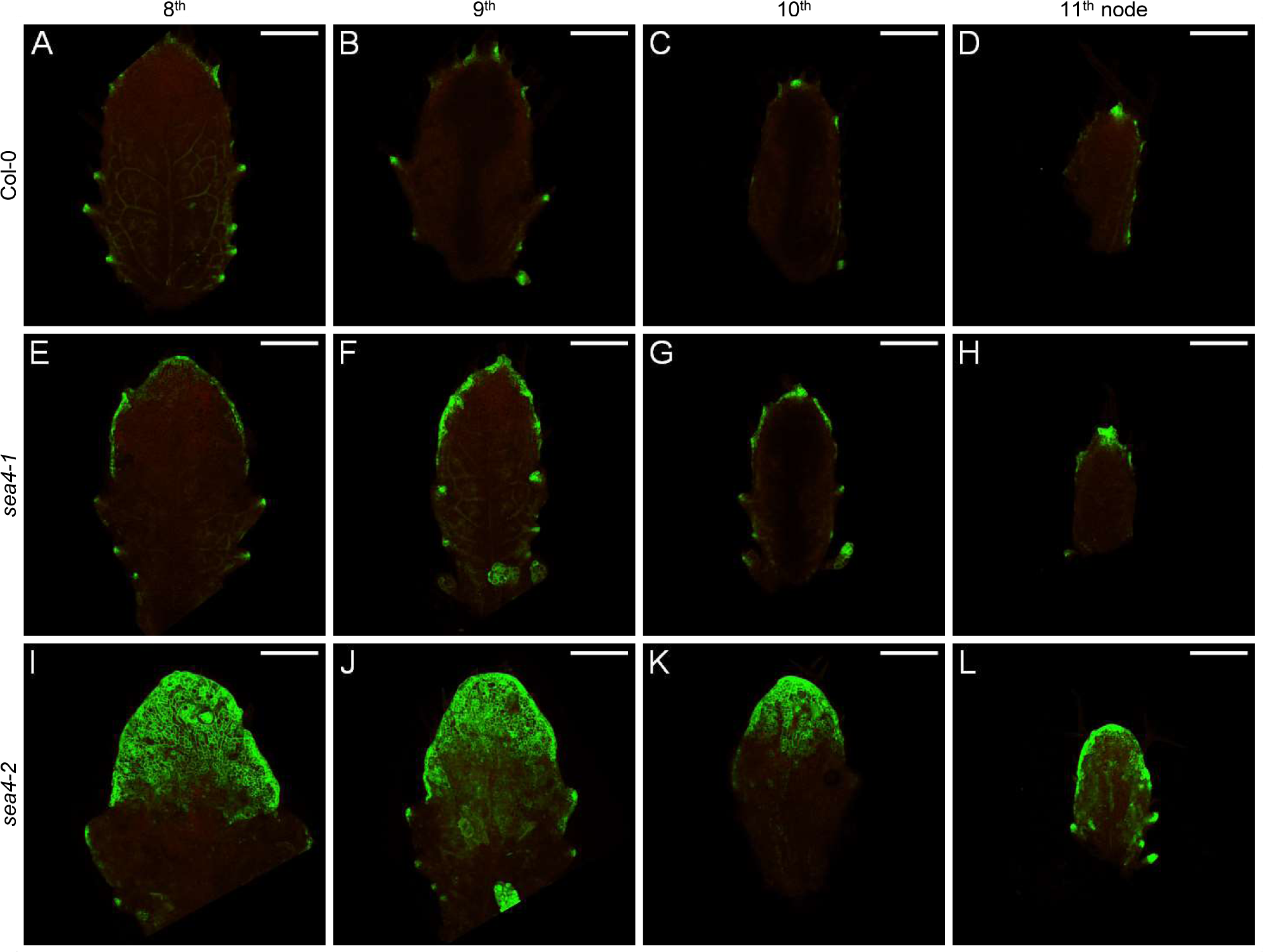
Expression pattern of the *DR5rev_pro_:GFP* reporter in leaf primordia of the *sea4* mutants. The visualization of *DR5rev_pro_:GFP* (green) expression in leaf primordia from successive nodes (8^th^-11^th^) is shown for (A-D) Col-0, (E-H) *sea4-1*, and (I-L) *sea4-2* plants. Chlorophyll autofluorescence is shown in red. The primordia were collected 20 das. Scale bars: 0.2 mm.

As primary root length is reduced in the *sea4* mutants, we also studied the spatial distribution of auxin in the apical root meristem using the *PIN1_pro_:PIN1:GFP* and *DR5rev_pro_:GFP* transgenic lines. The *PIN1_pro_:PIN1:GFP* signal localized to the center of the root in both Col-0 and the *sea4* mutants, but its intensity was weaker in the mutants, particularly in *sea4-1* (Figure S18). The distribution of PIN1 along the cytoplasmic membrane of the root cells was similar in the *sea4* mutants and Col-0. However, the cell divisions in the *sea4* mutants were poorly oriented, as evidenced by the *PIN1_pro_:PIN1:GFP* signal, which was more apparent in *sea4-1* root cells (Figure S19). The accumulation of auxin, as indicated by the *DR5rev_pro_:GFP* signal, exhibited the same spatial distribution in the *sea4* mutants as in Col-0. However, the intensity of the signal was lower in the mutants, suggesting that auxin levels were reduced at the root tip (Figure S20 and Figure S21).

## 4. DISCUSSION

### 4.1. *sea4-1* and *sea4-2* are the first viable alleles of *KEU*

Vesicle fusion is a key process in plant cytokinesis. The accurate fusion of vesicles in the appropriate orientation ultimately determines the capacity of plant tissues to grow and differentiate correctly. The presence of the KEU protein at the division plane enables the formation of trans-SNARE complexes among adjacent vesicles, resulting in the formation of the cellular plate (Park et al., 2012). Without the activity of these complexes, vesicles fuse randomly, leading to improperly oriented cytokinesis or even preventing cytokinesis altogether (Assaad et al., 1996). This is why all previously identified null *keu* mutants suffer failed cytokinesis and seedling lethality. In this study, we investigated the *sea4-1* and *sea4-2* mutants, the first known hypomorphic alleles of *KEU*, which allowed us to explore the effects of *KEU* gene alterations in adult plants.

The T-DNA insertional lines we studied, in which the *KEU* gene is interrupted at the 17^th^ intron (*keu21*), or 18^th^ exon (*keu22* and *keu23*), and the *keu^MM125^* mutant, which has a 100-bp deletion in the 16^th^ exon, are homozygous lethal. We would expect the same outcome for the *sea4-1* mutant allele, because its transcript encodes a SEA4-1 protein that is even shorter. The viability of the *sea4-1* mutant allele likely results from the combination of two factors: the small percentage of mRNA transcripts that are correctly spliced at the 9^th^ intron and its overexpression. Together, these factors result in the expression of normal mRNA in *sea4-1* at approximately 4% of the level found in wild-type Col-0. This limited expression appears to be sufficient to allow vesicle fusion at the division plane during cytokinesis, allowing the plant to complete its lifecycle.

The viability of *sea4-2* mutant allele can be explained by the observation that the S57L substitution that causes does not significantly alter the structure of KEU. This mutation reduces the flexibility of the protein and replaces only 1 of the 41 amino acids known to physically interact with syntaxins (Misura et al., 2000). Altogether, the *sea4-2* mutation appears to reduce the capacity of KEU to interact with syntaxins and form trans-SNARE complexes between adjacent vesicles, but it does not entirely disrupt its function.

### 4.2. In the *sea4* mutants, defects in cytokinesis reduce cell wall integrity and activate the unfolded protein response

The viability of our hypomorphic alleles of *KEU* allowed us to investigate their genetic interactions with related genes: those encoding its homolog SEC1B, the exocyst component SEC6, and the syntaxins SYP21, SYP132 and KN. Except for *KN*, the T-DNA insertional lines for the other genes were phenotypically wild type when homozygous. However, all their double mutant combinations with *sea4* mutations exhibited a synergistic phenotype that was quite similar: plants with fully expanded cotyledons but leaves that failed to develop correctly. These leaves produced small protuberances emerging from the shoot apical meristem and tiny, highly serrated leaves. By contrast, the *sea4/sea4;KN/kn^X37-2^*sesquimutants exhibited leaves that, while undulated and serrated, were more developed than *KN/kn^X37-2^* leaves.

Previous studies have indicated that mutants in *KEU* display some degree of polyploidy at the seedling stage (Assaad et al., 1996). We confirmed that this trait persists in adult *sea4* plants, with *sea4-1* showing more pronounced polyploidy. Higher ploidy levels typically correlate with altered levels of gene expression (Bomblies and Madlung, 2014), which we also observed in the *sea4* mutants. However, this alteration does not occur randomly: the *sea4* mutants exhibited the typical alterations in gene expression observed in mutants with reduced cell wall integrity (CWI; Gigli-Bisceglia et al., 2018). In such mutants, plant immunity is activated (Bacete et al., 2018). Indeed, the *sea4* mutants showed activation of PAMP- and effector-triggered immunity, increased activity of MAPK signaling pathways, and an increase in the biosynthesis of glucosinolate defense compounds. Other effects of changes in CWI include an increase in jasmonate (Ellis et al., 2002) and lignin biosynthesis (Caño-Delgado et al., 2003), as also observed in the *sea4* mutants. The main pathway through which CWI regulates cell division is via nitric oxide (NO) production, a process that triggers the degradation of cytokinin (Gigli-Bisceglia et al., 2018). NO also plays a crucial role in preserving auxin sensitivity in Arabidopsis root tips (Sanz et al., 2014). Mutants affected in genes involved in NO biosynthesis, such as *NITRIC OXIDE SYNTHASE1* (*NOA1*), *NITRATE REDUCTASE1* (*NIA1*) and *NIA2*, exhibit shorter roots than the wild type. In these mutants, the distribution of PIN1 is similar to that of the wild type, but the expression of *DR5_pro_:GUS* is reduced (Sanz et al., 2014). In the *sea4* mutants, *NOA1*, *NIA1*, and *NIA2* were downregulated, resulting in a similar root phenotype. Notably, *sea4* primordia maintained the same PIN1 distribution as Col-0, but *DR5rev_pro_:GFP* expression significantly increased. This difference may be attributed to the strong upregulation of the metabolic pathway responsible for synthesizing indole-3-acetic acid in the *sea4* mutants. This upregulation could potentially compensate for the reduced auxin sensitivity caused by the deficit in NO biosynthesis.

Defects in cytokinesis have been shown to induce the UPR, which in turn increases the activity of the ER to overcome these defects (Bicknell et al., 2007). One of the principal proteins initiating the UPR pathway is INCREASED ORGAN REGENERATION1 (IRE1); our RNA-seq data show that *IRE1* was upregulated in the *sea4* mutants. An increase in ER activity was also observed, with higher activities of enzymes that glycosylate proteins, transport them within the ER, and fold proteins. Genes encoding ubiquitination enzymes for misfolded proteins and the proteasome were also upregulated in the mutants, which reinforces the hypothesis of UPR activation. A possible explanation for the increase in proteolysis is that the cell detects the accumulation of vesicles and proteins in the division plane and increases the degradation of newly synthesized proteins related to cytokinesis. This explanation would also account for the reduced metabolism of some amino acids, aminoacyl-tRNA biosynthesis, and biosynthesis and elongation of fatty acids, as all these components are necessary for cytokinesis and may be in excess.

Photosynthetic light-harvesting and carbon fixation were also significantly affected in the *sea4* mutants. Numerous genes encoding proteins of LHCI and II, photosystems I and II, the F-type ATPase, and the electron transport chain were downregulated in the mutants. Additionally, enzymes involved in the metabolic pathways that synthesize chlorophyll A and B and alpha- and beta-carotene were downregulated, as were enzymes responsible for carbon fixation in the Calvin cycle. The relationship between defects in cytokinesis or reduced CWI and the decrease in photosynthetic processes has not been previously observed. Perhaps due to their extended duration of cytokinesis, *sea4* cells remain in the M phase of the cell cycle for a longer period, leading to a reduced requirement for carbohydrate compounds.

### 4.3. Other phenotypic effects of the altered cytokinesis caused by the *sea4* mutations

The phenotypes observed in the *sea4* mutants seem to be a direct consequence of their molecular alterations. The *sea4* mutations lead to defects in cytokinesis with some degree of variability, a phenomenon previously observed in other *keu* mutants (Assaad et al., 1996). This variability is visible at the tissue level, where palisade mesophyll cells vary in size, and epidermal cells cannot reach the same degree of shape complexity as in the wild type. Cells of improper size disrupt tissue organization, which is also evident in the spongy mesophyll, causing non-uniform leaf thickness and resulting in bulges. This effect can also be observed in petals, leading to premature flower opening. Disruption of internal tissue organization may also affect the boundary between dorsal and ventral leaf tissues, influencing the production and distribution of mobile signals that shape lamina growth (Caggiano et al., 2017). However, it is no clear that alterations in the cell division plane are responsible for the serrations in *sea4* leaves. The Arabidopsis *lng1-1D* mutant of *TRM1* have leaves with serrated margins (Lee et al., 2006), but in the maize (*Zea mays*) *tan-1* mutant, there is no direct correlation between the orientation of cell divisions and the final shape of the leaf (Smith et al., 1996). This is because cell divisions take place after cell elongation, which primarily determines leaf shape. Nonetheless, alterations in the cell division plane could be the cause of the reduced leaf size in the *sea4* mutants, a trait that was also observed in the *tan-1* mutant.

Although the distribution of the polar auxin transporter PIN1 remains unaltered in *sea4* leaves, the upregulation of genes in the auxin biosynthetic pathway leads to the excess accumulation of auxin, which diffuses into regions where it should not be. This ectopic auxin distribution appears to be responsible for the serration observed in *sea4* leaves. Moreover, the presence of auxin maxima during primordia development has been linked to vascular tissue development (Aloni et al., 2003; Teale et al., 2006). The altered auxin maxima in the *sea4* mutants likely accounts for their denser and more complex vascular pattern. Reduced sensitivity to auxins in the root apical meristem via NO appears to be the reason for reduced root length. This phenomenon likely also occurs in the shoot apical meristem, explaining the observed reduction in plant height.

Leaf senescence is another characteristic trait of the *sea4* mutants. Genes related to plant immunity and MAPK signaling pathways are upregulated in these mutants due to a reduction in CWI. This activation triggers defense responses and, in some cases, cell death. Consequently, regions of the leaves with a high number of dead cells may form the senescent patches observed in these mutants. Defects in cytokinesis accumulate in each leaf over time and more mitosis events occur, explaining why younger leaves do not exhibit as many senescent patches as older leaves.

## 5. CONCLUSIONS

The SM KEU protein has a known key role in cytokinesis, coordinating the assembly of *trans*-SNARE complexes in vesicle fusions at the cell plate. Here, we studied the first hypomorphic, viable alleles of the *KEU* gene, *sea4-1* and *sea4-2*. In *sea4-1*, a transition at the splice donor site of the ninth exon leads to mis-splicing, yielding a truncated protein. In *sea4-2*, a transition in the third exon causes a S57L substitution that impacts an amino acid crucial for physical interactions with syntaxins and reduces the protein flexibility. In these viable mutants, cytokinesis is impaired but not abolished, allowing the functional study of the roles that *KEU* play in vegetative and reproductive development. Our RNA-seq study of *sea4* plants show that their phenotypes are associated to activation of the unfolded protein response and reduction in cell wall integrity. The characteristic early leaf senescence of *sea4* plants seems to be caused by the activation of plant immunity and the increase in jasmonate biosynthesis, two typical traits of mutants with reduced cell wall integrity. Their short primary shoots and roots are also a likely consequence of a reduction in auxin sensitivity at the meristems due to a decrease in NO biosynthesis, another typical trait of reduced cell wall integrity mutants. To overcome the reduced auxin sensitivity, the *sea4* mutants upregulate auxin biosynthesis, leading to its ectopic diffusion within leaf primordia, causing leaf margin serrations and a dense, complex venation pattern. The *sea4* leaf surface displays protuberances caused by internal tissue disorganization due to cytokinesis defects. Such disorganization seem to disrupt the boundaries between tissues that regulate leaf morphogenesis and may contribute to the formation of serrated leaf margins. The *sea4* mutants also exhibit a reduction in photosynthetic light-harvesting and carbon fixation, a phenomenon not previously linked to either cell wall integrity or the unfolded protein response, but that could be related to the delay in cytokinesis due to the *sea4* mutations.

## Supporting information

Data Set S1

Data Set S2

Data Set S3

## AUTHOR CONTRIBUTIONS

J.L.M. conceived, designed, and supervised the research, provided resources, and obtained funding. Several experiments were codesigned by A.R.-B., R.S.-M., and J.L.M. A.R.-B. performed most of the experiments. RM-V, FML, and MRP performed the low-resolution mapping of *sea4-1* and *sea4-2*. CC-R contributed to the phenotypic analysis of the *sea4* mutants. A.R.-B. and J.L.M. wrote the manuscript. All authors revised and approved the manuscript.

## ACKNOWLEDGEMENTS

The authors would like to express their gratitude to G. Jürgens for providing seeds, J. Castelló, J.M. Serrano, and M.J. Ñíguez for their excellent technical assistance, and A.M. Aguilar-García, Á. Valdés-Penalva, and E. Peñataro-González for their contributions to the phenotypic characterization of the *sea4* mutants.

## APPENDIX A. SUPPLEMENTARY MATERIAL

**Figure S1.** Mis-splicing of the *KEU* gene caused by the *sea4-1* mutation.

**Figure S2.** Sequence conservation between Arabidopsis KEU and the rat STXBP1 proteins.

**Figure S3.** Predicted effect of the S57L mutation on the dynamics of KEU protein.

**Figure S4.** Comparison of the 3D structures of wild-type and S57L mutant (SEA4-2) KEU proteins.

**Figure S5.** Phenotypic classes observed in *sea4-1* and *sea4-2* plants.

**Figure S7.** Morphological phenotypes of adult plants carrying mutant alleles of *KEU*.

**Figure S9.** Structure of the cell layers in first-node leaves of the *sea4* mutants.

**Figure S10.** Progression of leaf senescence in *sea4-1* and *sea4-2* plants.

**Figure S11.** Extent of senescence and bolting time in the *sea4* mutants.

**Figure S12.** Structures of the genes that genetically interact with *KEU*, indicating the nature and positions of their mutations.

**Figure S13.** Nuclear ploidy levels in *sea4* leaves.

**Figure S14.** Differentially expressed genes in the *sea4* mutants identified by RNA-seq.

**Figure S15.** Expression pattern of the *PIN1_pro_:PIN1:GFP* reporter in leaf primordia of the *sea4* mutants.

**Figure S16.** Localization of the *PIN1_pro_:PIN1:GFP* reporter in the cell membrane in leaf primordia of the *sea4* mutants.

**Figure S17.** Localization of the *DR5rev_pro_:GFP* reporter in the cell membrane in leaf primordia of the *sea4* mutants.

**Figure S18.** Expression pattern of the *PIN1_pro_:PIN1:GFP* reporter in the root apex of the *sea4* mutants.

**Figure S19.** Localization of the *PIN1_pro_:PIN1:GFP* reporter in the cell membrane in roots of the *sea4* mutants.

**Figure S20.** Expression pattern of the *DR5rev_pro_:GFP* reporter in the root apex of the *sea4* mutants.

**Figure S21.** Localization of the *DR5rev_pro_:GFP* reporter in the cell membrane in roots of the *sea4* mutants.

**Table S1.** Primer sets used in this work.

**Table S2.** Configuration parameters of the Leica Stellaris 8 STED confocal microscope.

**Table S3.** Data quality control summary of the RNA-seq assay.

**Table S4.** Candidate mutations identified in *sea4-1* by Easymap.

**Table S5.** Candidate mutations identified in *sea4-2* by Easymap.

**Table S6.** Retention of the 9^th^ intron of *KEU* in the RNA-seq assay.

**Table S7.** Expression of *KEU* in the RNA-seq assay.

**Table S8.** Predicted effects of the S57L mutations on the stability and dynamics of the KEU protein.

**Table S9.** Phenotypic variability in the progeny of selfed *sea4* plants.

**Table S10.** Venation pattern characteristics of *sea4* cotyledons and leaves.

**Table S11.** Nuclear DNA ploidy distribution (%) in the first pair of leaves of the *sea4* mutants.

## REFERENCES

R. Aloni, K. Schwalm, M. Langhans, and C.I. Ullrich, Gradual shifts in sites of free-auxin production during leaf-primordium development and their role in vascular differentiation and leaf morphogenesis in Arabidopsis, Planta 216 (2003) 841–853, 10.1007/s00425-002-0937-8.

F.F. Assaad, Y. Huet, U. Mayer, and G. Jürgens, The cytokinesis gene *KEULE* encodes a Sec1 protein that binds the syntaxin KNOLLE, The Journal of cell biology 152 (2001) 531–543, 10.1083/jcb.152.3.531.

F.F. Assaad, U. Mayer, G. Wanner, and G. Jürgens, The *KEULE* gene is involved in cytokinesis in *Arabidopsis*, Molecular & general genetics : MGG 253 (1996) 267–277, 10.1007/pl00008594.

L. Bacete, H. Mélida, E. Miedes, and A. Molina, Plant cell wall-mediated immunity: cell wall changes trigger disease resistance responses, The Plant journal : for cell and molecular biology 93 (2018) 614–636, 10.1111/tpj.13807.

G. Berná, P. Robles, and J.L. Micol, A mutational analysis of leaf morphogenesis in *Arabidopsis thaliana*, Genetics 152 (1999) 729–742, 10.1093/genetics/152.2.729.

A.A. Bicknell, A. Babour, C.M. Federovitch, and M. Niwa, A novel role in cytokinesis reveals a housekeeping function for the unfolded protein response, The Journal of cell biology 177 (2007) 1017–1027, 10.1083/jcb.200702101.

G.D. Bilsborough, A. Runions, M. Barkoulas, H.W. Jenkins, A. Hasson, C. Galinha, P. Laufs, A. Hay, P. Prusinkiewicz, and M. Tsiantis, Model for the regulation of *Arabidopsis thaliana* leaf margin development, Proceedings of the National Academy of Sciences of the United States of America 108 (2011) 3424–3429, 10.1073/pnas.1015162108.

M.R. Blatt, B. Leyman, and D. Geelen, Tansley review No. 108: molecular events of vesicle trafficking and control by SNARE proteins in plants, The New phytologist 144 (1999) 389–418, 10.1046/j.1469-8137.1999.00546.x.

K. Bomblies, and A. Madlung, Polyploidy in the Arabidopsis genus, Chromosome Res. 22 (2014) 117–134, 10.1007/s10577-014-9416-x.

J. Bühler, L. Rishmawi, D. Pflugfelder, G. Huber, H. Scharr, M. Hülskamp, M. Koornneef, U. Schurr, and S. Jahnke, phenoVein―A tool for leaf vein segmentation and analysis, Plant physiology 169 (2015) 2359–2370, 10.1104/pp.15.00974.

M.P. Caggiano, X. Yu, N. Bhatia, A. Larsson, H. Ram, C.K. Ohno, P. Sappl, E.M. Meyerowitz, H. Jönsson, and M.G. Heisler, Cell type boundaries organize plant development, eLife 6 (2017) e27421, 10.7554/eLife.27421.

H. Candela, A. Martínez-Laborda, and J.L. Micol, Venation pattern formation in *Arabidopsis thaliana* vegetative leaves, Dev. Biol. 205 (1999) 205–216, 10.1006/dbio.1998.9111.

A. Caño-Delgado, S. Penfield, C. Smith, M. Catley, and M. Bevan, Reduced cellulose synthesis invokes lignification and defense responses in *Arabidopsis thaliana*, The Plant journal : for cell and molecular biology 34 (2003) 351–362, 10.1046/j.1365-313x.2003.01729.x.

B. Desvoyes, E. Ramirez-Parra, Q. Xie, N.H. Chua, and C. Gutierrez, Cell type-specific role of the retinoblastoma/E2F pathway during Arabidopsis leaf development, Plant physiology 140 (2006) 67–80, 10.1104/pp.105.071027.

S. Drevensek, M. Goussot, Y. Duroc, A. Christodoulidou, S. Steyaert, E. Schaefer, E. Duvernois, O. Grandjean, M. Vantard, D. Bouchez, and M. Pastuglia, The *Arabidopsis* TRM1-TON1 interaction reveals a recruitment network common to plant cortical microtubule arrays and eukaryotic centrosomes, The Plant cell 24 (2012) 178–191, 10.1105/tpc.111.089748.

C. Ellis, I. Karafyllidis, C. Wasternack, and J.G. Turner, The Arabidopsis mutant *cev1* links cell wall signaling to jasmonate and ethylene responses, The Plant cell 14 (2002) 1557–1566, 10.1105/tpc.002022.

M. Fagard, T. Desnos, T. Desprez, F. Goubet, G. Refregier, G. Mouille, M. Mccann, C. Rayon, S. Vernhettes, and H. Höfte, *PROCUSTE1* encodes a cellulose synthase required for normal cell elongation specifically in roots and dark-grown hypocotyls of Arabidopsis, The Plant cell 12 (2000) 2409–2423, 10.1105/tpc.12.12.2409.

V. Frappier, M. Chartier, and R.J. Najmanovich, ENCoM server: exploring protein conformational space and the effect of mutations on protein function and stability, Nucleic Acids Res. 43 (2015) W395–400, 10.1093/nar/gkv343.

J. Friml, A. Vieten, M. Sauer, D. Weijers, H. Schwarz, T. Hamann, R. Offringa, and G. Jürgens, Efflux-dependent auxin gradients establish the apical-basal axis of *Arabidopsis*, Nature 426 (2003) 147–153, 10.1038/nature02085.

R. Fukuda, J.A. Mcnew, T. Weber, F. Parlati, T. Engel, W. Nickel, J.E. Rothman, and T.H. Söllner, Functional architecture of an intracellular membrane t-SNARE, Nature 407 (2000) 198–202, 10.1038/35025084.

D.W. Galbraith, K.R. Harkins, and S. Knapp, Systemic endopolyploidy in *Arabidopsis thaliana*, Plant physiology 96 (1991) 985–989, 10.1104/pp.96.3.985.

N. Gigli-Bisceglia, T. Engelsdorf, M. Strnad, L. Vaahtera, G.A. Khan, A. Yamoune, L. Alipanah, O. Novák, S. Persson, J. Hejatko, and T. Hamann, Cell wall integrity modulates *Arabidopsis thaliana* cell cycle gene expression in a cytokinin- and nitrate reductase-dependent manner, Development 145 (2018) dev166678, 10.1242/dev.166678.

T.D. Goddard, C.C. Huang, E.C. Meng, E.F. Pettersen, G.S. Couch, J.H. Morris, and T.E. Ferrin, UCSF ChimeraX: meeting modern challenges in visualization and analysis, Protein Sci. 27 (2018) 14–25, 10.1002/pro.3235.

M. Heese, X. Gansel, L. Sticher, P. Wick, M. Grebe, F. Granier, and G. Jürgens, Functional characterization of the KNOLLE-interacting t-SNARE AtSNAP33 and its role in plant cytokinesis, The Journal of cell biology 155 (2001) 239–249, 10.1083/jcb.200107126.

Z. Hong, A.J. Delauney, and D.P. Verma, A cell plate-specific callose synthase and its interaction with phragmoplastin, The Plant cell 13 (2001) 755–768, 10.1105/tpc.13.4.755.

G. Horiguchi, A. Mollá-Morales, J.M. Pérez-Pérez, K. Kojima, P. Robles, M.R. Ponce, J.L. Micol, and H. Tsukaya, Differential contributions of ribosomal protein genes to *Arabidopsis thaliana* leaf development, The Plant journal : for cell and molecular biology 65 (2011) 724–736, 10.1111/j.1365-313X.2010.04457.x.

S. Ittisoponpisan, S.A. Islam, T. Khanna, E. Alhuzimi, A. David, and M.J.E. Sternberg, Can predicted protein 3D structures provide reliable insights into whether missense variants are disease associated?, J. Mol. Biol. 431 (2019) 2197–2212, 10.1016/j.jmb.2019.04.009.

G. Jürgens, Plant cytokinesis: fission by fusion, Trends in cell biology 15 (2005) 277–283, 10.1016/j.tcb.2005.03.005.

M. Karnahl, M. Park, C. Krause, U. Hiller, U. Mayer, Y.D. Stierhof, and G. Jürgens, Functional diversification of *Arabidopsis* SEC1-related SM proteins in cytokinetic and secretory membrane fusion, Proceedings of the National Academy of Sciences of the United States of America 115 (2018) 6309–6314, 10.1073/pnas.1722611115.

R. Karnik, C. Grefen, R. Bayne, A. Honsbein, T. Köhler, D. Kioumourtzoglou, M. Williams, N.J. Bryant, and M.R. Blatt, *Arabidopsis* Sec1/Munc18 protein SEC11 is a competitive and dynamic modulator of SNARE binding and SYP121-dependent vesicle traffic, The Plant cell 25 (2013) 1368–1382, 10.1105/tpc.112.108506.

D. Kim, J.M. Paggi, C. Park, C. Bennett, and S.L. Salzberg, Graph-based genome alignment and genotyping with HISAT2 and HISAT-genotype, Nat. Biotechnol. 37 (2019) 907–915, 10.1038/s41587-019-0201-4.

T. Krupnova, M. Sasabe, L. Ghebreghiorghis, C.W. Gruber, T. Hamada, V. Dehmel, G. Strompen, Y.D. Stierhof, W. Lukowitz, B. Kemmerling, Y. Machida, T. Hashimoto, U. Mayer, and G. Jürgens, Microtubule-associated kinase-like protein RUNKEL needed for cell plate expansion in *Arabidopsis* cytokinesis, Current biology : CB 19 (2009) 518–523, 10.1016/j.cub.2009.02.021.

Y.K. Lee, G.T. Kim, I.J. Kim, J. Park, S.S. Kwak, G. Choi, and W.I. Chung, *LONGIFOLIA1* and *LONGIFOLIA2*, two homologous genes, regulate longitudinal cell elongation in *Arabidopsis*, Development 133 (2006) 4305–4314, 10.1242/dev.02604.

M.I. Love, W. Huber, and S. Anders, Moderated estimation of fold change and dispersion for RNA-seq data with DESeq2, Genome Biol. 15 (2014) 550, 10.1186/s13059-014-0550-8.

W. Lukowitz, U. Mayer, and G. Jürgens, Cytokinesis in the Arabidopsis embryo involves the syntaxin-related KNOLLE gene product, Cell 84 (1996) 61–71, 10.1016/s0092-8674(00)80993-9.

S.D. Lup, C. Navarro-Quiles, and J.L. Micol, Versatile mapping-by-sequencing with Easymap v.2, Front. Plant Sci. 14 (2023) 1042913, 10.3389/fpls.2023.1042913.

S.D. Lup, D. Wilson-Sánchez, S. Andreu-Sánchez, and J.L. Micol, Easymap: a user-friendly software package for rapid mapping-by-sequencing of point mutations and large insertions, Front. Plant Sci. 12 (2021) 655286, 10.3389/fpls.2021.655286.

S.D. Lup, D. Wilson-Sánchez, and J.L. Micol, Mapping-by-sequencing of point and insertional mutations with Easymap, Methods. Mol. Biol. 2484 (2022) 343–361, 10.1007/978-1-0716-2253-7_23.

V. Lupashin, and E. Sztul, Golgi tethering factors, Biochim. Biophys. Acta 1744 (2005) 325–339, 10.1016/j.bbamcr.2005.03.013.

F. Madeira, Y.M. Park, J. Lee, N. Buso, T. Gur, N. Madhusoodanan, P. Basutkar, A.R.N. Tivey, S.C. Potter, R.D. Finn, and R. Lopez, The EMBL-EBI search and sequence analysis tools APIs in 2019, Nucleic Acids Res. 47 (2019) W636–W641, 10.1093/nar/gkz268.

M. Margittai, J. Widengren, E. Schweinberger, G.F. Schröder, S. Felekyan, E. Haustein, M. König, D. Fasshauer, H. Grubmüller, R. Jahn, and C.a.M. Seidel, Single-molecule fluorescence resonance energy transfer reveals a dynamic equilibrium between closed and open conformations of syntaxin 1, Proceedings of the National Academy of Sciences of the United States of America 100 (2003) 15516–15521, 10.1073/pnas.2331232100.

K.M.S. Misura, R.H. Scheller, and W.I. Weis, Three-dimensional structure of the neuronal-Sec1–syntaxin 1a complex, Nature 404 (2000) 355–362, 10.1038/35006120.

S. Müller, E. Fuchs, M. Ovecka, J. Wysocka-Diller, P.N. Benfey, and M.T. Hauser, Two new loci, *PLEIADE* and *HYADE*, implicate organ-specific regulation of cytokinesis in Arabidopsis, Plant physiology 130 (2002) 312–324, 10.1104/pp.004416.

S. Müller, and G. Jürgens, Plant cytokinesis―No ring, no constriction but centrifugal construction of the partitioning membrane, Seminars in cell & developmental biology 53 (2016) 10–18, 10.1016/j.semcdb.2015.10.037.

C. Navarro-Quiles, E. Mateo-Bonmatí, H. Candela, P. Robles, A. Martínez-Laborda, Y. Fernández, J. Simura, K. Ljung, V. Rubio, M.R. Ponce, and J.L. Micol, The Arabidopsis ATP-Binding Cassette E protein ABCE2 is a conserved component of the translation machinery, Front. Plant Sci. 13 (2022) 1009895, 10.3389/fpls.2022.1009895.

T.C. Nickle, and D.W. Meinke, A cytokinesis-defective mutant of *Arabidopsis* (*cyt1*) characterized by embryonic lethality, incomplete cell walls, and excessive callose accumulation, The Plant journal : for cell and molecular biology 15 (1998) 321–332, 10.1046/j.1365-313x.1998.00212.x.

M. Park, S. Touihri, I. Müller, U. Mayer, and G. Jürgens, Sec1/Munc18 protein stabilizes fusion-competent syntaxin for membrane fusion in *Arabidopsis* cytokinesis, Developmental cell 22 (2012) 989–1000, 10.1016/j.devcel.2012.03.002.

E.F. Pettersen, T.D. Goddard, C.C. Huang, E.C. Meng, G.S. Couch, T.I. Croll, J.H. Morris, and T.E. Ferrin, UCSF ChimeraX: structure visualization for researchers, educators, and developers, Protein Sci. 30 (2021) 70–82, 10.1002/pro.3943.

D.E.V. Pires, D.B. Ascher, and T.L. Blundell, DUET: a server for predicting effects of mutations on protein stability using an integrated computational approach, Nucleic Acids Res. 42 (2014a) W314–319, 10.1093/nar/gku411.

D.E.V. Pires, D.B. Ascher, and T.L. Blundell, mCSM: predicting the effects of mutations in proteins using graph-based signatures, Bioinformatics 30 (2014b) 335–342, 10.1093/bioinformatics/btt691.

M.R. Ponce, V. Quesada, and J.L. Micol, Rapid discrimination of sequences flanking and within T-DNA insertions in the *Arabidopsis* genome, The Plant journal : for cell and molecular biology 14 (1998) 497–501, 10.1046/j.1365-313x.1998.00146.x.

M.R. Ponce, P. Robles, F.M. Lozano, M.A. Brotóns, and J.L. Micol, Low-resolution mapping of untagged mutations, Methods. Mol. Biol. 323 (2006) 105–113, 10.1385/1-59745-003-0:105.

M.R. Ponce, P. Robles, and J.L. Micol, High-throughput genetic mapping in *Arabidopsis thaliana*, Molecular & general genetics : MGG 261 (1999) 408–415, 10.1007/s004380050982.

V. Quesada, M.R. Ponce, and J.L. Micol, Genetic analysis of salt-tolerant mutants in *Arabidopsis thaliana*, Genetics 154 (2000) 421–436, 10.1093/genetics/154.1.421.

P. Robles, and J.L. Micol, Genome-wide linkage analysis of *Arabidopsis* genes required for leaf development, Mol. Genet. Genomics 266 (2001) 12–19, 10.1007/s004380100535.

C.H.M. Rodrigues, D.E.V. Pires, and D.B. Ascher, DynaMut: predicting the impact of mutations on protein conformation, flexibility and stability, Nucleic Acids Res. 46 (2018) W350–W355, 10.1093/nar/gky300.

C.H.M. Rodrigues, D.E.V. Pires, and D.B. Ascher, DynaMut2: assessing changes in stability and flexibility upon single and multiple point missense mutations, Protein Sci. 30 (2021) 60–69, 10.1002/pro.3942.

L. Sanz, M. Fernández-Marcos, A. Modrego, D.R. Lewis, G.K. Muday, S. Pollmann, M. Dueñas, C. Santos-Buelga, and O. Lorenzo, Nitric oxide plays a role in stem cell niche homeostasis through its interaction with auxin, Plant physiology 166 (2014) 1972–1984, 10.1104/pp.114.247445.

J. Serrano-Cartagena, H. Candela, P. Robles, M.R. Ponce, J.M. Pérez-Pérez, P. Piqueras, and J.L. Micol, Genetic analysis of *incurvata* mutants reveals three independent genetic operations at work in Arabidopsis leaf morphogenesis, Genetics 156 (2000) 1363–1377, 10.1093/genetics/156.3.1363.

L.G. Smith, S. Hake, and A.W. Sylvester, The *tangled-1* mutation alters cell division orientations throughout maize leaf development without altering leaf shape, Development 122 (1996) 481–489, 10.1242/dev.122.2.481.

R. Söllner, G. Glässer, G. Wanner, C.R. Somerville, G. Jürgens, and F.F. Assaad, Cytokinesis-defective mutants of Arabidopsis, Plant physiology 129 (2002) 678–690, 10.1104/pp.004184.

K. Steinborn, C. Maulbetsch, B. Priester, S. Trautmann, T. Pacher, B. Geiges, F. Küttner, L. Lepiniec, Y.D. Stierhof, H. Schwarz, G. Jürgens, and U. Mayer, The Arabidopsis *PILZ* group genes encode tubulin-folding cofactor orthologs required for cell division but not cell growth, Genes & development 16 (2002) 959–971, 10.1101/gad.221702.

G. Strompen, F. El Kasmi, S. Richter, W. Lukowitz, F.F. Assaad, G. Jürgens, and U. Mayer, The Arabidopsis *HINKEL* gene encodes a kinesin-related protein involved in cytokinesis and is expressed in a cell cycle-dependent manner, Current biology : CB 12 (2002) 153–158, 10.1016/s0960-9822(01)00655-8.

S. Struk, and P. Dhonukshe, MAPs: cellular navigators for microtubule array orientations in *Arabidopsis*, Plant Cell Rep. 33 (2014) 1–21, 10.1007/s00299-013-1486-2.

W.D. Teale, I.A. Paponov, and K. Palme, Auxin in action: signalling, transport and the control of plant growth and development, Nat. Rev. Mol. Cell Biol. 7 (2006) 847–859, 10.1038/nrm2020.

M. Thellmann, K. Rybak, K. Thiele, G. Wanner, and F.F. Assaad, Tethering factors required for cytokinesis in Arabidopsis, Plant physiology 154 (2010) 720–732, 10.1104/pp.110.154286.

K. Thiele, G. Wanner, V. Kindzierski, G. Jürgens, U. Mayer, F. Pachl, and F.F. Assaad, The timely deposition of callose is essential for cytokinesis in Arabidopsis, The Plant journal : for cell and molecular biology 58 (2009) 13–26, 10.1111/j.1365-313X.2008.03760.x.

K. Tunyasuvunakool, J. Adler, Z. Wu, T. Green, M. Zielinski, A. Žídek, A. Bridgland, A. Cowie, C. Meyer, A. Laydon, S. Velankar, G.J. Kleywegt, A. Bateman, R. Evans, A. Pritzel, M. Figurnov, O. Ronneberger, R. Bates, S.a.A. Kohl, A. Potapenko, A.J. Ballard, B. Romera-Paredes, S. Nikolov, R. Jain, E. Clancy, D. Reiman, S. Petersen, A.W. Senior, K. Kavukcuoglu, E. Birney, P. Kohli, J. Jumper, and D. Hassabis, Highly accurate protein structure prediction for the human proteome, Nature 596 (2021) 590–596, 10.1038/s41586-021-03828-1.

M. Varadi, S. Anyango, M. Deshpande, S. Nair, C. Natassia, G. Yordanova, D. Yuan, O. Stroe, G. Wood, A. Laydon, A. Žídek, T. Green, K. Tunyasuvunakool, S. Petersen, J. Jumper, E. Clancy, R. Green, A. Vora, M. Lutfi, M. Figurnov, A. Cowie, N. Hobbs, P. Kohli, G. Kleywegt, E. Birney, D. Hassabis, and S. Velankar, AlphaFold Protein Structure Database: massively expanding the structural coverage of protein-sequence space with high-accuracy models, Nucleic Acids Res. 50 (2022) D439–D444, 10.1093/nar/gkab1061.

I. Waizenegger, W. Lukowitz, F. Assaad, H. Schwarz, G. Jürgens, and U. Mayer, The *Arabidopsis KNOLLE* and *KEULE* genes interact to promote vesicle fusion during cytokinesis, Current biology : CB 10 (2000) 1371–1374, 10.1016/s0960-9822(00)00775-2.

A.M.D. Wiedemeier, J.E. Judy-March, C.H. Hocart, G.O. Wasteneys, R.E. Williamson, and T.I. Baskin, Mutant alleles of *Arabidopsis RADIALLY SWOLLEN 4* and *7* reduce growth anisotropy without altering the transverse orientation of cortical microtubules or cellulose microfibrils, Development 129 (2002) 4821–4830, 10.1242/dev.129.20.4821.

R.E. Williamson, J.E. Burn, R. Birch, T.I. Baskin, T. Arioli, A.S. Betzner, and A. Cork, Morphology of *rsw1*, a cellulose-deficient mutant of *Arabidopsis thaliana*, Protoplasma 215 (2001) 116–127, 10.1007/BF01280308.

C.L. Worth, R. Preissner, and T.L. Blundell, SDM―a server for predicting effects of mutations on protein stability and malfunction, Nucleic Acids Res. 39 (2011) W215–222, 10.1093/nar/gkr363.

J. Wu, X. Tan, C. Wu, K. Cao, Y. Li, and Y. Bao, Regulation of cytokinesis by exocyst subunit SEC6 and KEULE in *Arabidopsis thaliana*, Mol. Plant 6 (2013) 1863–1876, 10.1093/mp/sst082.

J. Xu, H. Hofhuis, R. Heidstra, M. Sauer, J. Friml, and B. Scheres, A molecular framework for plant regeneration, Science 311 (2006) 385–388, 10.1126/science.1121790.

L. Zapata, J. Ding, E.M. Willing, B. Hartwig, D. Bezdan, W.B. Jiao, V. Patel, G. Velikkakam James, M. Koornneef, S. Ossowski, and K. Schneeberger, Chromosome-level assembly of *Arabidopsis thaliana* L*er* reveals the extent of translocation and inversion polymorphisms, Proceedings of the National Academy of Sciences of the United States of America 113 (2016) E4052–4060, 10.1073/pnas.1607532113.

J. Zuo, Q.W. Niu, N. Nishizawa, Y. Wu, B. Kost, and N.H. Chua, KORRIGAN, an Arabidopsis endo-1,4-β-glucanase, localizes to the cell plate by polarized targeting and is essential for cytokinesis, The Plant cell 12 (2000) 1137–1152, 10.1105/tpc.12.7.1137.

