## Supplementary Material for "Postembryonic developmental roles of the Arabidopsis *KEULE* gene"

### Supplementary Figures and Tables

Supplementary Material included in this file:

- Figures S1-S21
- Tables S1-S11

Supplementary Material not included in this file:

- Data Sets S1-S3

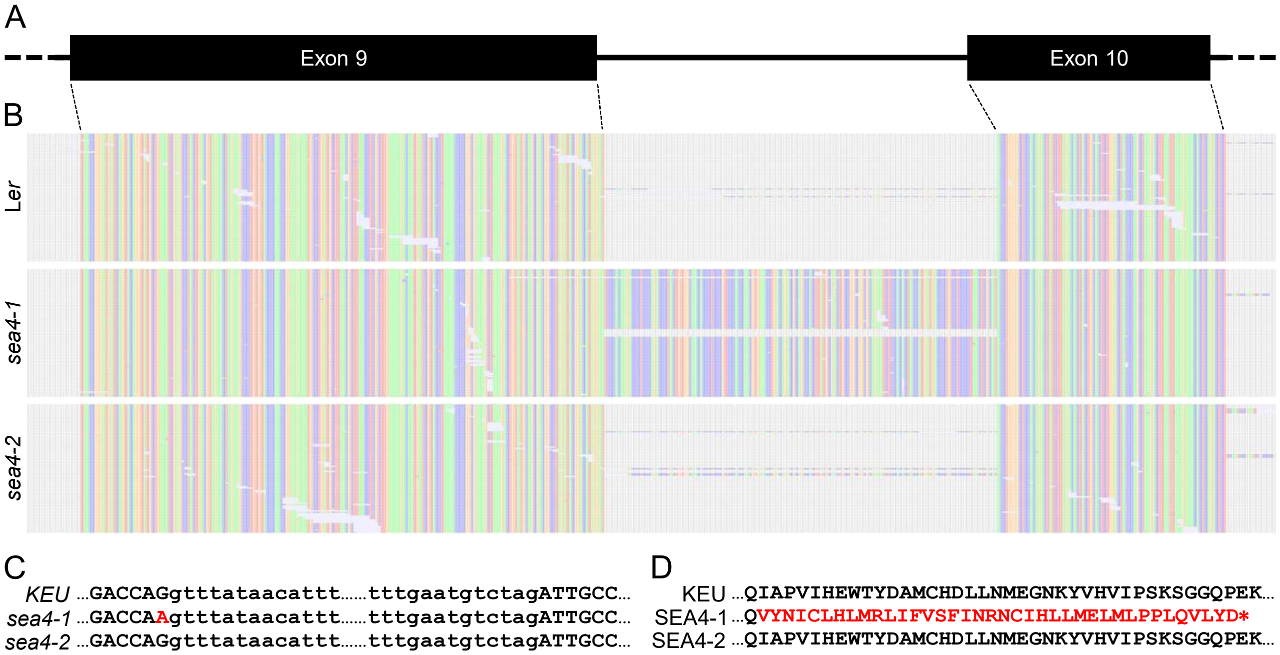


**Figure S1.** Mis-splicing of the *KEU* gene caused by the *sea4-1* mutation. (A) Schematic representation of the 9th and 10th exons of the *KEU* gene. (B) Alignment of L*er*, *sea4-1*, and *sea4-2* cDNA reads from the *KEU* gene of the Col-0 reference genome. The adenine, cytosine, thymine, and guanine bases are represented as green, orange, blue, and red squares, respectively. Positions without aligned bases are represented as gray squares. (C) Alignment of the DNA sequences of the *KEU* wild-type allele with the *sea4-1* and *sea4-2* mutant alleles. Uppercase and lowercase letters indicate exons and introns, respectively. The point mutation of *sea4-1* is indicated in red. (D) Predicted amino acid sequence of the KEU wild-type protein and the SEA4-1 and SEA4-2 mutant proteins. Intron retention in *sea4-1* generates a new amino acid sequence and a premature stop codon, both shown in red.

A

```
STXBP1       1 ------------MAPIGLKAVVGEKIMHDV---IKKVKKKGEWKVLVVDQLSMRMLSSCCKMTDIMTEGITIVEDINKRREPLPS
KEULE        1 MSYSDSDSSSHGGEYKNFRQITRERLLYEMLRSAKTGSSKSTWKVLIMDKLTVKIMSYACKMADITQEGVSLVEDIFRRRQPLPS
Consensus    1                 .:: :. *::::::    *. ..*. ****::*:*:::::* .***:**  **:::**** :**:****

STXBP1      71 LEAVYLITPSEKSVHSLISDFKDPPTAKYRAAHVFFTDSCPDALFNELVKS-RAAKVIKTLTEINIAFLPYESQVYSLDSADSFQ
KEULE       86 MDAIYFIQPTKENVIMFLSDMSGK-SPLYKKAFVFFSSPVSKELVGHIKKDSSVLPRIGALREMNLEFFAIDSQGFITDHERALE
Consensus   86 ::*:*:* *:::.*  ::**:..  :  *: *.***:.   . *...: *.  .   * :* *:*: *:  :** : *   :::

STXBP1     155 SFYSPHKA-QMKNPILERLAEQIATLCATLKEYPAVRYRGEYKDNALLAQ----LIQDK-----LDAYKADDPTMGEGPDKARSQ
KEULE      170 DLFGDEETSRKGDACLNVMASRIATVFASLREFPAVRYRAAKSLDASTMTTLRDLIPTKLAAGIWNCLAKHKQSIENFPQTETCE
Consensus  171 .::. .:: :  :  *: :*.:***: *:*:*:******.  . :*        **  *      :.   .. :: : *:.  .:

STXBP1     230 LLILDRGFDPSSPVLHELTFQAMSYDLLPIENDVYKYE--TSGIGEARVKEVLLDEDDDLWIALRHKHIAEVSQEVTRSLKDFSS
KEULE      255 LLILDRSIDQIAPVIHEWTYDAMCHDLLNMEGNKYVHVIPSKSGGQPEKKDVLLEEHDPIWLELRHAHIADASERLHDKMTNFLS
Consensus  256 ******.:*  :**:** *::**.:*** :*.: * :   :.. *: . *:***:*.* :*: *** ***:.*:.:  .:.:* *

STXBP1     313 SKR------MNTGEKTTMRDLSQMLKKMPQYQKELSKYSTHLHLAEDCMKHY-QGTVDKLCRVEQDLAMGTDAEGEKIKDPMRAI
KEULE      340 KNKAAQLQGKRDGAELSTRDLQKMVQALPQYSEQIDKLSLHVEIARKLNDLIREQGLRELGQLEQDLVFGDAGMKDVIKYL----
Consensus  341 .::       . * : : ***.:*:: :***.:::.* * *:.:*..  .   :  : :* ::****.:*  .  : **

STXBP1     391 VPILLDANVSTYDKIRIILLYI-FLKNGITEENLNKLIQHAQIPPEDSEIITNMAHLGVPIVT----------DSTLRRRSKPER
KEULE      421 ---STQEEASREGKLRLLMILATIYPEKFEGEKGQNLMKLAKLSSDDMTAVNNMSLLGSAVDAKKNTPGGFTLKFDLHKKKRAVR
Consensus  426      : :.*  .*:*::::   :  :  : *: ::*:: *::  :*   :.**: **  : :          .  *:::.:  *

STXBP1     465 KERISEQTYQLSRWTPIIKDIMEDTIEDKLDTKHYPYISTRSSASFSTT----------------AVSARYGHWHKNKAP-----
KEULE      503 KERQEEAAWQLSRFYPMIEELIEKLSKGELPKEDFPCMNDPSPSFHGSTSLSSAASSSQGQAAQSMRSRRTPTWAKPRGSDDGYS
Consensus  511 *** .* ::****: *:*::::*.  :.:* .:.:* :.  * : ..:*                  * *   * * :.

STXBP1     529 ----------GEYRSGPRLIIFILGGVSLNEMRCAYEVTQANGKWEVLIGSTHILTPQKLLDTLKKLNKTDEEISS----
KEULE      588 SDSVLRHASSDFRKMGQRIFVFIVGGATRSELKVCHKLS-TKLKREVILGSTSLDDPPQFITKLKLLTANDDLSLDDLQI
Consensus  596           .  : * *:::**:**.: .*:: .:::: :: * **::*** :  * ::: .** *. .*:   .
```

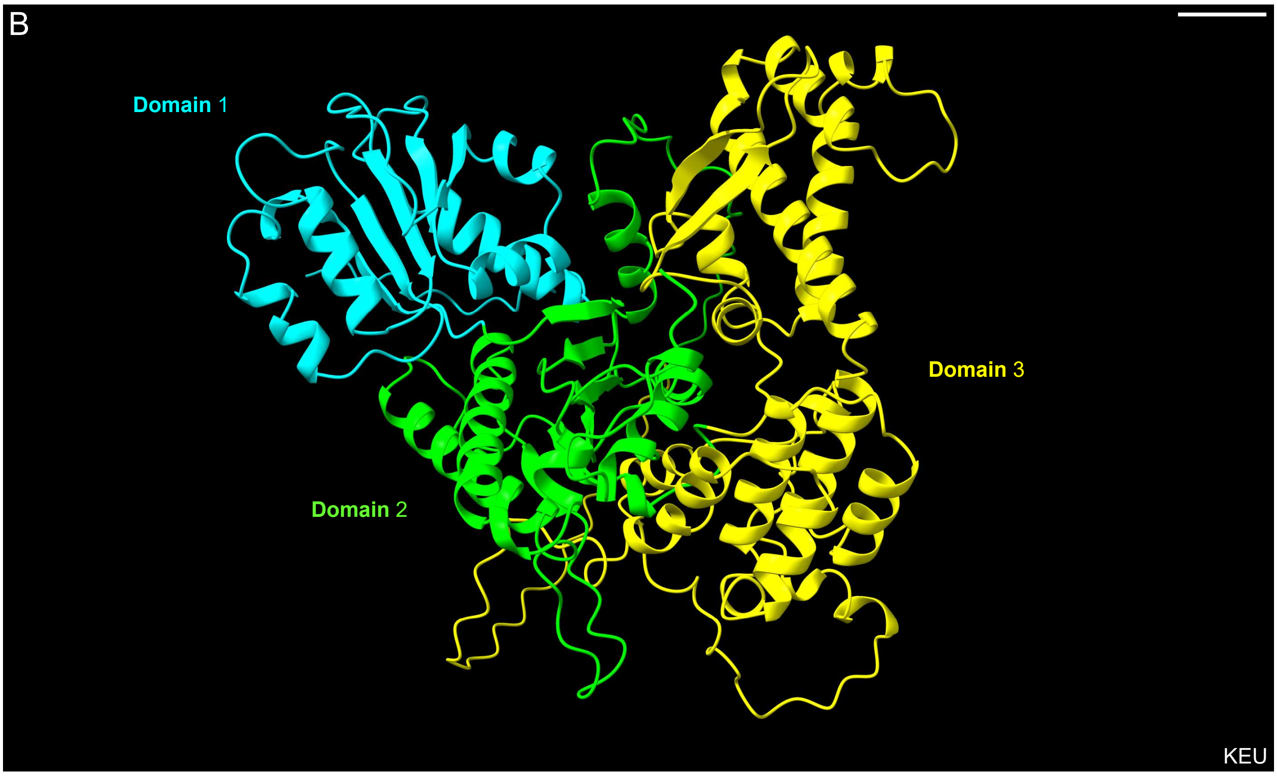


**Figure S2.** Sequence conservation between Arabidopsis KEU and the rat STXBP1 proteins. (A) Amino acid residues are indicated by numbers. The amino acids that physically interact with syntaxins are underlined. Residues of STXBP1 that form the secondary structures of domains 1, 2, and 3 are highlighted in blue, green, and yellow, respectively. The amino acids affected by the mutations of *sea4-1* (264) and *sea4-2* (57) are shaded in red. The amino acids with identical, similar, or weakly similar properties are denoted on the consensus line using asterisks, colons, and periods, respectively. (B) Cartoon representation of the 3D structure of KEU, with domains 1, 2, and 3 highlighted using the same color scheme. Scale bar: 10 Å.

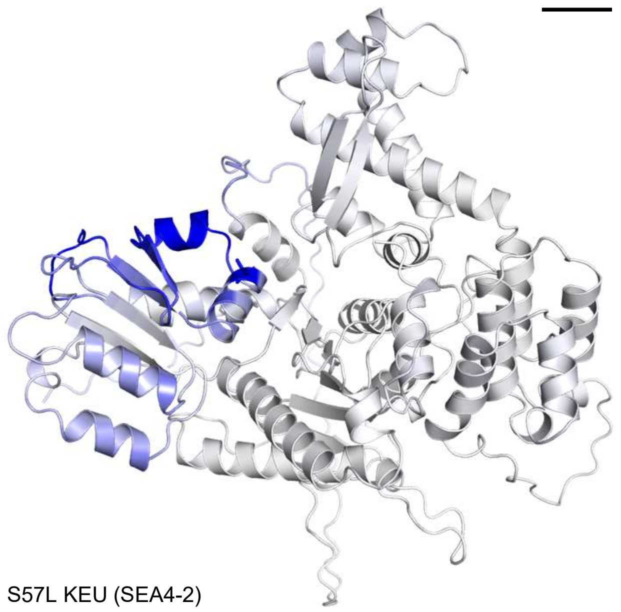


**Figure S3.** Predicted effect of the S57L mutation on the dynamics of the KEU protein. Cartoon representation of the 3D structure of the mutated KEU protein, colored according to the change in vibrational entropy ($\Delta\Delta S_{Vib}$ ENCoM) predicted by the DynaMut server. Blue corresponds to increased rigidity in the protein structure. Scale bar: 10 Å.

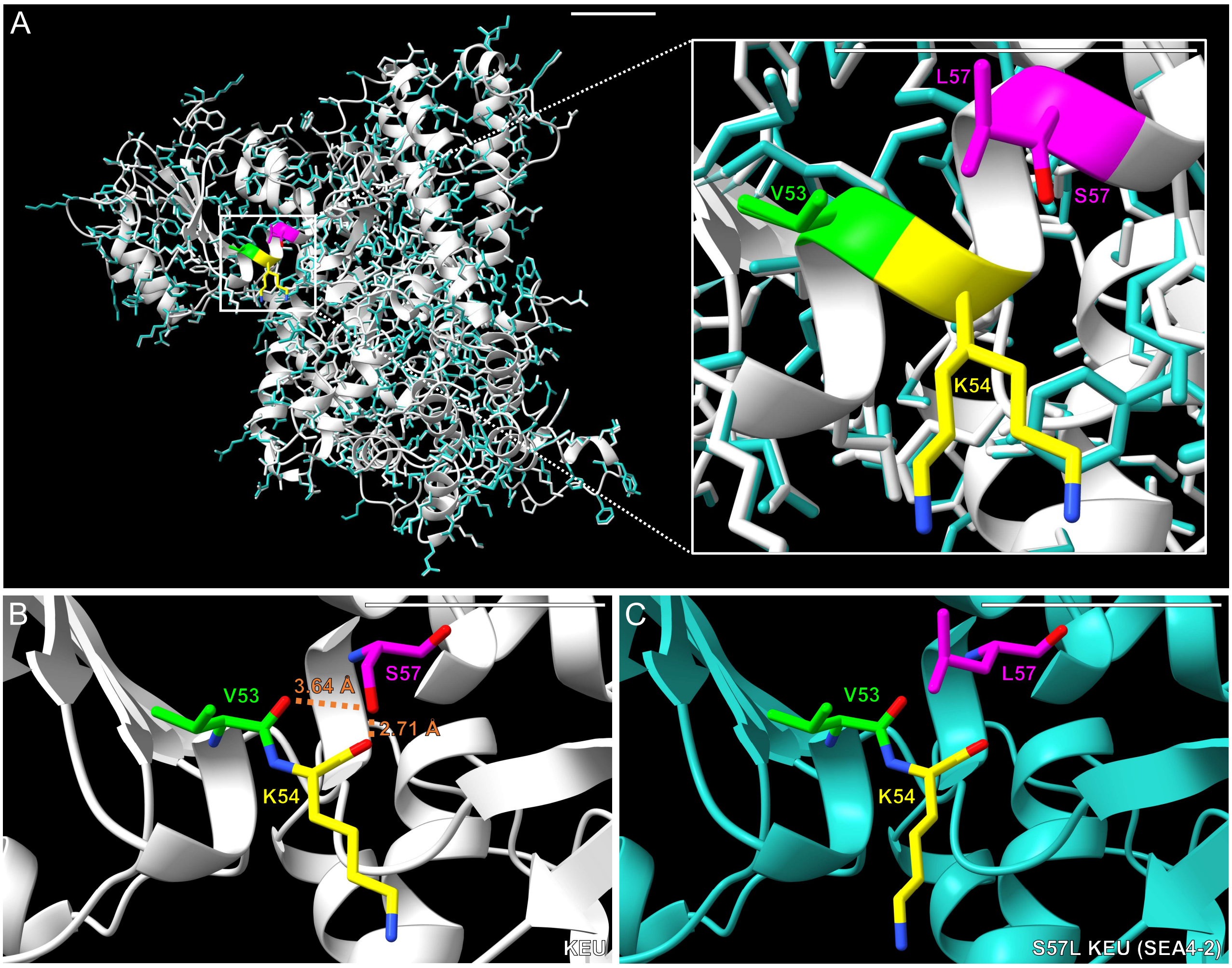


**Figure S4.** Comparison of the 3D structures of the wild-type and S57L mutant (SEA4-2) KEU proteins. (A) Cartoon representation showing the overlay of the 3D structures of KEU (colored in gray) and SEA4-2 (colored in blue), with a close-up view of the vicinity of the substituted S residue. All residues affected by the mutation are labeled and displayed in different colors. Hydrogen bonds between amino acids in (B) KEU, which are disrupted by the S57L mutation in (C) SEA4-2, are represented as orange dotted lines. The distances between the interacting atoms are indicated in angstroms (Å). The oxygen and nitrogen atoms of the affected amino acids are highlighted in red and blue, respectively. Scale bars: 10 Å.

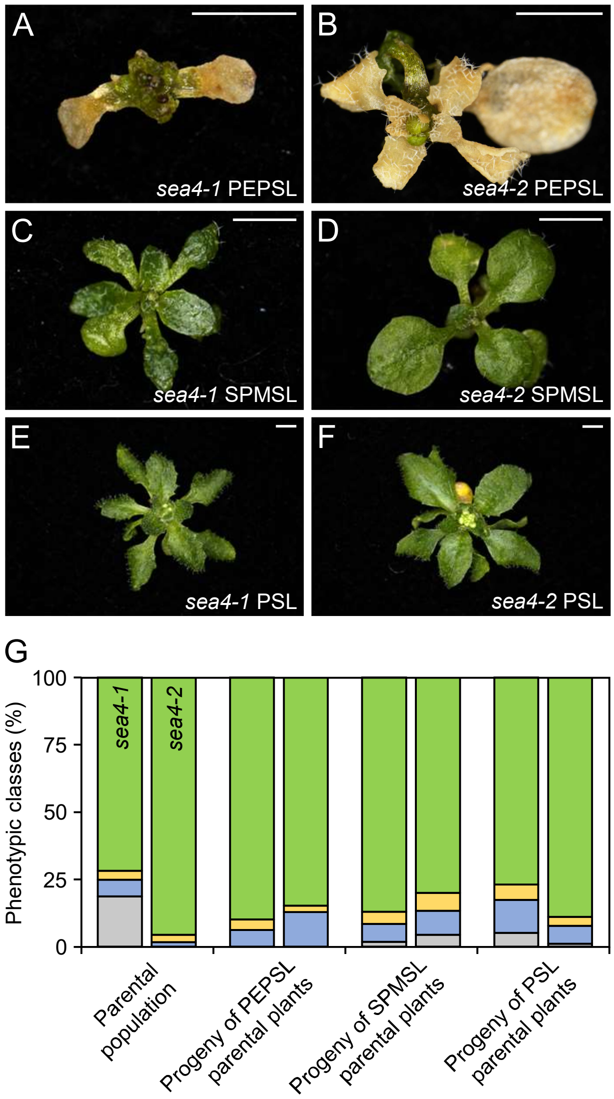


**Figure S5.** Phenotypic classes observed in *sea4-1* and *sea4-2* plants. A total of 180 (A, C, E) *sea4-1* and (B, D, F) *sea4-2* seeds were sown and the plants were grouped into four phenotypic classes based on their morphological phenotypes at 21 das: non-germinated seeds (NGS; not shown), (A, B) plants with poorly expanded and prematurely senescent leaves (PEPSL), (C, D) stunted plants with mildly serrated leaves (SPMSL), and (E, F) plants with serrated leaves (PSL). From each of the four viable classes, three plants were allowed to self to collect their seeds. 180 seeds per class were sown and the resulting plants classified as described above. (G) Stacked bar charts representing the percentages of plants in each phenotypic class defined above for two-year-old seeds (labeled as "Parental" on the X-axis) and the progeny of three selfed plants from each phenotypic class. Gray, blue, yellow, and green represent the NGS, PEPSL, SPMSL, and PSL classes, respectively. Photographs were taken 21 das. Scale bars: 2 mm.

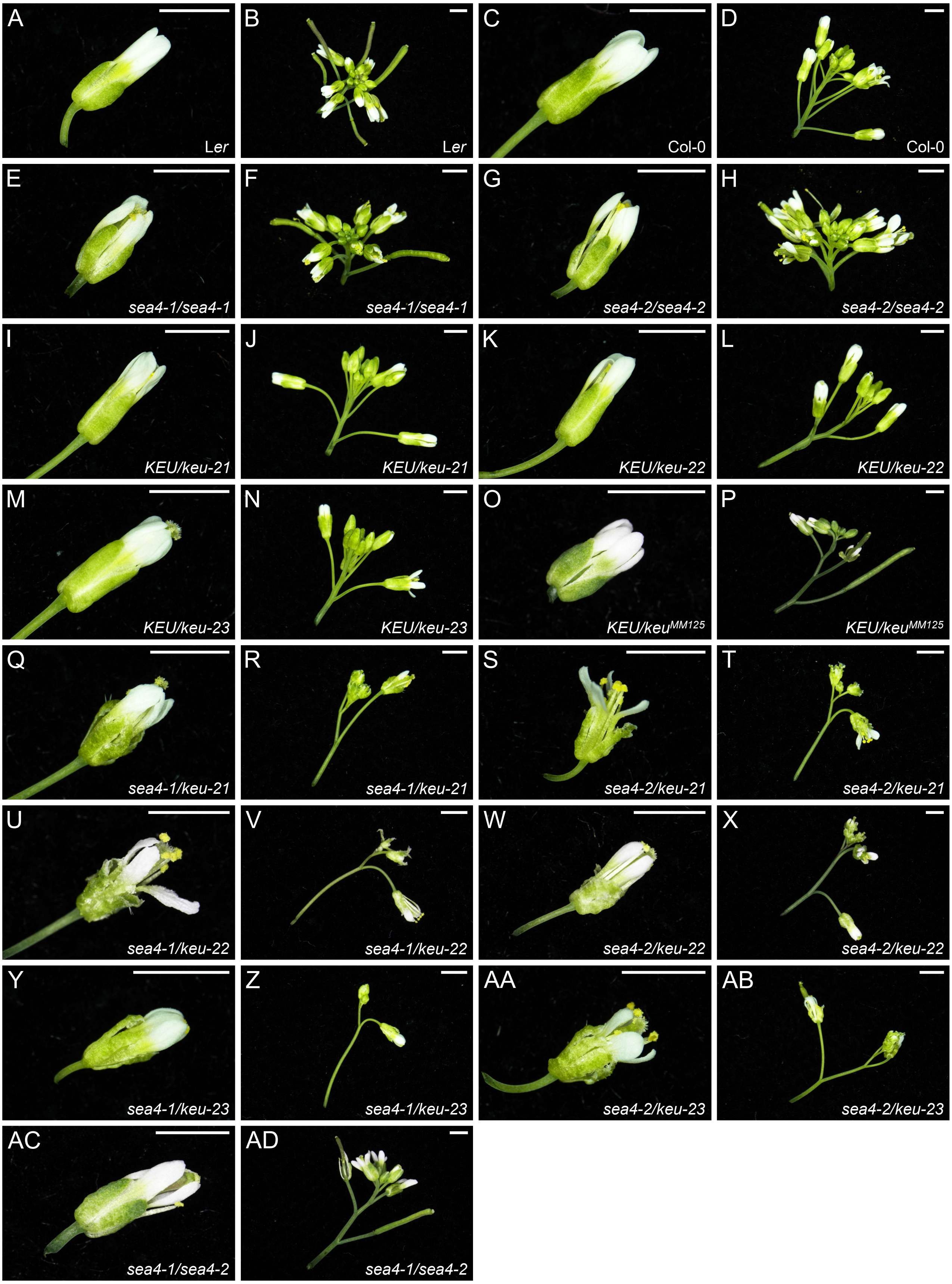


**Figure S6.** Flower and inflorescence morphology of plants carrying mutant alleles of *KEU*. (A, C, E, G, I, K, M, O, Q, S, U, W, Y, AA, AC) Flowers and (B, D, F, H, J, L, N, P, R, T, V, X, Z, AB, AD) inflorescences of wild-type (A, B) L*er* and (C, D) Col-0, the (E, F) *sea4-1/sea4-1* and (G, H) *sea4-2/sea4-2* homozygous mutants, and the (I, J) *KEU/keu-21*, (K, L) *KEU/keu-22*, (M, N) *KEU/keu-23*, (O, P) *KEU/keu*$^{MM125}$, (Q, R) *sea4-1/keu-21*, (S, T) *sea4-2/keu-21*, (U, V) *sea4-1/keu-22*, (W, X) *sea4-2/keu-22*, (Y, Z) *sea4-1/keu-23*, (AA, AB) *sea4-2/keu-23* and (AC, AD) *sea4-1/se4-2* heterozygotes. Photographs were taken 49 das. Scale bars: 2 mm.

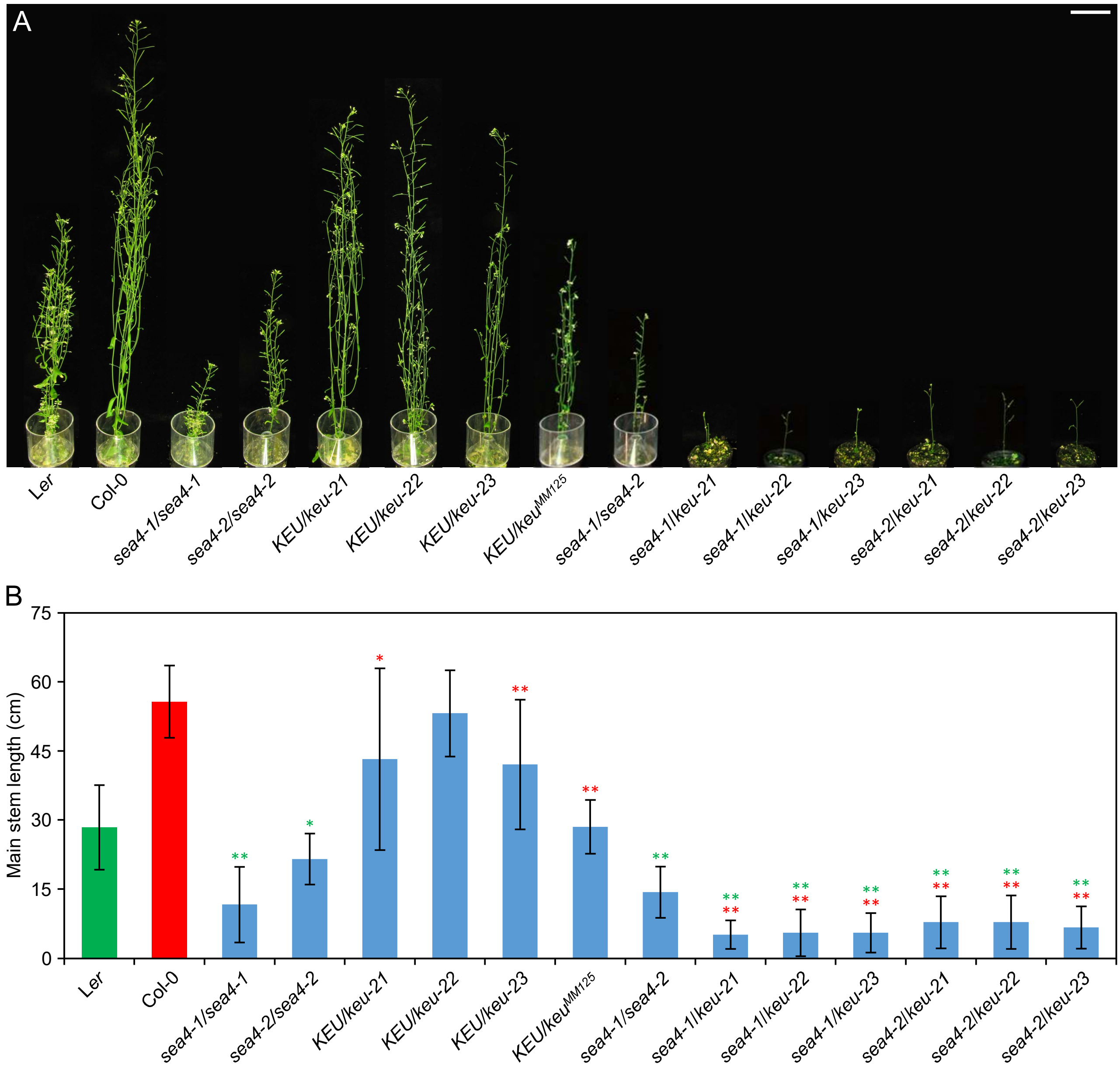


**Figure S7.** Morphological phenotypes of adult plants carrying mutant alleles of *KEU*. (A) From left to right, representative adult plants of L*er*, Col-0, *sea4-1/sea4-1*, *sea4-2/sea4-2*, *KEU/keu-21*, *KEU/keu-22*, *KEU/keu-23*, *KEU/keu*$^{MM125}$, *sea4-1/sea4-2*, *sea4-1/keu-21*, *sea4-1/keu-22*, *sea4-1/keu-23*, *sea4-2/keu-21*, *sea4-2/keu-22*, and *sea4-2/keu-23*. (B) Length of the main stem in adult plants. Error bars represent the standard deviation. Asterisks indicate a significant difference from the corresponding parental lines (indicated by the color code) in a Student's *t* test ($6 < n < 30$; **P* < 0.05, ***P* < 0.01). Photographs were taken 48 das. Scale bar: 5 cm.

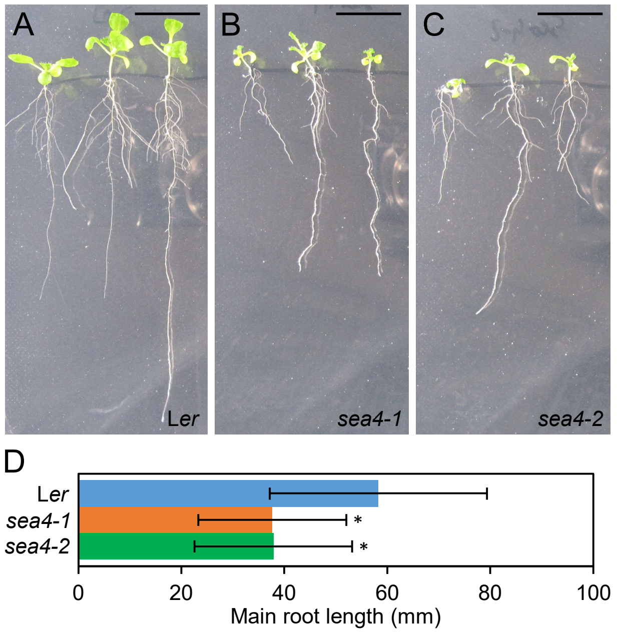


**Figure S8.** Main root length of *sea4-1* and *sea4-2* plants. Photographs of (A) L*er*, (B) *sea4-1*, and (C) *sea4-2* plants grown in vertical plates. (D) Main root length of Ler (n = 60), *sea4-1* (n = 30), and *sea4-2* (n = 30). Asterisks indicate a significant difference from L*er* in a Student's *t* test (P < 0.01). Photographs were taken 14 das. Scale bars: 5 mm.

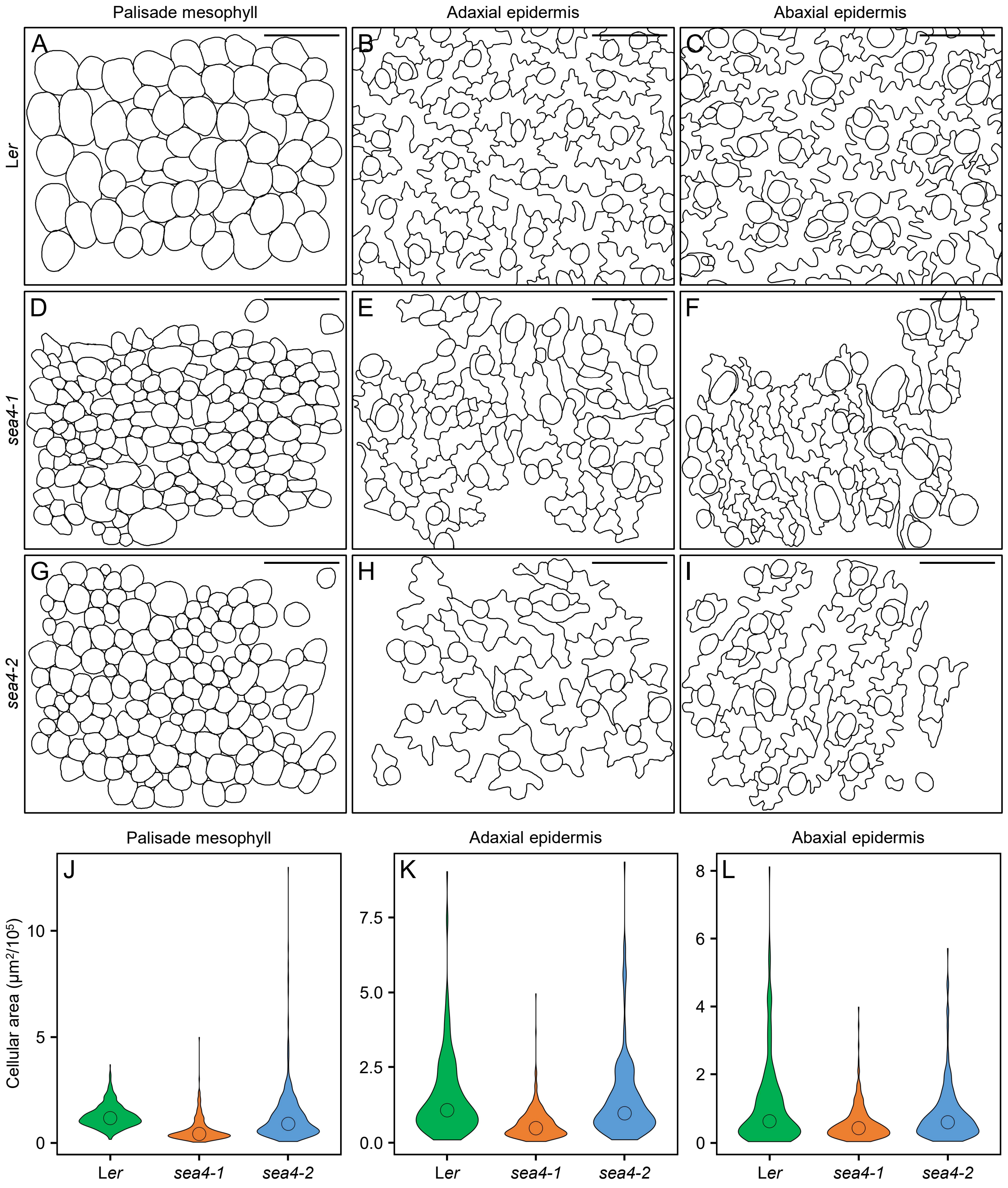


**Figure S9.** Structure of the cell layers in first-node leaves of the *sea4* mutants. (A-I) Diagrams illustrating cells from the (A, D, G) palisade mesophyll, (B, E, H) adaxial epidermis, and (C, F, I) abaxial epidermis of (A, B, C) L*er*, (D, E, F) *sea4-1*, and (G, H, I) *sea4-2* plants. Leaves were collected 21 das. Scale bars: 20 μm. (J-L) Violin plots representing the distribution of cell sizes in the (J) palisade mesophyll, (K) adaxial epidermis, and (L) abaxial epidermis. The median is represented with a circle.

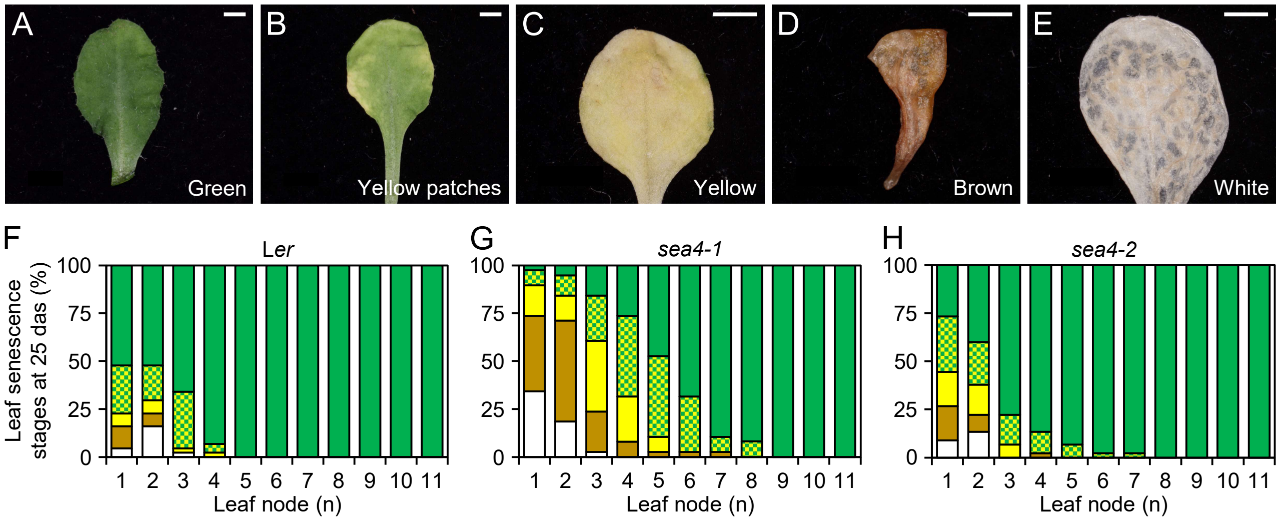


**Figure S10.** Progression of leaf senescence in *sea4-1* and *sea4-2* plants. All rosette leaves from 48 plants per genotype were grouped into the following categories at 25 das based on their degree of senescence: (A) green, (B) green with yellow patches, (C) yellow, (D) brown, or (E) white. The percentage of leaves from each node classified into these categories is represented for (F) L*er*, (G) *sea4-1*, and (H) *sea4-2*. The green, green/yellow pattern, yellow, brown, and white colors correspond to the categories shown in A, B, C, D, and E, respectively. Photographs were taken 25 das. Scale bars: 2 mm.

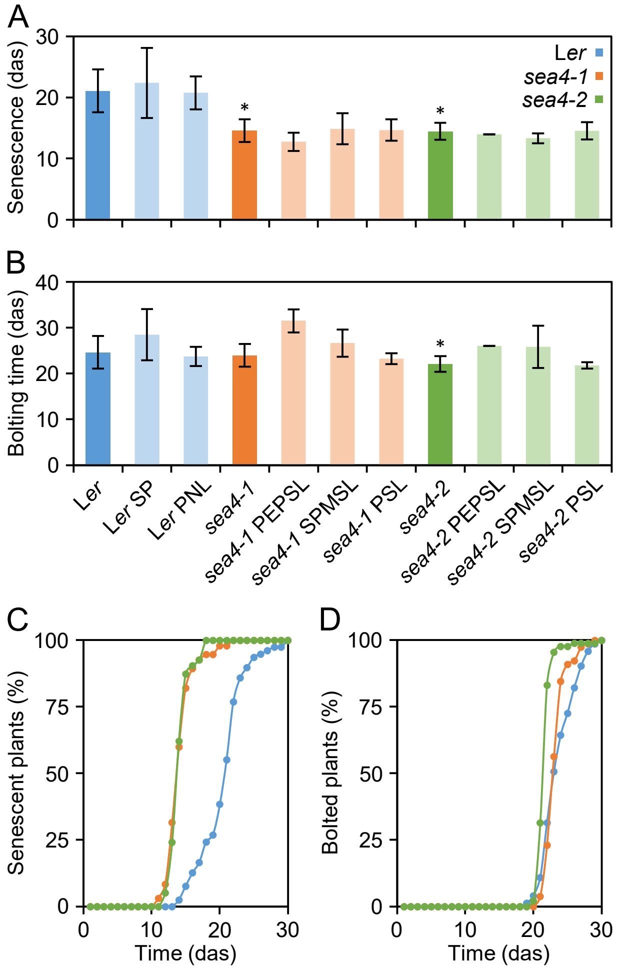


**Figure S11.** Extent of senescence and bolting time in the *sea4* mutants. Time at which (A) the first senescent leaf became visible in a plant and (B) bolting became visible. The mean values of all L*er*, *sea4-1*, and *sea4-2* plants studied are shown in dark blue, dark orange, and dark green, respectively. The mean values of L*er*, *sea4-1*, and *sea4-2* plants classified as in Figure S5 are shown in pale blue, pale orange, and pale green, respectively. Error bars represent standard deviation. Asterisks indicate a significant difference from L*er* in a Student's *t* test (78 < n < 90; **P* < 0.01). Percentage of plants (C) with senescent leaves and (D) exhibiting bolting over time after stratification. Colors for L*er*, *sea4-1*, and *sea4-2* are the same as in A.

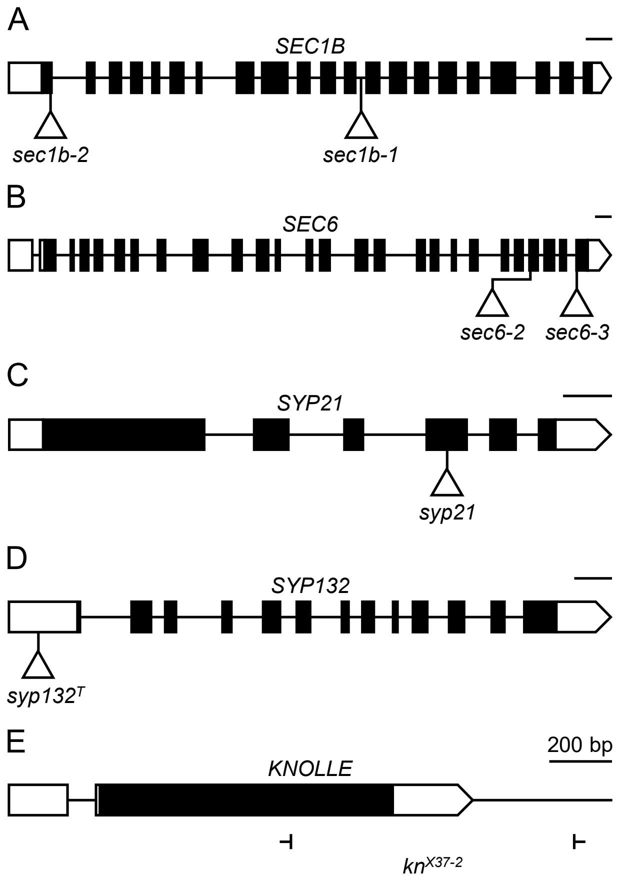


**Figure S12.** Structures of the genes that genetically interact with *KEU*, indicating the nature and positions of their mutations. Structures are shown for (A) SEC1B, (B) SEC6, (C) SYP21, (D) SYP132 and (E) KNOLLE genes. Boxes and lines between boxes indicate exons and introns, respectively. White boxes represent the 5′- and 3′-UTRs. Triangles indicate the T-DNA insertions in *kn*$^{X37\text{-}2}$, *sec1b-2*, *sec1b-1*, *syp21*, *syp132*$^{T}$, *sec6-2*, and *sec6-3*, and the ⊢ symbols delimitate the deletion in *kn*$^{X37\text{-}2}$. For simplicity, only one isoform of *SYP132* is shown.

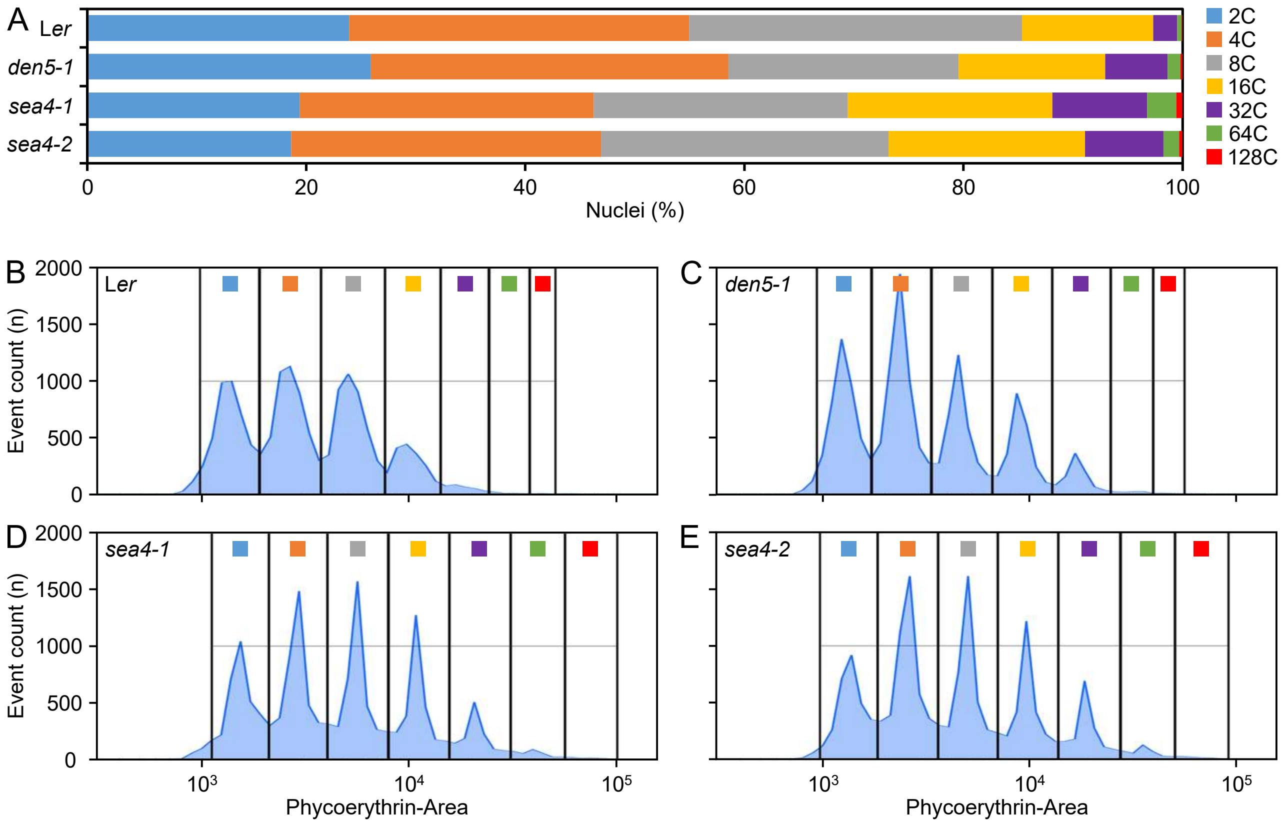


**Figure S13.** Nuclear ploidy levels in *sea4* leaves. (A) Distribution of nuclear DNA ploidy in L*er*, *den5-1*, *sea4-1*, and *sea4-2* plants. Populations with different ploidy levels are indicated by different colors. (B-E) Histograms representing the frequency distribution of propidium iodide fluorescence (detected by the phycoerythrin [PE] detector of the flow cytometer) in the first pair of leaves from (B) L*er*, (C) *den5-1*, (D) *sea4-1*, and (E) *sea4-2* plants at 21 das. Gates delineate the populations of nuclei with different ploidy levels. Squares with the same color scheme as in A represent the populations with different ploidy levels.

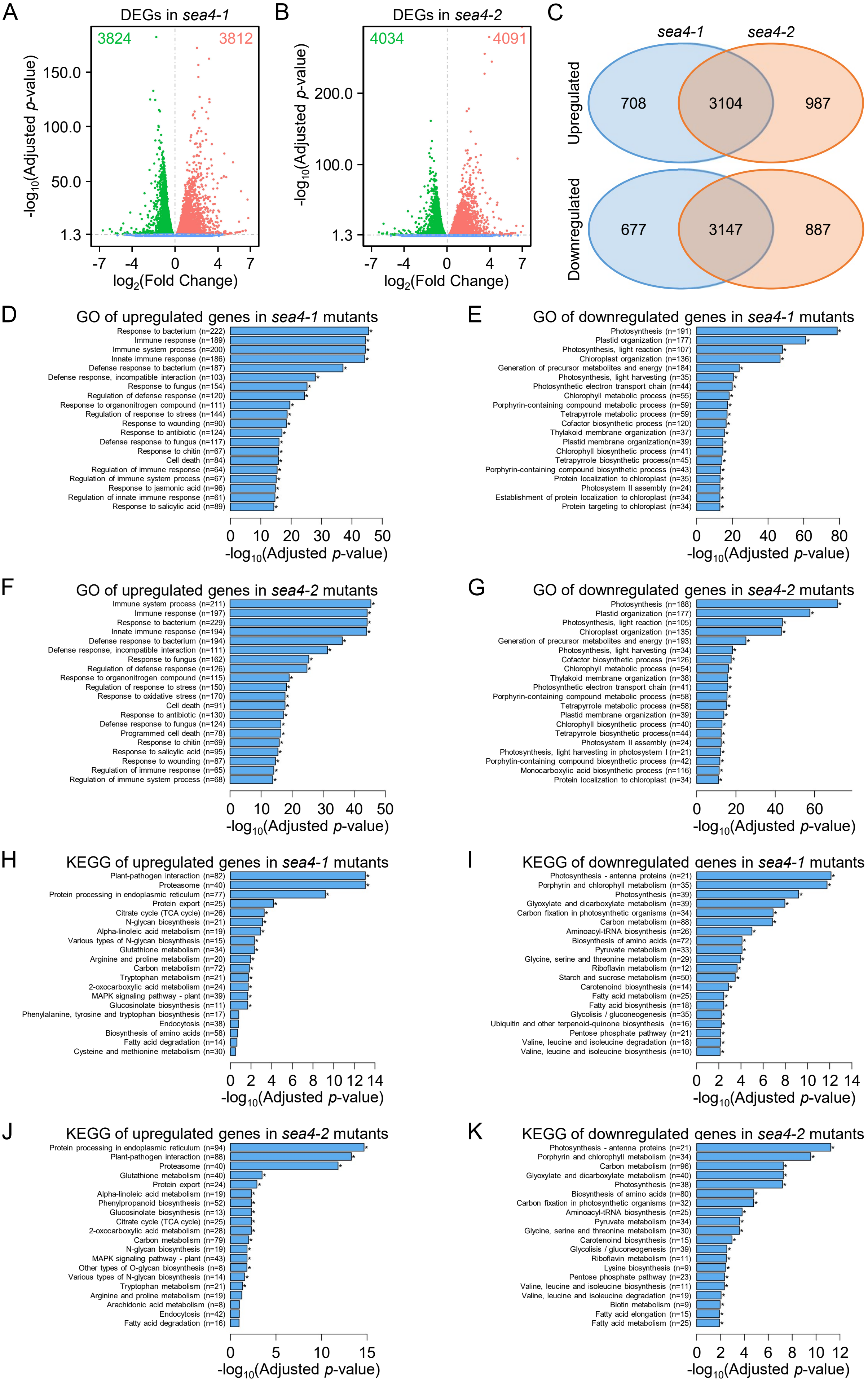
A
DEGs in sea4-1
3824
3812
-log10(Adjusted p-value)
150.0
100.0
50.0
1.3
-7 -4 0 4 7
log2(Fold Change)
B
DEGs in sea4-2
4034
4091
-log10(Adjusted p-value)
200.0
100.0
1.3
-7 -4 0 4 7
log2(Fold Change)
C
sea4-1
sea4-2
Upregulated
708
3104
987
Downregulated
677
3147
887
D
GO of upregulated genes in sea4-1 mutants
Response to bacterium (n=222)
Immune response (n=189)
Immune system process (n=200)
Innate immune response (n=186)
Defense response to bacterium (n=187)
Defense response, incompatible interaction (n=103)
Response to fungus (n=154)
Regulation of defense response (n=120)
Response to organonitrogen compound (n=111)
Regulation of response to stress (n=144)
Response to wounding (n=90)
Response to antibiotic (n=124)
Defense response to fungus (n=117)
Response to chitin (n=67)
Cell death (n=84)
Regulation of immune response (n=64)
Regulation of immune system process (n=67)
Response to jasmonic acid (n=96)
Regulation of innate immune response (n=61)
Response to salicylic acid (n=89)
0 10 20 30 40 50
-log10(Adjusted p-value)
E
GO of downregulated genes in sea4-1 mutants
Photosynthesis (n=191)
Plastid organization (n=177)
Photosynthesis, light reaction (n=107)
Chloroplast organization (n=136)
Generation of precursor metabolites and energy (n=184)
Photosynthesis, light harvesting (n=35)
Photosynthetic electron transport chain (n=44)
Chlorophyll metabolic process (n=55)
Porphyrin-containing compound metabolic process (n=59)
Tetrapyrrole metabolic process (n=59)
Cofactor biosynthetic process (n=120)
Thylakoid membrane organization (n=37)
Plastid membrane organization(n=39)
Chlorophyll biosynthetic process (n=41)
Tetrapyrrole biosynthetic process(n=45)
Porphyrin-containing compound biosynthetic process (n=43)
Protein localization to chloroplast (n=35)
Photosystem II assembly (n=24)
Establishment of protein localization to chloroplast (n=34)
Protein targeting to chloroplast (n=34)
0 20 40 60 80
-log10(Adjusted p-value)
F
GO of upregulated genes in sea4-2 mutants
Immune system process (n=211)
Immune response (n=197)
Response to bacterium (n=229)
Innate immune response (n=194)
Defense response to bacterium (n=194)
Defense response, incompatible interaction (n=111)
Response to fungus (n=162)
Regulation of defense response (n=126)
Response to organonitrogen compound (n=115)
Regulation of response to stress (n=150)
Response to oxidative stress (n=170)
Cell death (n=91)
Response to antibiotic (n=130)
Defense response to fungus (n=124)
Programmed cell death (n=78)
Response to chitin (n=69)
Response to salicylic acid (n=95)
Response to wounding (n=87)
Regulation of immune response (n=65)
Regulation of immune system process (n=68)
0 10 20 30 40 50
-log10(Adjusted p-value)
G
GO of downregulated genes in sea4-2 mutants
Photosynthesis (n=188)
Plastid organization (n=177)
Photosynthesis, light reaction (n=105)
Chloroplast organization (n=135)
Generation of precursor metabolites and energy (n=193)
Photosynthesis, light harvesting (n=34)
Cofactor biosynthetic process (n=126)
Chlorophyll metabolic process (n=54)
Thylakoid membrane organization (n=38)
Photosynthetic electron transport chain (n=41)
Porphyrin-containing compound metabolic process (n=58)
Tetrapyrrole metabolic process (n=58)
Plastid membrane organization (n=39)
Chlorophyll biosynthetic process (n=40)
Tetrapyrrole biosynthetic process(n=44)
Photosystem II assembly (n=24)
Photosynthesis, light harvesting in photosystem I (n=21)
Porphyrin-containing compound biosynthetic process (n=42)
Monocarboxylic acid biosynthetic process (n=116)
Protein localization to chloroplast (n=34)
0 20 40 60
-log10(Adjusted p-value)
H
KEGG of upregulated genes in sea4-1 mutants
Plant-pathogen interaction (n=82)
Proteasome (n=40)
Protein processing in endoplasmic reticulum (n=77)
Protein export (n=25)
Citrate cycle (TCA cycle) (n=26)
N-glycan biosynthesis (n=21)
Alpha-linoleic acid metabolism (n=19)
Various types of N-glycan biosynthesis (n=15)
Glutathione metabolism (n=34)
Arginine and proline metabolism (n=20)
Carbon metabolism (n=72)
Tryptophan metabolism (n=21)
2-oxocarboxylic acid metabolism (n=24)
MAPK signaling pathway - plant (n=39)
Glucosinolate biosynthesis (n=11)
Phenylalanine, tyrosine and tryptophan biosynthesis (n=17)
Endocytosis (n=38)
Biosynthesis of amino acids (n=58)
Fatty acid degradation (n=14)
Cysteine and methionine metabolism (n=30)
0 2 4 6 8 10 12 14
-log10(Adjusted p-value)
I
KEGG of downregulated genes in sea4-1 mutants
Photosynthesis - antenna proteins (n=21)
Porphyrin and chlorophyll metabolism (n=35)
Photosynthesis (n=39)
Glyoxylate and dicarboxylate metabolism (n=39)
Carbon fixation in photosynthetic organisms (n=34)
Carbon metabolism (n=88)
Aminoacyl-tRNA biosynthesis (n=26)
Biosynthesis of amino acids (n=72)
Pyruvate metabolism (n=33)
Glycine, serine and threonine metabolism (n=29)
Riboflavin metabolism (n=12)
Starch and sucrose metabolism (n=50)
Carotenoid biosynthesis (n=14)
Fatty acid metabolism (n=25)
Fatty acid biosynthesis (n=18)
Glycolysis / gluconeogenesis (n=35)
Ubiquitin and other terpenoid-quinone biosynthesis (n=16)
Pentose phosphate pathway (n=21)
Valine, leucine and isoleucine degradation (n=18)
Valine, leucine and isoleucine biosynthesis (n=10)
0 2 4 6 8 10 12 14
-log10(Adjusted p-value)
J
KEGG of upregulated genes in sea4-2 mutants
Protein processing in endoplasmic reticulum (n=94)
Plant-pathogen interaction (n=88)
Proteasome (n=40)
Glutathione metabolism (n=40)
Protein export (n=24)
Alpha-linoleic acid metabolism (n=19)
Phenylpropanoid biosynthesis (n=52)
Glucosinolate biosynthesis (n=13)
Citrate cycle (TCA cycle) (n=25)
2-oxocarboxylic acid metabolism (n=28)
Carbon metabolism (n=79)
N-glycan biosynthesis (n=19)
MAPK signaling pathway - plant (n=43)
Other types of O-glycan biosynthesis (n=8)
Various types of N-glycan biosynthesis (n=14)
Tryptophan metabolism (n=21)
Arginine and proline metabolism (n=19)
Arachidonic acid metabolism (n=8)
Endocytosis (n=42)
Fatty acid degradation (n=16)
0 5 10 15
-log10(Adjusted p-value)
K
KEGG of downregulated genes in sea4-2 mutants
Photosynthesis - antenna proteins (n=21)
Porphyrin and chlorophyll metabolism (n=34)
Carbon metabolism (n=96)
Glyoxylate and dicarboxylate metabolism (n=40)
Photosynthesis (n=38)
Biosynthesis of amino acids (n=80)
Carbon fixation in photosynthetic organisms (n=32)
Aminoacyl-tRNA biosynthesis (n=25)
Pyruvate metabolism (n=34)
Glycine, serine and threonine metabolism (n=30)
Carotenoid biosynthesis (n=15)
Glycolysis / gluconeogenesis (n=39)
Riboflavin metabolism (n=11)
Lysine biosynthesis (n=9)
Pentose phosphate pathway (n=23)
Valine, leucine and isoleucine biosynthesis (n=11)
Valine, leucine and isoleucine degradation (n=19)
Biotin metabolism (n=9)
Fatty acid elongation (n=15)
Fatty acid metabolism (n=25)
0 2 4 6 8 10 12
-log10(Adjusted p-value)

**Figure S14.** Differentially expressed genes in the *sea4* mutants identified by RNA-seq. Volcano plots for (A) *sea4-1* and (B) *sea4-2*, illustrating the overall distribution of differentially expressed genes (DEGs). Upregulated, downregulated, and non-differentially regulated genes are represented by red, green, and blue dots, respectively. Total number of upregulated and downregulated genes is indicated using the same color scheme. (C) Venn diagrams displaying the overlap between upregulated and downregulated genes in *sea4-1* and *sea4-2*. (D-G) GO and (H-K) KEGG enrichment analyses of upregulated (D, F, H, J) and downregulated (E, G, I, K) genes in (D, E, H, I) *sea4-1* and (F, G, J, K) *sea4-2*. The assays were performed with three biological replicates. The threshold for significance was set at padj < 0.05. n represents the number of DEGs associated with a specific GO or KEGG term.

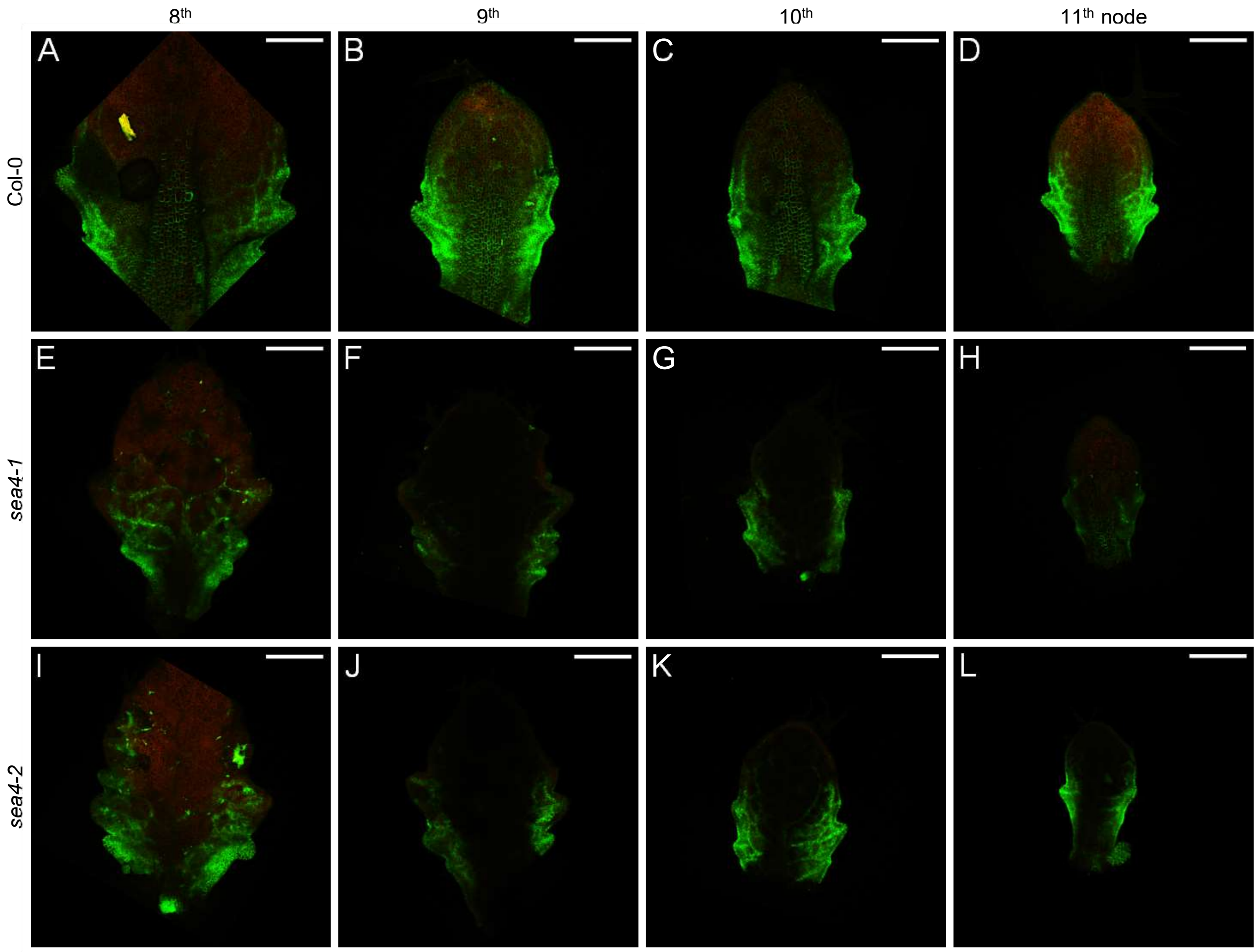


**Figure S15.** Expression pattern of the *PIN1*$_{pro}$*:PIN1:GFP* reporter in leaf primordia of the *sea4* mutants. The visualization of *PIN1*$_{pro}$*:PIN1:GFP* (green) expression in leaf primordia from successive nodes (8$^{th}$-11$^{th}$) is shown for (A-D) Col-0, (E-H) *sea4-1*, and (I-L) *sea4-2* plants. Chlorophyll autofluorescence is shown in red. The primordia were collected 20 das. In all figures, the primordia shown for each genotype were collected from the same plant. Scale bars: 0.2 mm.

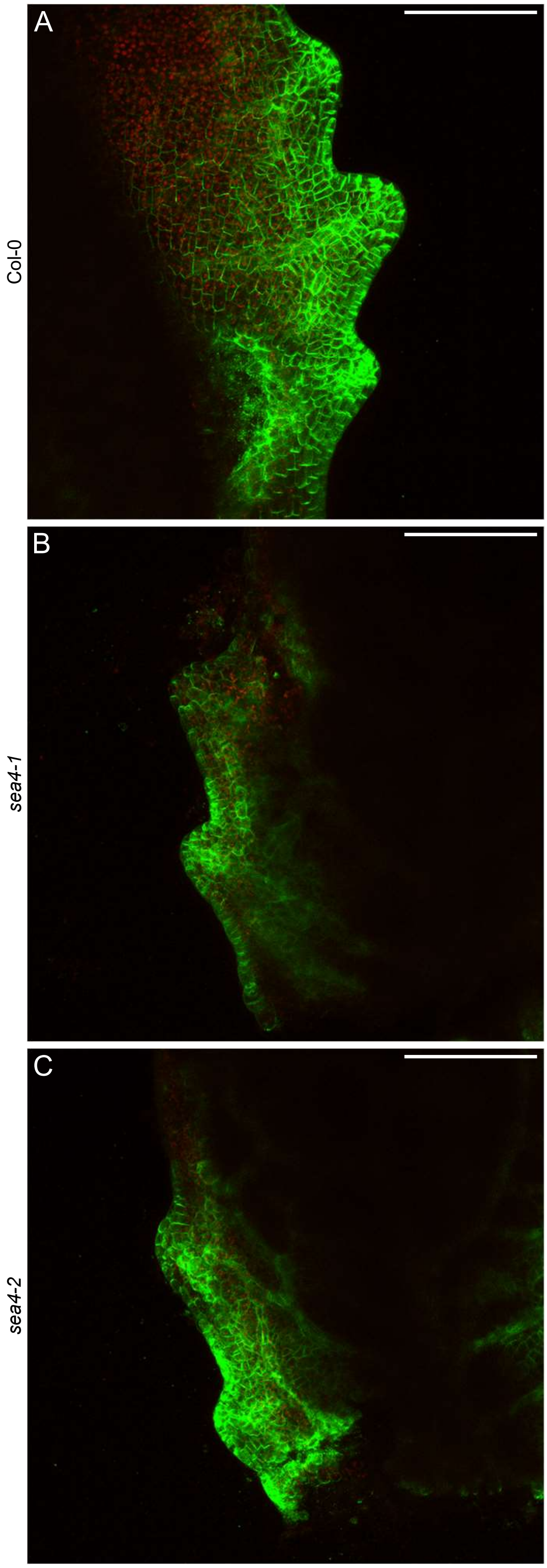


**Figure S16.** Localization of the *PIN1$_{pro}$:PIN1:GFP* reporter in the cell membrane in leaf primordia of the *sea4* mutants. The visualization of *PIN1$_{pro}$:PIN1:GFP* (green) expression at the margin of leaf primordia from the 10$^{th}$ node is shown for (A) Col-0, (B) *sea4-1*, and (C) *sea4-2* plants. Chlorophyll autofluorescence is shown in red. The primordia were collected 20 das. Scale bars: 0.2 mm.

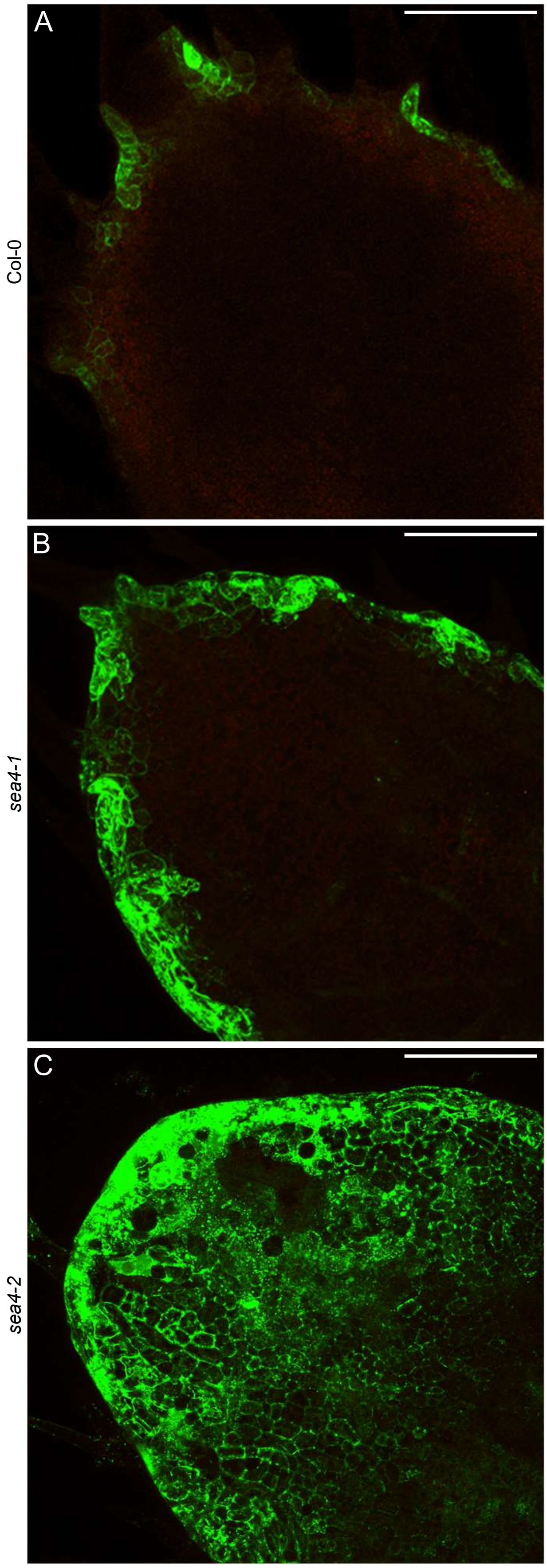


**Figure S17.** Localization of the *DR5rev*$_{pro}$*:GFP* reporter in the cell membrane in leaf primordia of the *sea4* mutants. The visualization of *DR5rev*$_{pro}$*:GFP* (green) expression at the apex of leaf primordia from the 9$^{th}$ node is shown for (A) Col-0, (B) *sea4-1*, and (C) *sea4-2* plants. Chlorophyll autofluorescence is shown in red. The primordia were collected 20 das. Scale bars: 0.2 mm.

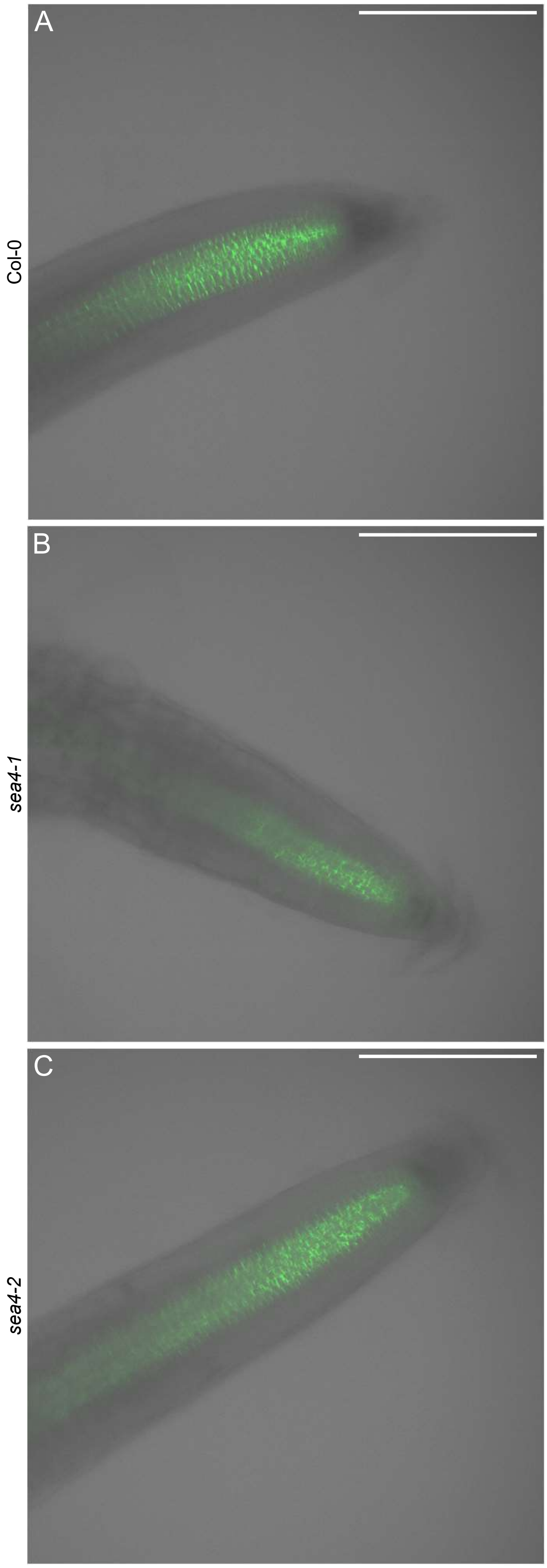


**Figure S18.** Expression pattern of the *PIN1$_{pro}$:PIN1:GFP* reporter in the root apex of the *sea4* mutants. The visualization of *PIN1$_{pro}$:PIN1:GFP* (green) expression in the root apex is shown for (A) Col-0, (B) *sea4-1*, and (C) *sea4-2* plants. Transillumination is shown in gray. The roots were collected 7 das. Scale bars: 0.2 mm.

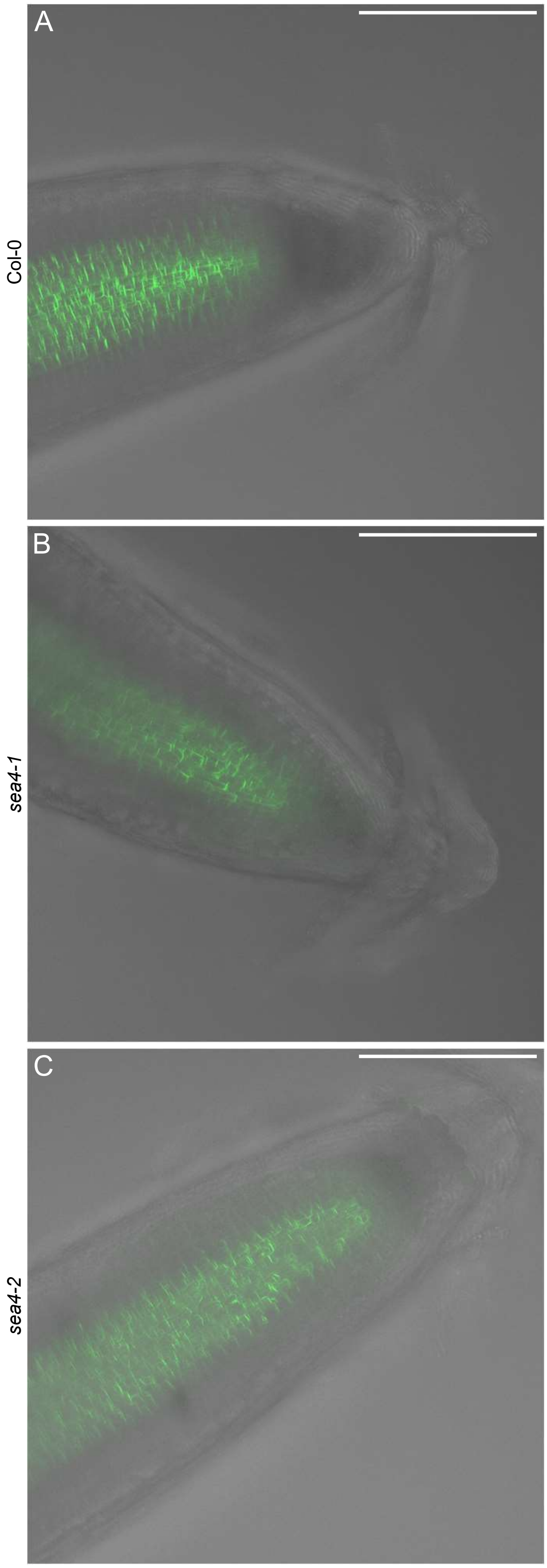


**Figure S19.** Localization of the *PIN1*$_{pro}$*:PIN1:GFP* reporter in the cell membrane in roots of the *sea4* mutants. The visualization of *PIN1*$_{pro}$*:PIN1:GFP* (green) expression in the root apex is shown for (A) Col-0, (B) *sea4-1*, and (C) *sea4-2* plants. Transillumination is shown in gray. The roots were collected 7 das. Scale bars: 0.2 mm.

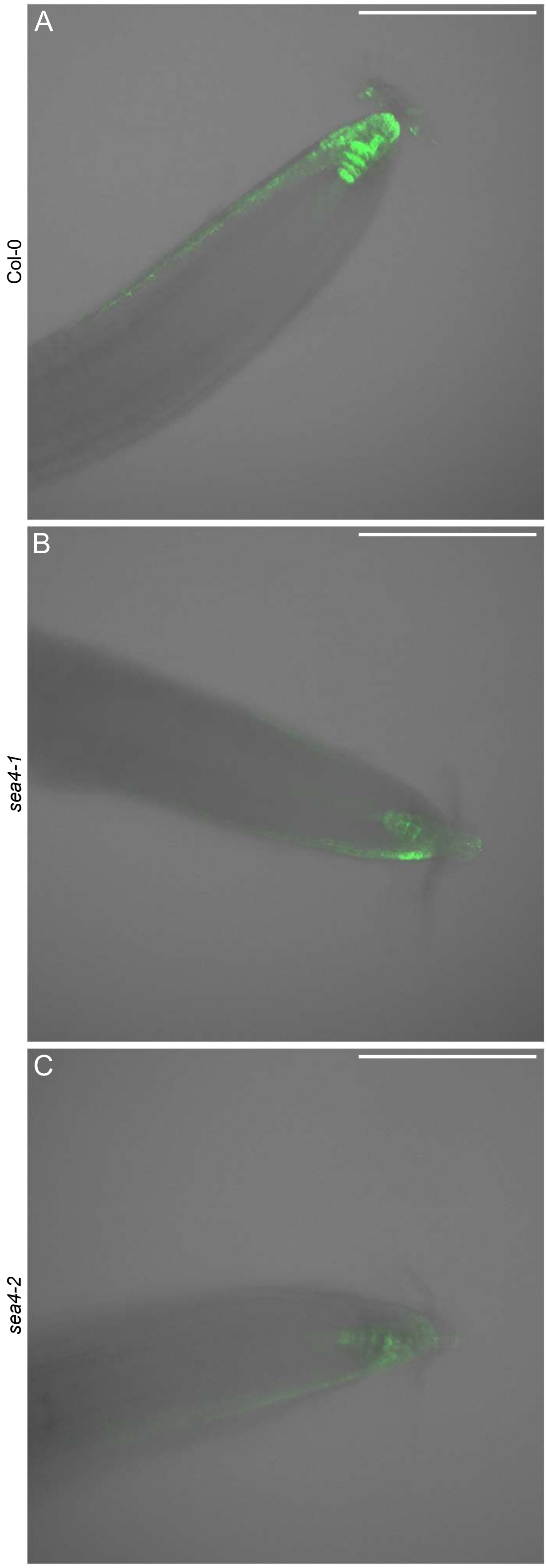


**Figure S20.** Expression pattern of the *DR5rev*$_{pro}$*:GFP* reporter in the root apex of the *sea4* mutants. The visualization of *DR5rev*$_{pro}$*:GFP* (green) expression in the root apex is shown for (A) Col-0, (B) *sea4-1*, and (C) *sea4-2* plants. Transillumination is shown in gray. The roots were collected 7 das. Scale bars: 0.2 mm.

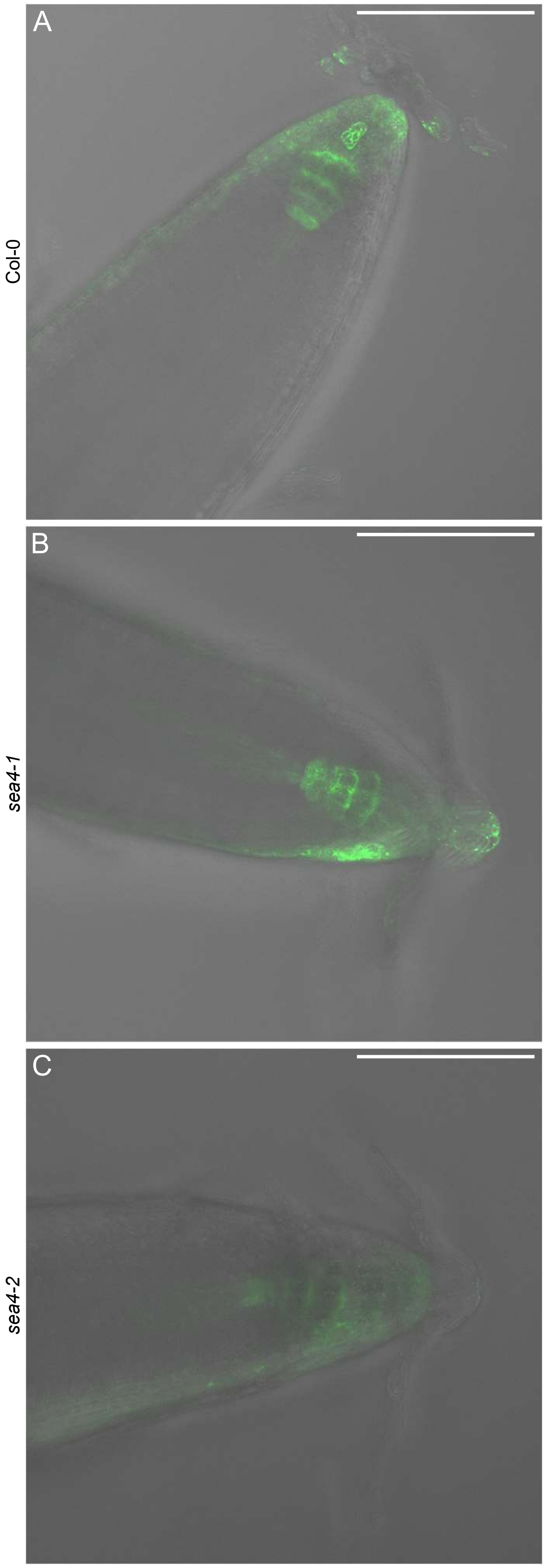


**Figure S21.** Localization of the *DR5rev*$_{pro}$*:GFP* reporter in the cell membrane in roots of the *sea4* mutants. The visualization of *DR5rev*$_{pro}$*:GFP* (green) expression in the root apex is shown for (A) Col-0, (B) *sea4-1*, and (C) *sea4-2* plants. Transillumination is shown in gray. The roots were collected 7 das. Scale bars: 0.2 mm.

**Table S1.** Primer sets used in this work

| Purpose | Names | Oligonucleotide sequence (5′ → 3′) | |
|---|---|---|---|
| | | Forward primer (F or D) | Reverse primer (R or C) |
| Low-resolution mapping | nga59-F/R[a] | ATCTGTGTTCACTCGCCGCC | GCCTAACAAATTTAAAGTTAAGACT |
| | nga63-F/R | AACCAAGGCACAGAAGCG | ACCCAAGTGATCGCCACC |
| | cer465532-F/R | CATTACACTAGAGACTAGAC | TAATATACATGTGAGCATCCTGAC |
| | F16J7-TRB-F/R | GTGTCTTGATACGCGTCGATC | TGATGTTGAGATCTGTGTGCAG |
| | cer448767-F/R | GCTAGCAGTCGTAACTCATAA | TGGGATAAACTCGTTGTCGTT |
| | cer474022-F/R | GGAGATAAGCGATGAACGAGA | CGGATTAGCGCAGAGGGTTT |
| | cer451935-F/R | CCAACAAATTCGTAAACGGCGAA | GGGCTAGGTTAGTCTCCCTT |
| | JV18/19-F/R[a] | TGTCGTATATCAATCGAAAAAGAG | AATTCAGTATCGAGATACCCCT |
| | AthZFPG-F/R[b] | TTGCGTTTCCACATTTGTTT | TGGGTCAATTCACATGTAGAGA |
| Sequencing | At1g12360-5F/R | GCTCACAATGAAAGCATGTACTG | CTCATGTATAACAGGGGCAATC |
| Genotyping of *keu-21, keu-22* and *keu-23* | At1g12360-2F/2R | GGTACTTGGGGTTTTAGCGC | TGAATGCCTTGGTATTGTTACTTG |
| *keu*$^{MM125}$ | KEU in 14/KEU in 17[c] | CATGAGATATTGAGCTGATGAGC | AGTAGTGTCCTGTGCAAATAGG |
| *kn*$^{X37-2}$ | X37-2DIII/2CIII[c] | GGGATGGATATGGTGGTGC | AAGGCTCTCTGGGACTCCGG |
| SALK_072337C and SALK_100970 | At1g71820-2F/2R | TCGTGGTGAATACTGGATGCA | ATCCTAGTGCAAGTGGCCAC |
| SAIL_580_C04 | At5g16830-F/R | GGAGATATGAAGGAACGAGGA | CATGCATTTTGCTCTCGAGAG |
| SAIL_403_B09 | At5g08080-F/R | TGCCTGGATGTAACACGTGTA | ACTCTCACCTTCAGAAGATCG |
| GK-601G09 | At4g12120-F/R | AGCTGCCCGAGATCTCTTAG | GATGTATGTCGTGCCTCCTC |
| GK-283F10 | At4g12120-2F/2R | AGCTTCCATTTCTTTCTGGCC | CCCAGGAACGGCAAAAATCTA |
| T-DNA insertion verification | LBb1.3[d] | ATTTTGCCGATTTCGGAAC | |
| | LB (SAIL)[d] | GCCTTTTCAGAAATGGATAAATAGCCTTGCTTCC | |
| | LB-PAC161[d] | ATATTGACCATCATACTCATTGC | |

The *sea4-1* and *sea4-2* alleles were genotyped using the At1g12360-5F/R primer pair. [a-d]Sequences taken from [a]Ponce et al. (2006), [b]Ponce et al. (1999), [c]Karnahl et al. (2018) and [d]http://signal.salk.edu/tdnaprimers.2.html.

**Table S2.** Configuration parameters of the Leica Stellaris 8 STED confocal microscope

| Transgenic line | Objective | WLL intensity (%) | PMT Trans gain | LUT values for GFP | | LUT values for chlorophyll autofluorescence | | LUT values for transillumination | |
|---|---|---|---|---|---|---|---|---|---|
| | | | | Max | Min | Max | Min | Max | Min |
| *$PIN1_{pro}$:PIN1:GFP* | 20x | 50 | 10.76 | 255 | 0 | 255 | 0 | 255 | 255 |
| | 40x | 50 | 13.06 | 255 | 0 | 100 | 0 | 255 | 255 |
| *$DR5rev_{pro}$:GFP* | 20x | 8 | 14.76 | 200 | 0 | 60 | 0 | 255 | 255 |
| | 40x | 8 | 16.76 | 255 | 0 | 100 | 0 | 255 | 255 |

WLL: white light laser. PMT Trans: photon multiplier tube transillumination. LUT: look-up tables.

**Table S3.** Data quality control summary of the RNA-seq assay

| Sample | Number of clean reads | Q30 quality score* (%) | Mapped reads (%) |
|---|---|---|---|
| L*er* replicate #1 | 28585837 | 94.11 | 96.05 |
| L*er* replicate #2 | 31042172 | 93.68 | 95.43 |
| L*er* replicate #3 | 27461285 | 93.92 | 95.80 |
| *sea4-1* replicate #1 | 29491840 | 94.01 | 95.08 |
| *sea4-1* replicate #2 | 35082779 | 93.48 | 95.03 |
| *sea4-1* replicate #3 | 27397456 | 93.85 | 95.07 |
| *sea4-2* replicate #1 | 27252781 | 93.81 | 95.16 |
| *sea4-2* replicate #2 | 30189575 | 93.97 | 95.14 |
| *sea4-2* replicate #3 | 29108548 | 93.99 | 95.16 |

*Percentage of bases whose correct base recognition rates are greater than 99.9% in total bases.

**Table S4.** Candidate mutations identified in *sea4-1* by Easymap

| Position[a] | Allele Frequency | DTP[b] | Nucleotide (ref → alt)[c] | Gene | Gene element | Amino acid (ref → alt)[c] |
|---|---|---|---|---|---|---|
| 1787746 | 0.89 | – 1212254 | G → A | At1g05890 | CDS | G → E |
| 2165044 | 0.90 | – 834956 | G → A | At1g07030 | CDS | – |
| 2791816 | 0.95 | – 208184 | G → A | At1g08730 | CDS | – |
| 3010996 | 0.93 | 10996 | G → A | At1g09320 | Intron | – |
| 3253461 | 0.96 | 253461 | G → A | At1g09970 | CDS | – |
| 3363247 | 0.95 | 363247 | G → A | – | – | – |
| 3500496 | 0.96 | 500496 | G → A | At1g10540 | CDS | L → F |
| 3578311 | 1.00 | 578311 | G → A | At1g10700 | CDS | G → S |
| 3660993 | 0.92 | 660993 | G → A | At1g10890 | Promoter | – |
| 3980566 | 0.97 | 980566 | G → A | At1g11700 | CDS | G → R |
| 4045453 | 0.90 | 1045453 | G → A | – | – | – |
| 4112858 | 1.00 | 1112858 | G → A | At1g12060 | CDS | E → K |
| 4198211 | 0.95 | 1198211 | G → A | At1g12260 | Intron | – |
| 4241496 | 0.93 | 1241496 | G → A | At1g12360 | CDS | – |
| 4308352 | 0.98 | 1308352 | G → A | At1g12540 | CDS | – |
| 4464407 | 0.92 | 1464407 | G → A | At1g12990 | CDS | R → H |
| 4779263 | 0.89 | 1779263 | G → A | At1g13860 | CDS | L → F |
| 4919214 | 0.91 | 1919214 | G → A | At1g14310 | CDS | R → Q |
| 4949064 | 0.93 | 1949064 | G → A | At1g14370 | Intron | – |
| 4982999 | 0.93 | 1982999 | G → A | At1g14460 | CDS | – |

[a]Position of the nucleotide mutated in the reference genome. [b]Distance to selected position. [c]ref: reference. alt: alternative.

**Table S5.** Candidate mutations identified in *sea4-2* by Easymap

| Position[a] | Allele Frequency | DTP[b] | Nucleotide (ref → alt)[c] | Gene | Gene element | Amino acid (ref → alt)[c] |
|---|---|---|---|---|---|---|
| 1615441 | 0.87 | – 1384559 | C → T | – | – | – |
| 1705363 | 0.93 | – 1294637 | C → T | At1g05670 | CDS | A → T |
| 1905773 | 0.90 | – 1094227 | C → T | At1g06220 | CDS | P → L |
| 2027916 | 0.86 | – 972084 | C → T | At1g06590 | CDS | G → E |
| 2538673 | 0.84 | – 461327 | C → T | – | – | – |
| 2797435 | 0.93 | – 202565 | C → T | At1g08750 | CDS | L → F |
| 2928659 | 0.94 | – 71341 | C → T | At1g09080 | CDS | – |
| 3055323 | 0.88 | 55323 | C → T | At1g09470 | Intron | – |
| 3101327 | 0.91 | 101327 | C → T | At1g09575 | Intron | – |
| 3121169 | 0.96 | 121169 | C → T | At1g09640 | CDS | P → L |
| 3240677 | 0.91 | 240677 | C → T | At1g09940 | Intron | – |
| 3407630 | 0.93 | 407630 | C → T | At1g10320 | Intron | – |
| 3520858 | 0.90 | 520858 | C → T | At1g10586 | Intron | – |
| 3732394 | 0.93 | 732394 | C → T | – | – | – |
| 3843073 | 0.91 | 843073 | C → T | At1g11340 | CDS | V → I |
| 4088965 | 0.96 | 1088965 | C → T | At1g12000 | CDS | G → D |
| 4240281 | 0.87 | 1240281 | C → T | At1g12360 | CDS | S → L |
| 4510248 | 0.93 | 1510248 | C → T | At1g13130 | Intron | – |
| 4515482 | 0.90 | 1515482 | C → T | – | – | – |
| 4634835 | 0.92 | 1634835 | C → T | At1g13410 | Intron | – |
| 4895717 | 0.87 | 1895717 | C → T | At1g14230 | Intron | – |
| 4924756 | 0.89 | 1924756 | C → T | At1g14330 | Intron | – |

[a]Position of the nucleotide mutated in the reference genome. [b]Distance to selected position. [c]ref: reference. alt: alternative.

**Table S6.** Retention of the 9th intron of *KEU* in the RNA-seq assay

| Sample | Number of reads | | Total reads | |
|---|---|---|---|---|
| | with retention | without retention | with retention (%) | without retention (%) |
| L*er* replicate #1 | 0 | 172 | 2.32 | 97.68 |
| L*er* replicate #2 | 7 | 212 | | |
| L*er* replicate #3 | 6 | 164 | | |
| *sea4-1* replicate #1 | 899 | 13 | 99.17 | 0.83 |
| *sea4-1* replicate #2 | 1101 | 4 | | |
| *sea4-1* replicate #3 | 733 | 6 | | |
| *sea4-2* replicate #1 | 15 | 177 | 2.40 | 97.60 |
| *sea4-2* replicate #2 | 0 | 263 | | |
| *sea4-2* replicate #3 | 2 | 250 | | |

**Table S7.** Expression of *KEU* in the RNA-seq assay

| Sample | *KEU* reads | RNA-seq reads | SC[a] | Mean SC | Relative mean SC |
|---|---|---|---|---|---|
| L*er* replicate #1 | 1354 | 57171674 | 23.67 | 24.98 | 1.00 |
| L*er* replicate #2 | 1536 | 62084344 | 24.73 | | |
| L*er* replicate #3 | 1456 | 54922570 | 26.52 | | |
| *sea4-1* replicate #1 | 7897 | 58983680 | 133.85 | 127.36 | 5.10 |
| *sea4-1* replicate #2 | 9319 | 70165558 | 132.75 | | |
| *sea4-1* replicate #3 | 6328 | 54794912 | 115.47 | | |
| *sea4-2* replicate #1 | 2049 | 5450562 | 37.60 | 38.76 | 1.55 |
| *sea4-2* replicate #2 | 2337 | 60379150 | 38.69 | | |
| *sea4-2* replicate #3 | 2328 | 58217096 | 40.00 | | |

[a]Standardized coverage (SC) represents the division of *KEU* reads by the total RNA-seq reads for each sample (measured in millions).

**Table S8.** Predicted effects of the S57L mutations on the stability and dynamics of the KEU protein

| Predictor | $\Delta\Delta G$ (kcal·mol$^{-1}$) | $\Delta\Delta S_{Vib}$ (kcal·mol$^{-1}$·K$^{-1}$) |
|---|---|---|
| DynaMut | 0.648 | – 0.303 |
| mCSM | – 0.194 | – |
| SDM | 1.310 | – |
| DUET | 0.463 | – |
| DynaMut2 | – 0.380 | – |

Values indicate predicted differences in the unfolding Gibbs free energy ($\Delta\Delta G$) and vibrational entropy energy ($\Delta\Delta S_{Vib}$) between wild-type and mutant proteins. Positive and negative $\Delta\Delta G$ values indicate conformational stabilization and destabilization, respectively. Negative $\Delta\Delta S_{Vib}$ values indicate loss of flexibility.

**Table S9.** Phenotypic variability in the progeny of selfed *sea4* plants

| | | Ler | | | *sea4-1* | | | | *sea4-2* | | | |
|---|---|---|---|---|---|---|---|---|---|---|---|---|
| | | NGS | SP | PNL | NGS | PEPSL | SPMSL | PSL | NGS | PEPSL | SPMSL | PSL |
| NGS[a] | NGS[b] | 6 | | | 73 | | | | 1 | | | |
| | SPEC[b] | | 1 | | | 1 | | | | | | |
| | SEC[b] | | | | | | 1 | | | | | |
| | PTL[b] | | | | | | | | | | | |
| SUC[a] | NGS[b] | | | | | | | | | | | |
| | SPEC[b] | | | | | 2 | | | | | | |
| | SEC[b] | | | | | 3 | 3 | | | | | |
| | PTL[b] | | | 1 | | | 1 | 2 | | | | |
| SDC[a] | NGS[b] | | | | | | | | | | | |
| | SPEC[b] | | 2 | | | | | | | | | |
| | SEC[b] | | 4 | | | 6 | 1 | | | | | |
| | PTL[b] | | 3 | 10 | | | 6 | 5 | | 1 | | 2 |
| SCEC[a] | NGS[b] | | | | | | | | | | | |
| | SPEC[b] | | 1 | | | | 1 | | | | | |
| | SEC[b] | | 12 | 1 | | 11 | | 1 | | 1 | 1 | |
| | PTL[b] | | 7 | 174 | | 4 | 5 | 130 | | 16 | 5 | 213 |

[a,b]Phenotypes observed [a]7 das and [b]14 das. All the remaining data were collected 21 das. NGS: non-germinated seeds. SUC: seedlings with unexpanded cotyledons. SDC: seedlings with defined cotyledons. SCEC: seedlings with completely expanded cotyledons. SPEC: seedlings with partially expanded cotyledons. SEC: seedlings with expanded cotyledons. PTL: plants with true leaves. SP: small plants. PNL: plants with normal leaves. PEPSL: plants with poorly expanded and prematurely senescent leaves. SPMSL: stunted plants with mildly serrated leaves. PSL: plants with serrated leaves.

**Table S10.** Venation pattern characteristics of *sea4* cotyledons and leaves

| | | L*er* | *sea4-1* | *sea4-2* |
|---|---|---|---|---|
| Leaf area ($mm^2$) | Cotyledon | 8.62 ± 1.88 | 7.40 ± 1.32 | 6.93 ± 0.89* |
| | 1st node leaf | 33.51 ± 10.25 | 6.30 ± 4.80** | 9.78 ± 3.70** |
| | 3rd node leaf | 52.53 ± 17.53 | 12.85 ± 4.48** | 12.60 ± 6.09** |
| Vein length/area ($mm/mm^2$) | Cotyledon | 1.56 ± 0.15 | 1.56 ± 0.12 | 1.72 ± 0.16* |
| | 1st node leaf | 2.71 ± 0.44 | 3.58 ± 0.87** | 2.93 ± 0.49 |
| | 3rd node leaf | 3.40 ± 0.41 | 3.67 ± 0.47 | 4.43 ± 0.78** |
| Bifurcations/area ($n/mm^2$) | Cotyledon | 0.76 ± 0.15 | 0.75 ± 0.15 | 1.13 ± 0.29** |
| | 1st node leaf | 3.84 ± 1.34 | 7.37 ± 3.81** | 5.07 ± 1.78 |
| | 3rd node leaf | 5.83 ± 1.55 | 7.34 ± 2.09 | 11.55 ± 3.79** |
| Terminal veins /area ($n/mm^2$) | Cotyledon | 0.11 ± 0.12 | 0.22 ± 0.14* | 0.21 ± 0.19 |
| | 1st node leaf | 1.16 ± 0.25 | 3.16 ± 1.60** | 2.06 ± 0.72** |
| | 3rd node leaf | 1.41 ± 0.26 | 2.48 ± 0.64** | 2.96 ± 0.57** |
| Leaf circularity | Cotyledon | 1.21 ± 0.12 | 1.16 ± 0.09 | 1.21 ± 0.09 |
| | 1st node leaf | 1.12 ± 0.09 | 1.24 ± 0.14* | 1.32 ± 0.21** |
| | 3rd node leaf | 1.22 ± 0.07 | 1.26 ± 0.13 | 1.40 ± 0.13** |

Values are mean ± standard deviation. Asterisks indicate a statistically significant difference from L*er* in a Student's *t* test (*$P < 0.05$; **$P < 0.01$).

**Table S11.** Nuclear DNA ploidy distribution (%) in the first pair of leaves of the *sea4* mutants

| Ploidy level | L*er* | *den5-1* | *sea4-1* | *sea4-2* |
|---|---|---|---|---|
| 2C | 23.91 ± 1.88 | 25.89 ± 1.47 | 19.43 ± 0.40* | 18.62 ± 1.01* |
| 4C | 31.06 ± 0.60 | 32.67 ± 1.16 | 26.82 ± 0.89** | 28.29 ± 0.88* |
| 8C | 30.39 ± 1.56 | 20.99 ± 0.62** | 23.22 ± 1.14** | 26.26 ± 2.27 |
| 16C | 11.97 ± 1.18 | 13.40 ± 1.34 | 18.68 ± 0.23** | 17.94 ± 1.09** |
| 32C | 2.21 ± 0.74 | 5.71 ± 0.58** | 8.62 ± 0.51** | 7.15 ± 2.60* |
| 64C | 0.39 ± 0.21 | 1.15 ± 0.56 | 2.69 ± 0.54** | 1.43 ± 0.84 |
| 128C | 0.07 ± 0.07 | 0.19 ± 0.02* | 0.55 ± 0.16** | 0.31 ± 0.21 |

Values are mean ± standard deviation. Asterisks indicate a statistically significant difference from L*er* in a Student's *t* test (*$P < 0.05$; **$P < 0.01$).
